# Mechanistic insights on metabolic dysfunction in PTSD: Role of glucocorticoid receptor sensitivity and energy deficit

**DOI:** 10.1101/492827

**Authors:** Pramod R. Somvanshi, Synthia H. Mellon, Janine D. Flory, Duna Abu-Amara, The PTSD Systems Biology Consortium, Owen M. Wolkowitz, Rachel Yehuda, Marti Jett, Charles Marmar, Francis J. Doyle, Leroy Hood, Kai Wang, Inyoul Lee, Rasha Hammamieh, Aarti Gautam, Bernie J. Daigle, Ruoting Yang

**Affiliations:** Harvard John Paulson School of Engineering and Applied Sciences, Harvard University, Cambridge, MA; Department of Obstetrics, Gynecology & Reproductive Sciences, University of California, San Francisco; Department of Psychiatry, James J. Peters VA Medical Center, Bronx, NY; Department of Psychiatry, Icahn School of Medicine at Mount Sinai, NY; Department of Psychiatry, New York Langone Medical School, New York, NY; Department of Psychiatry, University of California, San Francisco; Integrative Systems Biology, US Army Medical Research and Materiel Command, USACEHR, Fort Detrick, Frederick, MD; Institute for Systems Biology, Seattle, WA; Department of Biological Sciences and Computer Science, The University of Memphis, Memphis, TN; Advanced Biomedical Computing Center, Frederick National Laboratory for Cancer Research, Frederick, MD

**Keywords:** GR sensitivity, HPA axis, systems biology, PTSD

## Abstract

PTSD is associated with metabolic comorbidities; however it is not clear how the neuroendocrine disturbances affect metabolism. To analyze this we employed a systems biological approach using an integrated mathematical model of metabolism, HPA axis and inflammation. We combined the metabolomics, neuroendocrine, clinical lab and cytokine data from combat-exposed veterans with and without PTSD, to characterize the differences in regulatory effects. We used the pattern of fold change in metabolites representing pathway level differences as reference for metabolic control analysis (MCA) using the model. MCA revealed parameters constituting the HPA axis, inflammation and GPCR pathway that yielded metabolic dysfunction consistent with PTSD. To support this, we performed causal analysis between regulatory components and the significantly different metabolites in our sample. Causal inference revealed that the changes in glucocorticoid receptor sensitivity were mechanistically associated with metabolic dysfunction and the effects were jointly mediated by insulin resistance, inflammation, oxidative stress and energy deficit.

---

Post-traumatic stress disorder (PTSD) is defined by a complex set of criteria including intrusive reminders, fear memories, emotional distress, hypervigilance and exaggerated startle responses that develop and persist after an exposure to trauma ^1^. Multiple physiological systems at the neuronal, metabolic, inflammatory, genomic and epigenomic levels are known to be affected in PTSD ^2^,^3^. Previous studies from our PTSD Systems Biology consortium reported associations between PTSD and insulin resistance ^4^, inflammation ^5^, reduced mitochondrial copy number ^6^, and lower methylation of the glucocorticoid receptor (NR3C1) gene ^7^ in the same cohorts assessed in the present study. The data from animal models and humans with PTSD reveal association between inflammation and metabolic syndrome ^8^. Analysis of metabolomics have revealed metabolic changes consistent with dysregulation in mitochondrial functioning in PTSD ^9^. Another study reported differences in the mitochondrial DNA (SNPs) located in NADH dehydrogenase and ATP synthase genes in PTSD ^10^. Variants of mitochondrial genes and dysregulation of their associated networks have been reported in the postmortem brains of patients with PTSD ^11^.

One of the major physiological regulatory axes, Hypothalamic-Pituitary-Adrenal (HPA) axis is implicated in the pathogenesis of PTSD and associated metabolic disorders ^12^,^13^. The HPA axis relays stress signals from the brain to peripheral parts through the release of glucocorticoids (GCs) and catecholamine hormones. GCs orchestrate the activities of several physiological functions through glucocorticoid receptor (GRs) signaling. Therefore, the feedback sensitivity of GC signaling among other factors can influence downstream effects of stress exposure. GR signaling is regulated at transcriptional and epigenetic levels and is dysregulated in HPA axis associated disorders ^14^. Recent studies have highlighted the association of genetic polymorphisms of the GC receptor gene with metabolic disorders and PTSD ^15^. The role of GCs and GC receptor functioning in individuals with PTSD has been characterized ^16^; however, the underlying mechanisms by which the abnormalities in GR signaling contribute to psychiatric and metabolic disorders is not well delineated. Therefore, in light of the complexity of GC’s downstream regulatory effects, a systems level analysis is required to understand dysregulation of the HPA axis and its metabolic consequences.

In view of these findings, we investigated whether the co-occurrence of the multiple metabolic abnormalities are independent or arise from a common underlying regulatory defect and its relative independence with respect to neuro-endocrine changes observed in PTSD. To address this question, we used an integrative systems biological approach to identify regulatory connections in metabolic dysfunction in PTSD. The overall analysis is divided into two sections, namely model-based inferences: hypothesis generation using metabolic control analysis (MCA); and data-based verification of the hypothesis using causal inference ^17^. The pipeline used for the current analysis is depicted in Appendix I-Figure S1.

Initially, we performed statistical analyses on the data obtained from 83 combat-exposed veterans who develop PTSD and 82 combat-exposed veterans without PTSD, to identify the significantly different metabolites, inflammatory and neuroendocrine features in their plasma samples. The demographics of our sample are reported in Appendix I-Table S1. We used the pattern of change in significantly different metabolic pathway components as the reference for MCA. These pathways were represented by 12 metabolites, namely glucose, pyruvate, lactate, citrate, alanine, glutamine, long chain fatty acids, triglycerides, carnitines, arginine, ornithine and insulin (Table 1: subsections represent pathways). We refer the pattern of change in these 12 metabolites as metabolic dysfunction (MD) signature. We used ordinary differential equations (ODEs) based mathematical models from the literature to integrate metabolism with HPA axis activity and inflammation to identify the putative mechanisms underlying this metabolic signature in combat-related PTSD. The details of the model development are given in Appendix-II. Using this model, MCA was conducted to determine potential perturbations in the model that could yield consistent metabolic differences as observed in our PTSD sample. The parameters associated with these perturbations were used to generate hypotheses on the processes that could potentially be affected in PTSD. From the MCA, we determined the parameters representing dysregulation in HPA axis, GR signaling, G-protein coupled receptor (GPCR) signaling and inflammatory pathways could yield the MD signature as observed in our PTSD subjects. Among these mechanisms, systemic GR sensitivity of HPA-immune axis showed higher control on the metabolic differences. Therefore, we focus our further analysis on the effects of GR sensitivity on metabolism.

**Table 1:** The features with statistically significant difference in controls and PTSD subjects with p<=0.05 and q<=0.01. Age & BMI adjusted p and q values are reported. (*) sign in suffix indicate additional features (with 0.05< p <= 0.1) that were included in analysis due to their association with significantly different pathways.

| Metabolites | p.values | q.values | Adjusted p.values | Adjusted q.values | median (FC) | Cohen's d |
| --- | --- | --- | --- | --- | --- | --- |
| **Glycolysis** |  |  |  |  |  |  |
| Lactate | 2.36E-07 | 3.42E-05 | 1.13E-06 | 1.73E-04 | 1.301 | -0.867 |
| Pyruvate | 0.001 | 0.024 | 0.014 | 0.104 | 1.308 | -0.453 |
| Citrate* | 0.039 | 0.138 | 0.049 | 0.188 | 0.949 | 0.346 |
| **Amino acids** |  |  |  |  |  |  |
| Alanine | 0.015 | 0.076 | 0.038 | 0.175 | 1.123 | -0.371 |
| Glutamine | 0.021 | 0.099 | 0.021 | 0.137 | 0.969 | 0.423 |
| Tyrosine | 0.005 | 0.050 | 0.007 | 0.068 | 1.074 | -0.483 |
| Isoleucine* | 0.044 | 0.138 | 0.505 | 0.511 | 1.048 | -0.168 |
| leucine* | 0.090 | 0.184 | 0.281 | 0.400 | 1.034 | -0.236 |
| Valine* | 0.086 | 0.182 | 0.557 | 0.526 | 1.039 | -0.152 |
| **Urea cycle** |  |  |  |  |  |  |
| Arginine* | 0.067 | 0.164 | 0.053 | 0.196 | 0.973 | 0.386 |
| Ornithine | 0.011 | 0.061 | 0.046 | 0.188 | 1.097 | -0.331 |
| **Long chain fatty acids** |  |  |  |  |  |  |
| 10-Nonadecenoate 19:1 (ω-9) | 0.004 | 0.047 | 0.001 | 0.030 | 0.775 | 0.460 |
| 10-Undecenoate 11:1 (ω-1) | 0.006 | 0.054 | 0.033 | 0.165 | 0.821 | 0.371 |
| 17-Methylstearate | 0.001 | 0.026 | 0.002 | 0.030 | 0.881 | 0.441 |
| 2-Hydroxypalmitate | 0.008 | 0.056 | 0.010 | 0.089 | 0.933 | 0.407 |
| Nonadecanoate 19:0 | 3.02E-05 | 0.002 | 2.17E-04 | 0.016 | 0.815 | 0.595 |
| Arachidonate 20:4 (ω-6) | 0.010 | 0.061 | 0.011 | 0.094 | 0.815 | 0.371 |
| Stearate 18:0 | 0.007 | 0.056 | 0.005 | 0.054 | 0.911 | 0.376 |
| **Essential fatty acids** |  |  |  |  |  |  |
| Dihomolinoleate 20:2 (ω-6) | 0.008 | 0.056 | 0.002 | 0.030 | 0.806 | 0.429 |
| Dihomolinolenate 20:3 (ω-3 or ω-6) | 1.84E-04 | 0.009 | 4.30E-04 | 0.016 | 0.785 | 0.504 |
| Docosahexaenoate (DHA) 22:6 (ω-3) | 0.001 | 0.024 | 0.003 | 0.041 | 0.812 | 0.479 |
| Docosapentaenoate (DPA) 22:5 (ω-3) | 0.001 | 0.024 | 0.001 | 0.023 | 0.730 | 0.480 |
| Eicosapentaenoate (EPA) 20:5 (ω-3) | 0.006 | 0.055 | 0.022 | 0.142 | 0.821 | 0.323 |
| Eicosenoate 20:1 | 0.002 | 0.026 | 0.002 | 0.030 | 0.801 | 0.472 |
| Linolenate 18:3 (ω-3 or ω-6) | 0.002 | 0.027 | 0.002 | 0.034 | 0.808 | 0.425 |
| **Carnitines** |  |  |  |  |  |  |
| Decanoylcarnitine | 0.011 | 0.061 | 0.031 | 0.164 | 1.234 | -0.328 |
| Octanoylcarnitine | 0.021 | 0.099 | 0.030 | 0.164 | 1.093 | -0.343 |
| Palmitoylcarnitine* | 0.044 | 0.138 | 0.148 | 0.337 | 1.103 | -0.268 |
| **Energy deficit and oxidative stress** |  |  |  |  |  |  |
| Hypoxanthine | 2.38E-04 | 0.009 | 0.002 | 0.030 | 1.354 | -0.557 |
| 5-oxoproline | 0.002 | 0.027 | 0.000 | 0.016 | 1.104 | -0.522 |
| Gamma glutamyltyrosine | 0.008 | 0.056 | 0.075 | 0.235 | 1.102 | -0.331 |
| Sphingosine 1 phosphate | 0.015 | 0.076 | 0.062 | 0.207 | 1.128 | -0.327 |
| Stearoyl sphingomyelin | 0.006 | 0.055 | 0.010 | 0.089 | 1.085 | -0.437 |
| **Other** |  |  |  |  |  |  |
| Threonate | 0.010 | 0.061 | 0.061 | 0.207 | 0.828 | 0.362 |
| Transurocanate | 0.003 | 0.035 | 0.013 | 0.103 | 0.778 | 0.446 |
| Bilirubin EE | 0.018 | 0.089 | 0.029 | 0.164 | 0.854 | 0.389 |
| Glycerate* | 0.028 | 0.111 | 0.047 | 0.188 | 0.908 | 0.369 |
| 3-Hydroxybutyrate (BHBA) | 0.008 | 0.056 | 0.061 | 0.207 | 0.804 | 0.287 |
| **Cytokines** |  |  |  |  |  |  |
| IL6 | 2.00E-04 | 0.007 | 0.014 | 0.161 | 1.308 | -0.441 |
| TNFα | 0.005 | 0.031 | 0.003 | 0.097 | 1.077 | -0.498 |
| **Clinical Labs** |  |  |  |  |  |  |
| Insulin | 0.002 | 0.017 | 0.025 | 0.188 | 1.122 | -0.497 |
| Glucose | 0.002 | 0.017 | 0.133 | 0.363 | 1.023 | -0.375 |
| HOMA-IR | 3.96E-04 | 0.007 | 0.008 | 0.132 | 1.146 | -0.553 |
| Triglyceride | 0.027 | 0.067 | 0.425 | 0.659 | 1.055 | -0.274 |
| Total protein | 0.001 | 0.012 | 0.006 | 0.132 | 1.015 | -0.490 |
| Albumin | 0.015 | 0.048 | 0.062 | 0.277 | 1.011 | -0.286 |
| Alkaline phosphatase | 0.006 | 0.031 | 0.084 | 0.304 | 1.037 | -0.375 |
| GGT | 3.70E-04 | 0.007 | 0.010 | 0.132 | 1.131 | -0.513 |
| hs-CRP | 0.016 | 0.048 | 0.033 | 0.195 | 1.298 | -0.455 |
| Potassium (K) | 0.006 | 0.031 | 0.046 | 0.227 | 1.025 | -0.391 |
| Chlorine (Cl) | 0.015 | 0.048 | 0.021 | 0.178 | 1.001 | -0.424 |
| White blood cells (WBC) | 0.009 | 0.036 | 0.016 | 0.162 | 1.049 | -0.444 |
| Red blood cells (RBC) | 0.011 | 0.041 | 0.064 | 0.277 | 1.018 | -0.371 |
| Platelets | 0.014 | 0.048 | 0.109 | 0.341 | 1.015 | -0.327 |
| MPV | 0.030 | 0.068 | 0.035 | 0.195 | 1.009 | -0.377 |
| **Neuro-endocrine features** |  |  |  |  |  |  |
| ACTH | 0.036 | 0.074 | 0.219 | 0.467 | 1.069 | -0.281 |
| Cortisol suppression | 0.005 | 0.031 | 0.022 | 0.178 | 1.021 | -0.388 |
| Urinary epinephrine | 0.050 | 0.092 | 0.099 | 0.331 | 1.226 | -0.285 |
| Cortisol* | 0.069 | 0.114 | 0.054 | 0.256 | 1.056 | -0.290 |
| IC50-Dex* | 0.082 | 0.123 | 0.086 | 0.304 | 0.913 | 0.284 |

To ascertain the MCA-based inferences on the underlying mechanisms for the observed metabolic differences, we performed correlational analysis and causal inference ^18^ using our data. Since our data comes from an observational study, we used covariate balancing propensity scores (CBPS) for weighting in estimation of average causal effects ^19^ and performed sensitivity analysis to determine the extent of violation of the ignorability assumption ^20^. We incorporated the results from neuroendocrine and clinical lab assays for our subjects including cortisol suppression (a measure of GC feedback sensitivity in the HPA axis ^21^), IC_50_ of lymphocyte proliferation in response to dexamethasone (DEX) treatment (concentration of DEX at which 50% of lysozyme activity was inhibited), cortisol, urinary epinephrine, hs-CRP (a marker for inflammation), homeostatic model assessment insulin resistance (HOMA-IR), hypoxanthine (a surrogate for hypoxia and energy deficit ^22^,^23^) and gamma-glutamyl transferase (GGT, a surrogate for oxidative stress ^24^) as the regulatory components, along with significantly different metabolites and cytokines for the analysis. We used eight covariates, namely age, BMI, education, race, ethnicity, current medications (10 categories: anti-depressants, sedatives, anti-convulsants, anti-diabetics, anti-allergic, anti-inflammatory, anti-hypertensive, pain medicine, antacid and statins), smoking and alcohol use, to ensure the covariate balance on the exposure variable.

Furthermore, the estimates for average causal effects were used to inform the causal mediation hypothesis that was tested using natural effects models with inverse probability weighting ^25^. We conducted causal mediation analysis with cortisol suppression (CS) by DEX as an exposure variable, HOMA-IR, hs-CRP, hypoxanthine and GGT as the joint mediators while controlling for covariates ^26^. To obtain CS specific effects we also controlled for urinary catecholamine to account for the effects of epinephrine on the β-adrenergic pathway, which could also yield the MD signature as observed by the MCA, and dehydroepiandresterone due to its effect on HPA axis and insulin sensitivity. Our analysis suggests that the mechanistic association of enhanced GC feedback sensitivity with metabolic dysfunction is partly mediated by inflammation, oxidative stress, insulin resistance and energy deficit.

## Results

### Group differences in features between PTSD and controls

To assess the group differences in our data, we performed a nonparametric Mann-Whitney U test followed by adjustments for false discovery rate and estimation of effect sizes given by Cohen’s d. The summary statistics for group differences in metabolites that showed p-value<0.05 and q-value<0.1 are reported in Table 1. We identified 31 metabolites, 2 cytokines, 14 clinical variables and 3 neuro-endocrine variables significantly different (at p-value<0.05 and q-value<0.1) between PTSD and control subjects. The alterations in the metabolite flows in the metabolic pathways are depicted in Figure 1. These findings can be attributable to mitochondrial dysfunction inferred by the disturbance in the inflow-outflow of metabolites (carbohydrates, fatty acids, amino acids) and mitochondrial metabolite processing (TCA cycle, beta-oxidation, urea cycle, and amino acid catabolism).

**Figure 1.**
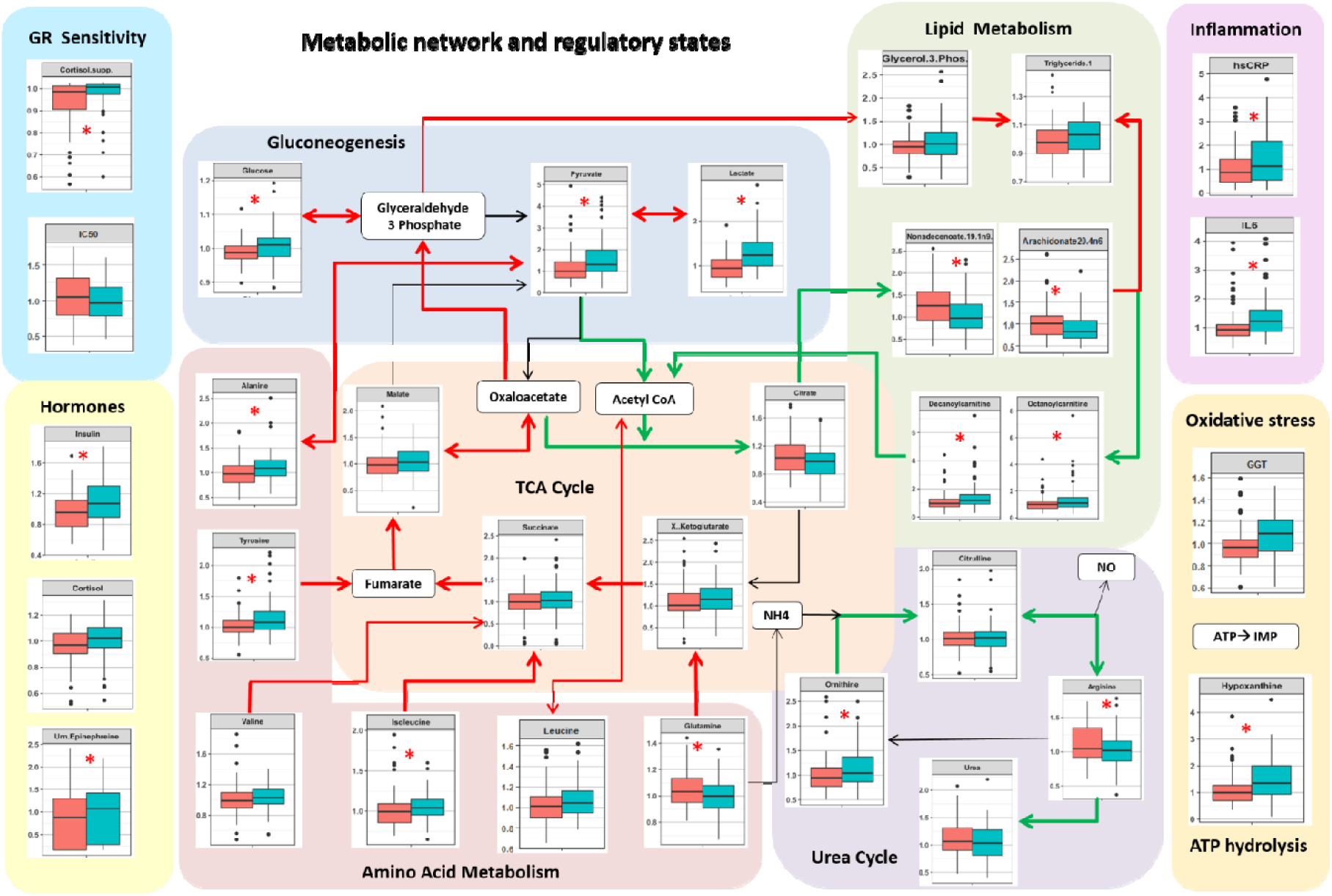
Boxplot representation of metabolite differences in controls and PTSD. Red arrows represent the upregulated fluxes and the green arrows represent down regulated metabolic fluxes. The side panels represent differences in metabolic regulatory hormones, tests for GR sensitivity and markers for inflammation, oxidative stress and ATP hydrolysis. A red star on the box plots represent the statistically significant difference (p<0.05) in PTSD versus controls. Green box = PTSD, red box = controls. It is noted that gluconeogenesis potentially in liver, hypoxic adaptation potentially in muscle, amino acid catabolism, triglyceride synthesis and ATP hydrolysis are upregulated, whereas, urea cycle, lipogenesis and beta-oxidation are down-regulated. The measures of GR sensitivity, inflammation and oxidative stress; and the hormones: insulin, cortisol, urinary epinephrine are also elevated in PTSD.

### MCA-based hypothesis for metabolic dysfunction in PTSD

To analyze the putative mechanisms underlying the observed changes in metabolism and the effects of variation in neuroendocrine and inflammatory response on metabolism, we used a systems level mathematical model composed of hepatic metabolism integrated with mathematical models for the HPA-axis, inflammation and hypoxia signalling from the literature. Using the model, MCA ^27^ was conducted to determine potential perturbations in the model that could yield a consistent MD signature. All the metabolic rate parameters and the parameters for signaling and transcriptional regulatory interactions from HPA-axis, inflammation and transcription pathways were considered for the MCA. Through MCA, we obtained metabolite concentration response coefficients (MCRCs) with respect to the trends in the 12 metabolites. MCRC quantifies the degree of control exerted by the change in a parameter value, on the concentration of a metabolite. We estimated MCRCs for 360 model parameters, and identified 34 rate parameters that could yield a mean cumulative response coefficients of at least 0.1 for 12 metabolites taken together (with MCRC of at least 0.001 for each metabolite), across 50% perturbation in the parameter. Figure 2A shows the mean cumulative control coefficients of parameters with relative contribution towards each metabolite perturbation. Figure 2B shows the relative contribution of each response coefficient to changes in metabolites in MD signature. These parameters, if perturbed, yielded a consistent MD signature, in terms of the direction of change (whether increased or decreased) in PTSD subjects with respect to controls. Among the most influential parameters, increasing systemic glucocorticoid receptor sensitivity (GR sensitivity of HPA axis central negative feedback and sensitivity of cytokine regulation taken together) yielded the highest control coefficient, followed by parameters for perturbations in inflammatory response and GPCR pathway (Figure 2A). The rate parameters and corresponding processes alongwith the mean cumulative MCRCs (MCRC of at least 0.001 for each the 12 metabolites) are summarized in Appendix III (Table S5).

**Figure 2.**
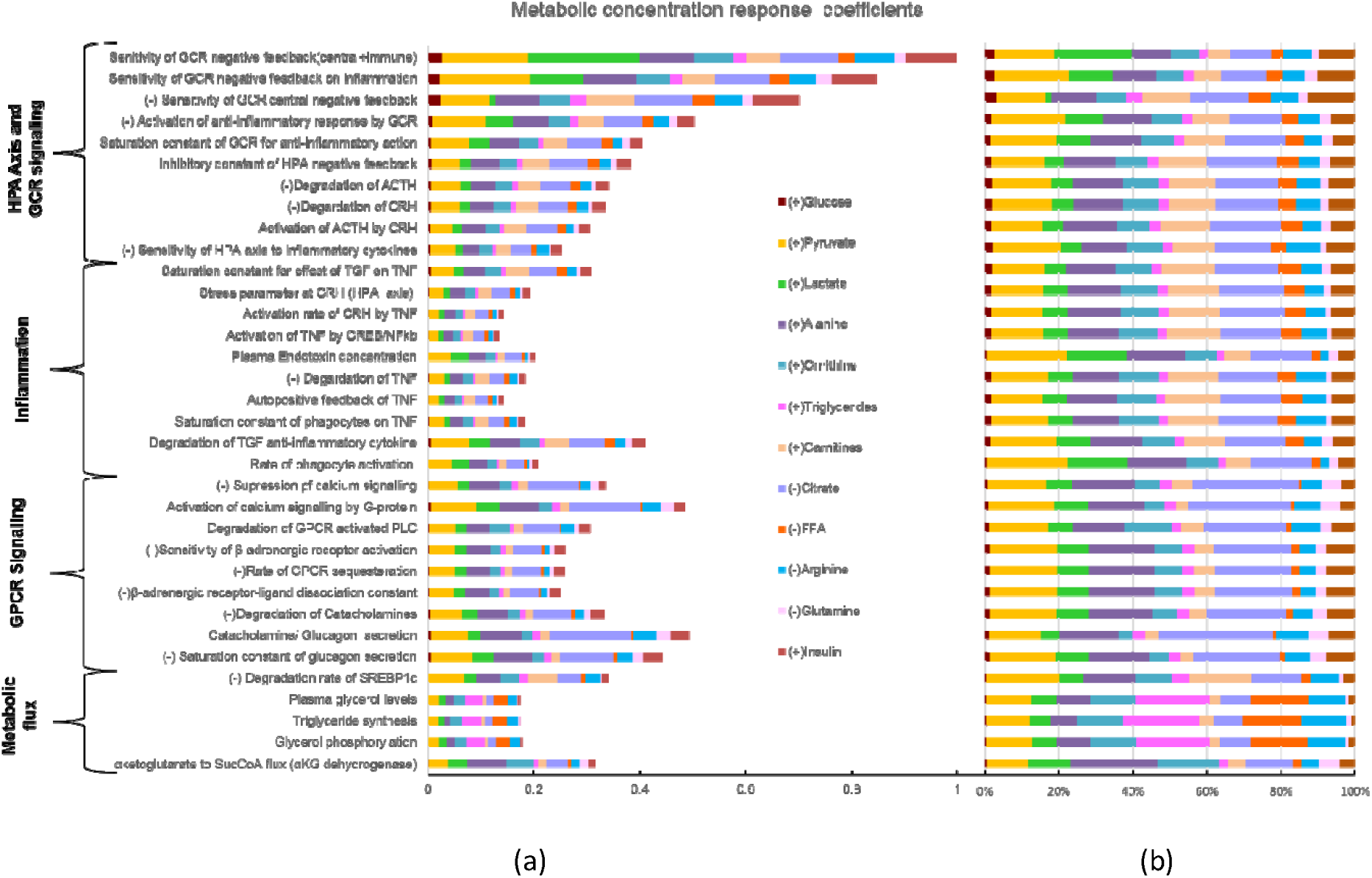
(A) Plot represents the result of MCA: 34 model parameters that elicited the metabolic signature as observed in PTSD subjects, along with their mean cumulative metabolic concentration respose coefficient measured across 12 metabolites (middle of the figure) with at least 0.1 mean cumulative MCRC. The sign (-) in prefix of the parameters represent the direction of change that yielded the MD signature. These parameters belong to the HPA axis and GR signaling, GPCR signaling, inflammation and metabolic fluxes for triglyceride synthesis, plasma lactate and amino acid levels. The color codes the fraction of the control coefficients corresponding to each metabolite. (B) The representation of percent contribution of the control coefficients with respect to individual metabolites per parameter. It was noted that citrate and pyruvate were mostly affected by these parameters contributing to around 40% of the total effect per parameter followed by the effect on amino acid (alanine and glutamine) concentration (∼15%), lipid metabolites (carnitine, fatty acids and triglycerides) concentration (∼15%), urea cycle metabolites (∼15%) and glucose, lactate and insulin together (∼15%).

To probe the mechanisms by which the MD signature was accrued for these 34 parameter perturbations, we further analyzed the corresponding metabolic regulatory states for each parameter perturbation through the model simulations. We recorded the states of regulatory components such as ATP/ADP and NADH/NAD ratios, anabolic and catabolic signaling pathway components, transcriptional regulators and inflammatory pathway as shown in Figure 3. The details on the regulatory states associated with these parameters can be found in Appendix III. On further accounting for the increase in the inflammatory cytokines (IL6 and TNF) and the HPA-axis variables (ACTH and cortisol) along with the 12 metabolic components, parameters constituting GPCR signaling, GCR signaling and inflammation produced a response consistent with MD signature comprising these 16 features. Therefore, from the MCA and the analysis of associated regulatory states, it can be inferred that the steady state perturbation in the parameters for HPA axis and β-adrenergic signaling are accompanied by the combination of changes in metabolic controller ratios (ATP/ADP and NADH/NAD), inflammatory response, insulin resistance and catabolic state associated with an overall energy deficit. Hence, we hypothesize that the trauma induced changes in the systemic glucocorticoid receptor sensitivity are mechanistically associated with insulin resistance, inflammation, oxidative stress, energy deficit and subsequent metabolic dysfunction in PTSD.

**Figure 3:**
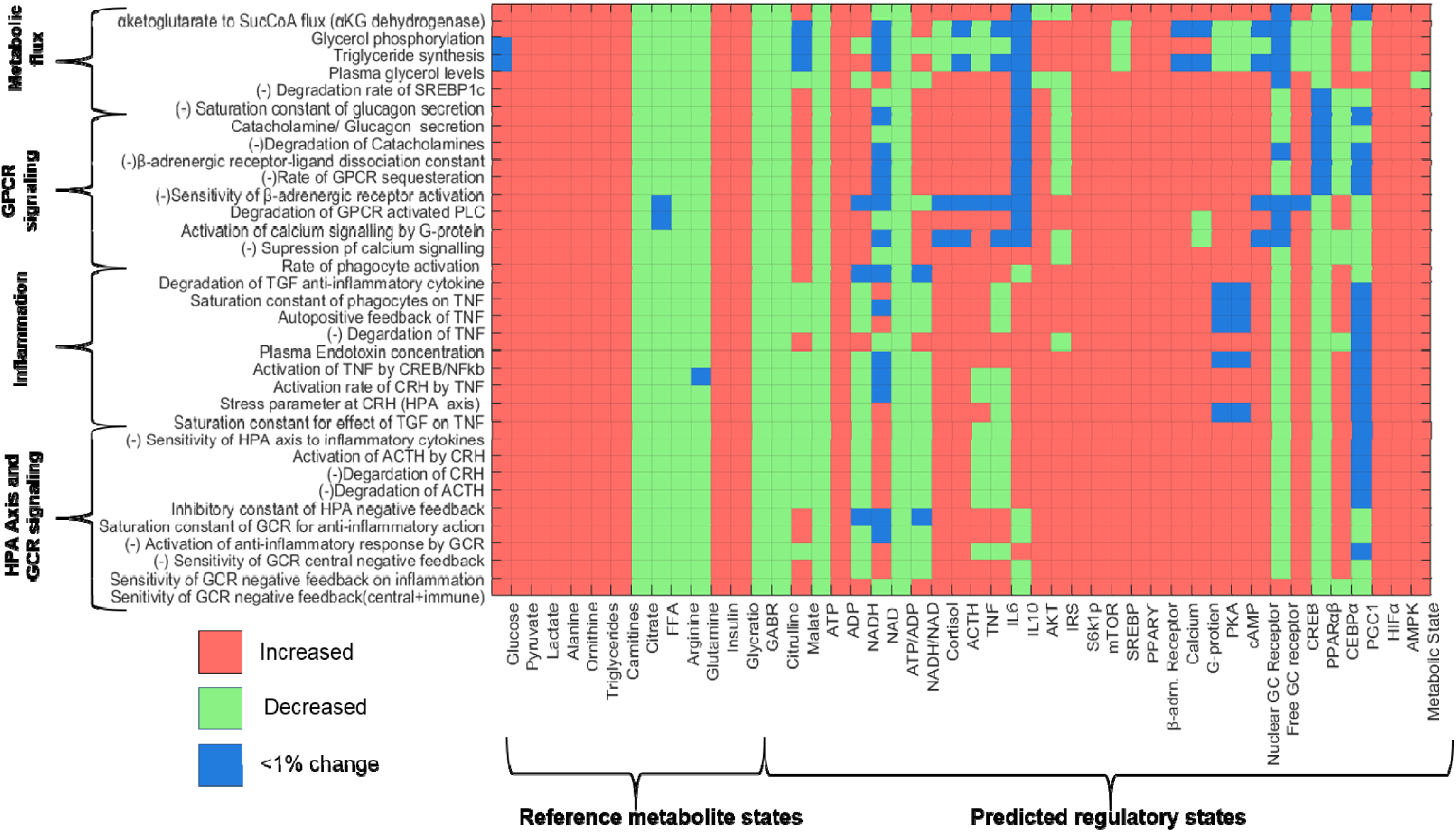
Matrix representation of the changes in the states of metabolite and regulatory component with respect to reference state for perturbation in the 34 parameter, identified through MCA. The states are coded as either increased (red) or decreased (green) and <1% change (blue) per cell corresponding to the parameter-state combination. It can be noted that the trends in reference metabolites (MD signature observed in PTSD) are replicated for the parameters on Y-axis. The predicted states of other regulatory components corresponding to these parameter perturbations are shown appended to reference metabolites. It can be noted that β-adrenergic-GPCR pathway is upregulated for all parameters along with an upregulation of catabolic state. The ATP/ADP ratio is also reduced for all the parameter perturbation along with upregulation of AMPK and HIFα. A net catabolic state can be observed (last column: metabolic state) for all the 34 parameter perturbations.

### Causal Inference

To assess the of model-based hypothesis on the effects of variables associated with pathways inferred from MCA, we performed correlation analysis followed by causal inference between the regulatory components and the metabolites that show statistically significant differences in the two cohorts. Please refer to Appendix IV on correlational analysis for the details on selection of metabolites and the significant correlations between regulatory components and the metabolites (See Figure S5 Table S6). Since we focused on analyzing the effect of GC receptor sensitivity, we tested the causal hypothesis for six regulatory components (dexamethasone cortisol suppression (CS), IC_50_, HOMA-IR, hs-CRP, GGT and hypoxanthine) with respect to 35 features by estimating and testing the sensitivity of average causal effects (ACEs: 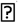) to unmeasured confounding. The average ACEs along with their statistics and sensitivity estimate (τ_1_ and τ_2_) are reported in Table 2. Figure 4 (A-F) shows the statistically significant ACEs for the six regulatory components.

**Table 2:**
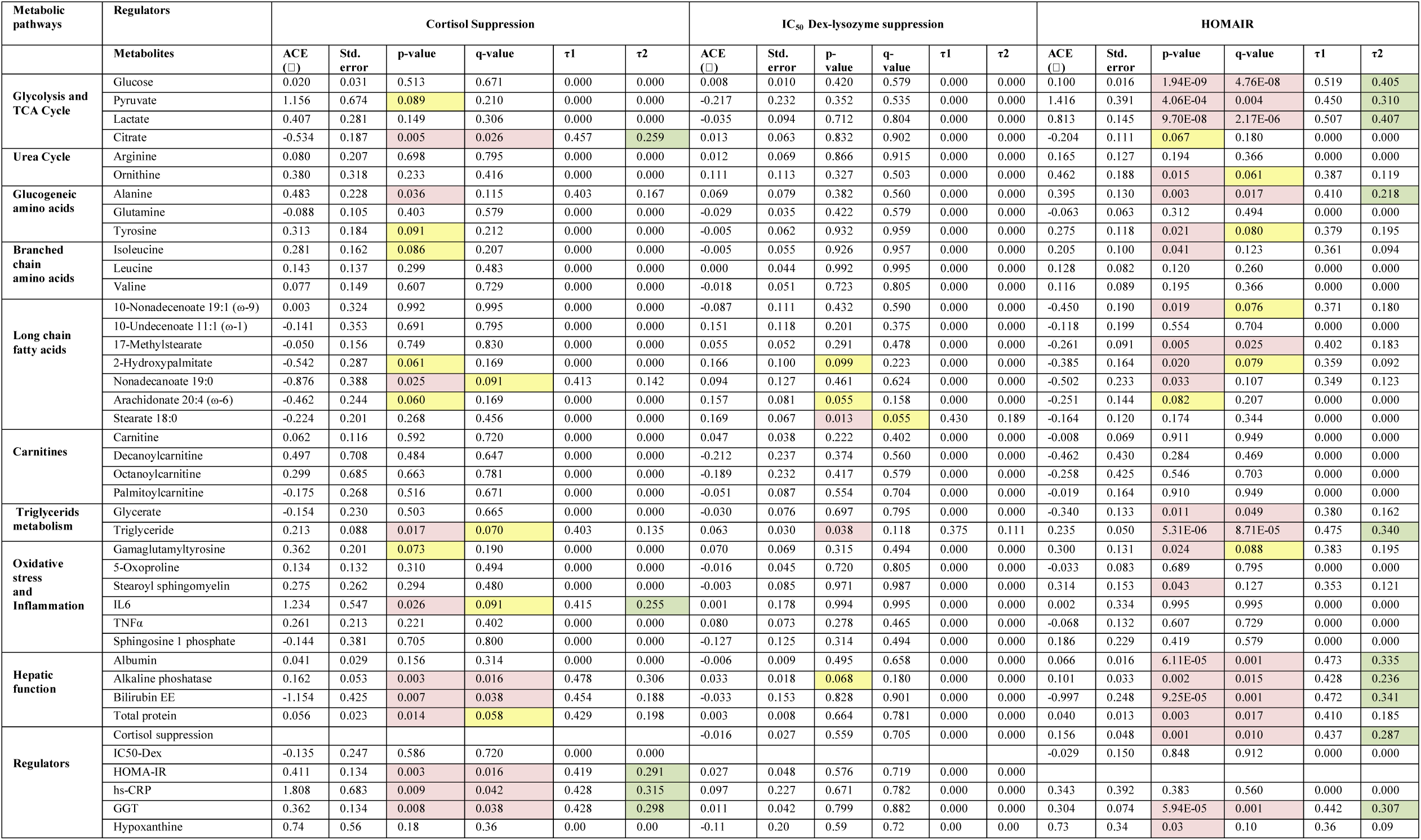
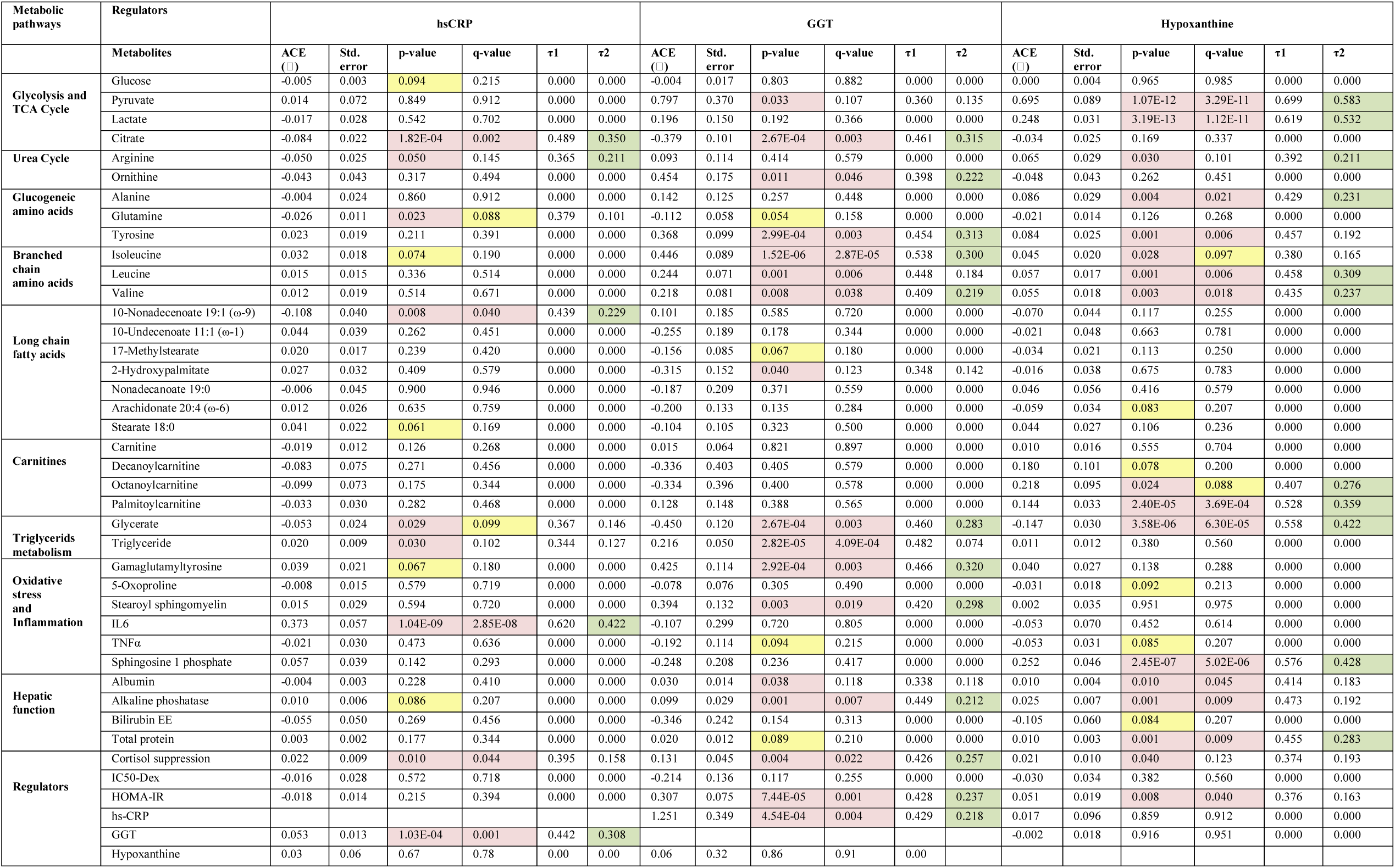
Population level average causal effects. Sensitivity coefficient τ1 representing the coefficient of unobserved confounder at which ACE=0, sensitivity coefficient τ2 representing the coefficient of unobserved confounder at which ACE becomes statistically insignificant (p>0.05). Red, yellow and green shade highlights the p and q values <0.05, between 0.05 and 0.1 and τ2 >0.2, respectively.

**Figure 4:**
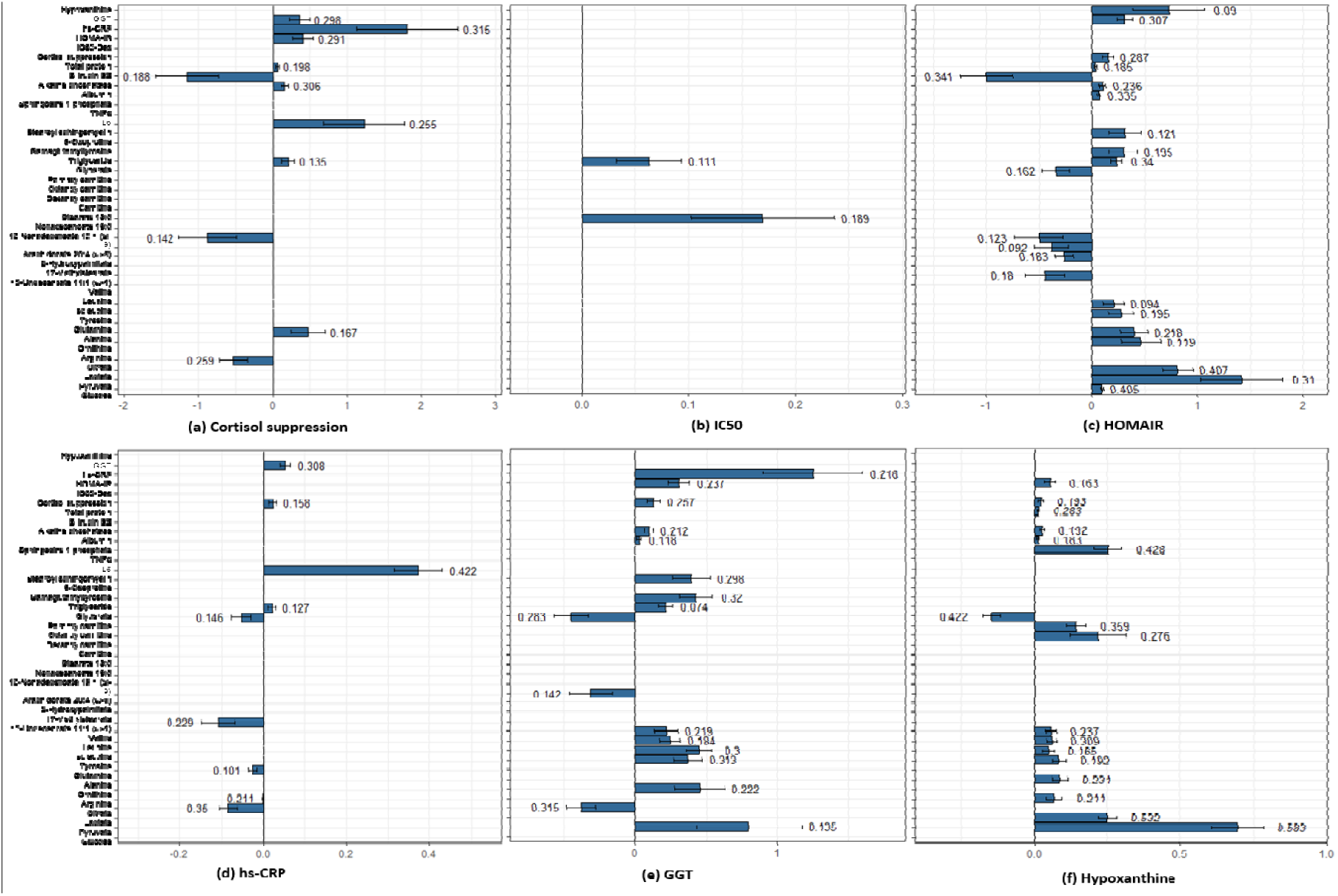
Plots for average causal estimates (ACE) for regulatory components on metabolites. The number on error bar is the sensitivity parameter. (A) Higher CS shows negative effect on citrate, bilirubin and nonadecenoate, whereas positive effect on triglycerides, IL6, alkaline phosphatas, plasma proteins, HOMA-IR, GGT and hs-CRP. (B) IC_50_ is shows a trend with stearate and triglycerides. (C) HOMA-IR shows an identical effect to MD signature for glycolytic metabolites, amino acids, fatty acids, CS, GGT and hypoxanthine along with hepatic function components. (D) hs-CRP shows a negative association with citrate, glutamine, unedecenoate, and glycerate and a positive effect on IL6, CS, triglyceride and GGT. (E) GGT shows a causal effect identical to the MD signature for pyruvate and citrate, ornithine, amino acids, triglycerides, stearoyl sphingomyelin and gammaglutamyltyrosine, hepatic function components, CS, HOMA-IR and hs-CRP. (F) Hypoxanthine corroborates with several features of MD signature: pyruvate, lactate, amino acids, carnitines and HOMA-IR along with glycerate, sphingosine 1 phosphate, hepatic function and CS.

We observed that cortisol suppression showed strong causal association with multiple regulatory components, such as HOMA-IR, hs-CRP and GGT; and these regulatory components were further causally associated with hypoxanthine and several metabolic pathways. This indicated the possibility of mediation of the effects of changes in GC receptor sensitivity on metabolites through these components. To validate this, we next performed causal mediation analysis on the causal structure as shown in Figure 5A. For the sake of simplicity we assume an acyclic graph to identify the causal effects. Due to possible bi-directionality of the effects within the regulatory components, we evaluated joint mediated effects through the mediator complex to satisfy ignorability. The detailed reporting of the summary statistics can be found in Table S7. Figure 5B shows the estimates with 95% confidence interval for natural indirect effect (joint mediated effects) by cortisol suppression or GR sensitivity on the metabolites. Refer to Appendix V and Figure S6 for significant mediated effects, natural direct effect (NDE) and total causal effects.

**Figure 5:**
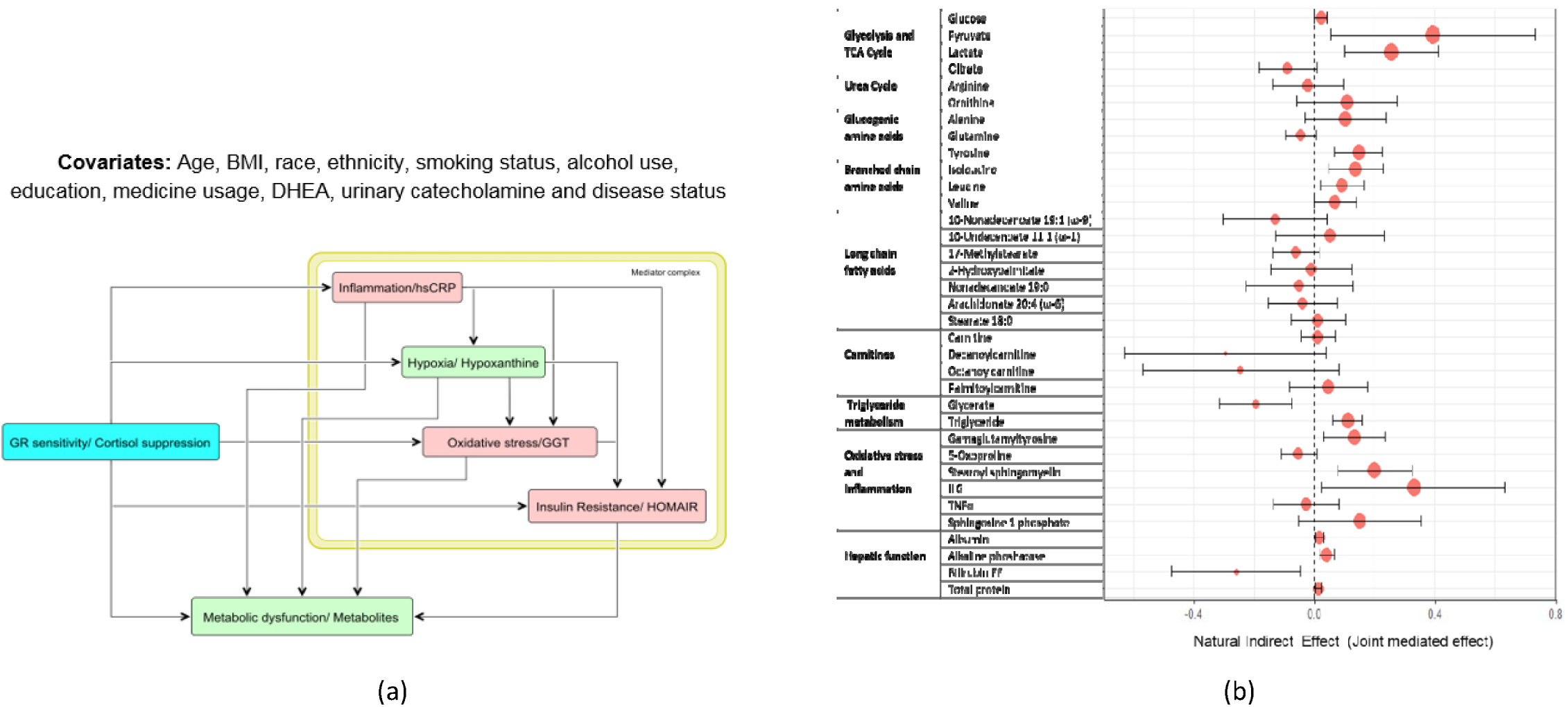
(A) Causal graph used to test for mediation hypothesis of the effect of glucocorticoid sensitivity measured by dexamethasone suppression test (DST). The mediator complex of hs-CRP, hypoxanthine, GGT and HOMA-IR were considered as the joint mediators as informed from the model analysis. (B) Forest plot representation of the natural indirect effects (joint mediated effects) of increased GC feedback sensitivity (measured by cortisol suppression test) on 35 metabolites for the causal hypothesis tested on the entire cohort adjusting for the group effects. The error bar represents 95% confidence intervals of the point estimates of the effects. It is noted that the joint mediated effects on pyruvate, lactate, citrate, gluconeogenic and branched chain amino acids, oxidative stress, inflammation and hepatic function components are statistically significant. Mediated effects on fatty acids and carnitines were insignificant.

### Mechanistic inference from MCA and causal inference

We reconcile the MCA-based findings to mechanistically explain the effect of stress induced changes in HPA axis on metabolism. We then consolidate the explanations with the findings from our data. The regulatory landscape observed in model simulations for the parameter perturbations (Table S5) indicates that changes in metabolism along with increased GR sensitivity and inflammation are mechanistically associated with energy deficit. At the physiological level, how the processes identified by the MCA (GR sensitization, inflammation, insulin resistance and GPCR activation) can result in an energy deficit and the metabolic phenotypes observed in our PTSD sample (Figure 1) are discussed below in light of the causal analysis and supporting literature.

#### 1. Increased insulin resistance

Confronted with exposure to a traumatic event, stress hormones (cortisol and catecholamine) are known to initiate gluconeogenesis to facilitate hepatic glucose production to cope with the energy requirements of an excessive activity in brain and muscles ^28^. These metabolic shifts are achieved by reducing insulin sensitivity and increasing the beta-adrenergic receptor signaling (acute stress) and glucocorticoid receptor signaling (containment of acute stress response) to enhance catabolism for glucose production and utilization. The effect of insulin insensitivity is further compensated by elevation of plasma insulin secretion to restore plasma glucose levels. In our data, this is observed as insulin resistance indicated by increased HOMA-IR (q=0.007) and a significant ACE of cortisol suppression on HOMA-IR (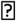 =0.411, p=0.003, q=0.016, τ_2_=0.291). Moreover, in insulin dependent tissues (muscle and adipose) insulin action is required for glucose transport and mitochondrial conversion of pyruvate to acetyl CoA, hence insulin resistance would affect efficient ATP generation. Our data shows a significant association of hypoxanthine with HOMA-IR (ρ=0.292, p=1.44E-4, q=0.001), corroborating plausibility of this mechanism.

#### 2. Increased gluconeogenesis

Induction of gluconeogenesis in liver is associated with increase in pyruvate derived from gluconeogenic precursors such as amino acids (alanine, glycine, cysteine and serine) ^29^. Gluconeogenesis consumes an equivalent of 11 ATPs for driving pyruvate to glucose with a loss of 8 ATPs that could be generated if pyruvate had been processed in mitochondria. Therefore, it is an energy consuming process and its sustained upregulation would lead to an energy deficit state with changes in ATP/ADP and NADH/NAD ratios, thereby affecting all the pathways that depend on adenosine phosphorylation and redox potential for their substrate utilization. Our data shows a significant association of cortisol suppression with glucose (ρ=0.214, p=0.006, q=0.02) and hypoxanthine with glucose (ρ=0.217, p=0.005, q=0.019), corroborating this mechanism.

#### 3. Potential hypoxic adaptation

In the liver, increased diversion of pyruvate to gluconeogenesis can reduce mitochondrial substrate availability resulting in reduced levels citrate (p=0.039) and subsequent alpha-ketoglutarate (αKG). It is known that HIF1α is sensitive to the changes in the level of (αKG), therefore reduced αKG would induce upregulation of HIF1α ^30^. Additionally, the upregulation of pro-inflammatory state is known to induce hypoxia with upregulation of HIF1α ^31^. Although HIF1α would act to sensitize insulin action ^32^, the concomitant inflammation due to GR sensitization (discussed in subsection 9 would maintain insulin resistance and subsequent gluconeogenesis in liver.

In the tissues like muscle that utilize glucose, upregulation of HIFα is known to influence glycolytic enzymes leading to lactate accumulation, which in turn stabilizes HIF1α ^33^. This leads to reduced TCA flux and lower ATP production through oxidative phosphorylation. Therefore in order to compensate for the lower supply of phosphate groups in metabolic reactions, an increased breakdown of ADP to AMP leads to activation of AMP kinase that upregulates glycolysis. These overall may lead to aerobic glycolysis and an energy deficit state. The plausibility of such a mechanism is supported by our data with the significant ACE of hypoxanthine on pyruvate (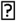 =0.695, p=1.07E-12, q=3.29E-11, τ_2_=0.583) and lactate (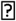 =0.248, p=3.19E-13, q=1.12E-11, τ_2_=0.532). We also observed a significant association of cortisol and cortisol suppression with pyruvate and lactate (See Table S6), and a negative ACE of cortisol suppression on citrate (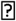 =-0.534, p=0.005, q=0.026, τ_2_=0.259). Moreover, hypoxanthine showed a statistical trend of negative association with plasma phosphate levels (ρ=-0.23, p=0.04) and a positive association with adenosine monophosphate (AMP) (ρ=0.21, p=0.06) in our PTSD subjects.

#### 4. Reduced β-oxidation of fatty acids

During catabolic state, it is crucial to maintain the cellular levels of NADH and ATP, therefore gluconeogenesis is usually associated with fatty acid β-oxidation to maintain the steady supply of NADH and ATP ^34^. The gain of ATP from β-oxidation is relatively high, such as the ATP yield from oxidation of a typical palmitoyl CoA is about 106 ATP accounting for 2 ATPs used in fatty acid activation. Although, the conventional catabolic signaling would upregulate fatty acid β-oxidation, the sustained higher levels of GCs or active GR signaling are known to inhibit β-oxidation in hepatocytes ^35^. This is also indicated in our data from a significant association between plasma cortisol levels and carnitine derivatives (See Table S6). Moreover, upregulation of HIFα (hypoxia) is shown to be associated with reduction in β-oxidation ^36^ limiting carnitine utilization, which is also corroborated in our data by the significant ACE of hypoxanthine on carnitine derivatives (palmitoylcarnitine: 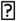 =0.252, p=2.45E-7, q=5.02E-6, τ_2_=0.576; also see Fig 3F). This action would reduce mitochondrial carnitine utilization and further reduce ATP availability from β-oxidation indicating another mechanism for energy deficit. Our PTSD subjects (Table S7) and another study on both animal and humans with PTSD ^37^ show an increased levels of carnitines indicating the possibility of this mechanism.

#### 5. Impaired lipid metabolism

While glucocorticoids induce insulin resistance (IR) in liver and muscle, it is known to sensitize insulin signaling in adipose tissues ^38^. Therefore, hyperinsulinemia due to IR would sensitize adipose for triglyceride synthesis, which is also associated with increased GC activity ^39^. Triglyceride synthesis is an energy intensive process that consumes 6 ATPs per 3 molecules of fatty acids and 1 molecule of glycerol. Therefore, the sustained higher activity of GCs would further add to the effect of lowering ATP availability. This action can be corroborated from our data by significant positive ACE of HOMA-IR (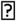 =0.235, p=5.31E-6, q=8.71E-5, τ_2_=0.34) and cortisol suppression (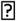 =0.213, p=0.017, q=0.07, τ_2_=0.135) on plasma triglyceride levels.

Fatty acid synthesis is a redox dependent process that consumes ATP and NADH and is regulated by insulin action ^40^. The reduction in mitochondrial citrate (p=0.039) due to increased lactate flux can lead to reduced substrate availability for lipogenesis. Therefore, the lower availability of these substrates and insulin resistance can reduce lipogenesis in non-adipose tissues, whereas high GCs drive fatty acids to triglyceride synthesis in adipose tissues. These dual action can lead to lower levels of circulating fatty acid. This is indicated by significant negative causal association of HOMA-IR and cortisol suppression with fatty acids in our data (See Table 2 and Table S6).

#### 6. Impaired amino acid metabolism

Proteins may be broken down to produce individual amino acids, ready to be converted into usable molecules for gluconeogenesis (or the TCA cycle to produce energy) through anaplerosis. While alanine (p=0.015) is directly converted to pyruvate, glutamine (p=0.021) and tyrosine (p=0.005) enters the TCA cycle at αKG and fumarate nodes, respectively. Stress induced GC activity is known to upregulate proteolysis ^41^. Similarly, our data shows a significant ACE of cortisol suppression on total plasma proteins (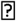 =0.056, p=0.014, q=0.058, τ_2_=0.198) and a trend level effect on alanine and tyrosine. Efficient insulin action is essential for inhibition of proteolysis ^42^; therefore, insulin resistance would enhance the proteolytic effect adding to amino acid pool in the plasma. This is indicated by significant causal association between HOMA-IR and amino acids (see Table 2) in our data. Under tonic hypoxia, HIFα is also known to upregulate glutaminolysis ^43^ in an attempt to restore mitochondrial redox potential. This is also indicated in our sample by negative association between hypoxanthine and glutamine (ρ=-0.159, p=0.041, q=0.098). Moreover, an efficient catabolism of branched chain amino acids (BCCAs) contributes to the pool of NADH and ATP generation in muscle, therefore impaired BCCA catabolism would lead to lower energy supply and increased BCCAs in the circulation as observed by significant associations between BCCAs, GGT and hypoxanthine in our data (See Table 2). Further, amino acids have an ability to influence glucagon secretion and induce gluconeogenesis ^44^ leading to an auto positive feedback effect.

#### 7. Impaired urea cycle and NO production

The processing of amino acids involves deamination generating ammonia that needs to be purged from the body through the urea cycle ^45^. In the urea cycle, formation of carbamoyl phosphate from ammonia and carbon dioxide requires 2 ATPs. The reduced availability of ATP can thus limit formation of carbamoyl phosphate and its subsequent reaction with ornithine, thereby leading to accumulation of ornithine (p=0.01) and reduced levels of citrulline and arginine (trend), as observed in our data. Further, elevated arginase expression is shown to be associated with increased inflammatory cytokines and GC activity ^46^, which could further lead to a lower urea cycle flux, reduced arginine levels and subsequent nitric oxide production. This is also supported by a significant positive association between cortisol suppression and ornithine (ρ=0.223, p=0.004, q=0.015) and a trend of causal association between hs-CRP and arginine (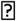 =- 0.05, p=0.05, q=0.145, τ_2_=0.211). The corresponding findings of reduced global arginine bioavailability in the individuals with PTSD were also previously reported in our sample ^47^.

#### 8. Increased β-adrenergic GPCR signaling

The upregulation of β-adrenergic GPCR signaling by catecholaminergic and non-genomic actions of glucocorticoid receptor signaling ^48^ is known to activate cAMP and calcium signaling that influence TCA enzymes and oxidative phosphorylation for production of NADH and ATP, respectively. However, sustained elevation the in cellular calcium signal stimulates higher mitochondrial ROS generation ^49^ due to enhanced electron flow through respiratory chain and subsequent inhibition of complex I & III of Q-cycle. ATP depletion and increase in AMP/ATP ratios is also associated with increase in ROS generation ^50^. In our sample, urinary epinephrine (activator of β-adrenergic signaling) showed a significant correlation with hypoxanthine (ρ=0.17, p=0.029, q=0.07) implying an additional role of enhanced GPCR signaling in energy deficient state.

#### 9. Increased GR Sensitivity and inflammation

Although glucocorticoids are widely known for their anti-inflammatory activity, the pro-inflammatory effects are also documented ^51^. Our model analysis showed that increasing GR sensitivity of central negative feedback of HPA axis and its anti-inflammatory activity, increased the pro-inflammatory response. This is because of the feedback loop that leads to reduction in the gain on GR synthesis due to lower ACTH stimulated cortisol release and subsequent reduction in the anti-inflammatory response (See Figure S7). This results in increased pro-inflammatory milieu that acts to further upregulate HPA axis in an attempt to restore homeostasis, however at the cost of dysregulated immune response ^52^. Our sample revealed a significant and robust positive causal association of cortisol suppression with hs-CRP (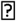 =1.808, p=0.009, q=0.038, τ_2_=0.315) and IL6 (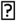 =1.234, p=0.026, q=0.091, τ_2_=0.255); and a trend level reduction in IC_50_ levels (p=0.08). These indicates the plausibility of such a mechanism for increased GR sensitivity of central negative feedback and immune system.

#### 10. Interplay of oxidative stress, inflammation, insulin resistance and energy deficit

Chronic exposure to higher glucocorticoid is associated with oxidative stress ^53^. Oxidative stress is known to inhibit insulin signaling thereby causing insulin resistance. Oxidative stress is also associated with induction of inflammation ^54^ and activation of HIFα. Systemic inflammation is known to induce insulin resistance ^55^ and oxidative stress. Moreover, inflammatory cytokines and ROS are known to activate sphingosine 1 phosphate (S1p) and is implicated in pathophysiology of hypoxia and psychiatric disorders ^56^,^57^. These interactions are also supported by our data with significant ACEs among HOMA-IR, GGT, hs-CRP and hypoxanthine (See Table 2 and Table S6). Corroborating in our data, we also observed the correlates of impaired antioxidant pathway noted by significant differences in gammaglutamyl tyrosine (p=0.008) and oxoproline (p=0.002), and the elevated levels of stearoyl sphingomyelin (p=0.006) that is implicated in oxidative stress, insulin resistance and inflammation ^58^. We also observed significant causal associations of S1p with hypoxanthine (Lr=0.252, q=5.02E-6, τ_2_=0.428), suggesting an association of inflammation and energy deficit. These multiple interaction between comorbidities would constitute a mechanism for positive feedback on mitochondrial dysfunction.

#### 11. Effects of sensitive GR on metabolism are jointly mediated by regulatory complex

Through the model simulations, for a modest increase in GC sensitivity, we observed increase in inflammatory cytokines, plasma insulin and glucose indicating insulin resistance, hypoxic response and impaired NADH/NAD and ATP/ADP ratios indicating energy deficit. These features were collectively responsible for the metabolic phenotype observed in the MD signature. The model-based observations matched well with the interplay of these phenotypes in our data as described in earlier sections. Accordingly, we observed a significant ACE of cortisol suppression on hs-CRP (Lr = 1.394, q=0.067, τ2=0.207), HOMA-IR (Lr = 0.372, q=0.01, τ2=0.265) and GGT (Lr = 0.32, q=0.028, τ2=0.256). Moreover, the causal mediation analysis showed that the effect of increased GR sensitivity on metabolic phenotypes of abnormalities in glycolysis, TCA cycle, amino acid metabolism, triglyceride metabolism, and hepatic function were jointly mediated by these regulatory components along with energy deficit (hypoxanthine) (See Table S7 and Appendix V for detail on significant joint mediated effects).

## Discussion

Our analysis involves a systems approach to identify mechanisms that may cause metabolic dysfunction consistent with group differences in our cohort. We made a novel attempt to integrate earlier observations at multiple levels of the HPA axis, immune system, metabolism and their regulations together in a form of systems model. A model-based analysis reveals how multiple pathways at metabolic, signaling and transcriptional levels interact together to ensure homeostasis and its disruption in disease state. This multilevel analysis provides insight on etiological aspects of physiological response to chronic stress and the resulting metabolic dysfunction. We used statistical analysis, mathematical modeling and causal inference to generate and verify the mechanistic hypothesis.

In our analysis, we have shown how the defects in GR signaling might affect systemic metabolic regulation and subsequent changes in metabolic pathways in PTSD. Based on our overall analysis we infer that the metabolic phenotype of MD observed in PTSD could at least in part be due to the effect of trauma induced glucocorticoid receptor sensitivity that may result in insulin resistance, inflammation, oxidative stress and subsequent energy deficit. The sustained elevation of these effects may induce (i) enhanced gluconeogenesis in liver; (ii) tonic pseudo-hypoxia induced aerobic glycolysis and reduced β-oxidation in muscle and liver; and (iii) enhanced triglyceride synthesis in adipose tissues and liver. These collectively may affects mitochondrial fuel processing leading to an apparent mitochondrial dysfunction. Moreover, we shed light on the mechanistic understanding of the process that can yield the state of energy deficit and concomitant changes in metabolic landscape as observed in our PTSD subjects. Although the analysis was focused on metabolic featues in PTSD, the mechanistic insights obtained from the model analysis can be generalized to glucocorticoid related diseases that involve metabolic dysfunction. Our findings of the defective energy metabolism also replicated the findings reported by other studies on patients with psychiatric disorders as reported by Zuccoli et al. (2017)^59^ and in PTSD specific population ^10^,^11^, implying mitochondrial dysfunction in PTSD. Our analysis indicates potential of targeting redox regulation ^60^, inflammation and modulation of GR sensitivity as a therapeutic alternatives for managing metabolic abnormalities in PTSD.

In this study, we have focused our analysis on perturbations in single process parameter at a time to analyze metabolic regulatory defects as observed in our PTSD sample, however, the possibility of perturbations in multiple procesess in PTSD is more likely and would need further exploration. The model could yield the MD signature for increasing the systemic GR sensitivity composed of GR-cytokine anti-inflammatory effect and central negative feedback effect, although by separately increasing the sensitivity of GR effects in these pathways produced opposite effects. The model reveals the MD signature for increasing the GR sensitivity of the cytokines but metabolites in our data did not show significant association with IC50. These discrepancies indicate a synergistic effect of the GR sensitivity of HPA-immune axis togather on the metabolism with the metabolic respone being sensitive to both an increase or decrease in the GR sensitivity. From the model analysis, our inferences are limited by what could be observed by model perturbations predominantly in hepatic metabolism; however the systemic effects may produce varied results. Moreover, modeling assumes proportionality between the serum and intracellular levels of metabolites, which may not be always true in reality. Not all individuals who are exposed to trauma will develop PTSD, therefore our design is observational. It would be practically infeasible and unethical to randomize individuals and expose them to trauma or modulate GR sensitivity in people. Hence, to test the model-based hypothesis, we used the results of dexamethasone suppression test as a proxy for systemic GR sensitivity in our subjects and the continuous exposure variable for causal inference. Since the causal structure assumed was an acyclic graph, it might be subject to weaker assumptions on sequential ignorability. Further controlled experiments are required to validate our hypothesis on the causal role of enhanced GC sensitivity in metabolic dysregulation in PTSD.

## Methods

### Participants

One hundred and sixty-five men between 20 and 60 years of age who served in Iraq (Operation Iraqi Freedom: OIF) or Afghanistan (Operation Enduring Freedom: OEF) were included in the analyses for this paper. Veterans were recruited to participate in the study as part of a systems biology approach to identify biomarkers for PTSD in OEF/OIF veterans. Participants were recruited at two sites including the James J. Peters Veterans Affairs Medical Center (JJPVAMC)/ Icahn School of Medicine at Mount Sinai (ISMMS), and New York University Langone Medical Center (NYULMC)/ NYU School of Medicine (NYUSM) through advertising in the clinic (VAMC) and community (local colleges and universities, vet centers, media advertisements). All participants provided written, informed consent for study procedures and the study was approved by the IRBs of the JJPVAMC, ISMMS, NYULMC and the Human Research Protection Office at the United States Army Medical Research and Materiel Command.

### Neuro-endocrine data

#### Procedures

All participants had exposure to a warzone-related DSM-IV PTSD Criterion a trauma while deployed and the presence of a diagnosis of PTSD was determined by a doctoral level psychologist using the Clinician Administered PTSD Scale (CAPS). Participants in the no-PTSD group were required to have a current (past month) CAPS scores ≤ 20 and had never met criteria for PTSD in the past. Participants in the PTSD group were required to have a current (past month) CAPS score ≥ 40 and to meet full DSM-IV criteria for PTSD. Participants were also asked to report their symptoms during the month when their symptoms were most distressing (lifetime). All cases were adjudicated in weekly consensus meetings across the two recruitment sites. The Structured Clinical Interview for DSM-IV (SCID) was used by the same clinician to determine other DSM-IV diagnoses. Participants with a lifetime history of any psychiatric disorder with psychotic features, bipolar disorder, current alcohol dependence, current drug abuse or dependence or obsessive-compulsive disorder, prominent suicidal or homicidal ideation or a suicide attempt in the past year were excluded. Medical exclusions included neurological disorder, loss of consciousness greater than 10 minutes, or other systemic illness affecting CNS function. Participants taking medications for psychiatric or medical conditions had to report consistent use for more than two months to be eligible to participate.

#### Blood sample collections and Dexamethasone Suppression Test

Participants reported to the laboratory at JJP VAMC or ISMMS between 7:30 and 8:00 after an overnight fast. Vital signs, weight, height and waist-hip ratio were measured and then approximately 160 cc of whole blood was collected and processed for subsequent assays, including the Dexamethasone Suppression Test (DST). Participants received a 0.50 mg tablet of Dexamethasone to ingest at 11:00 pm and returned the following morning (post-Dex) for collection of 10cc of blood. Blood samples were delivered to a CLIA certified lab at ISMMS or JJP VAMC for assessment of a variety of clinical labs (e.g., gamma-glutamyl transferase (GGT), high-sensitivity C-reactive protein (hs-CRP).

#### Lysozyme IC_50_-DEX

For the lysozyme IC_50_-DEX assay, mononuclear leukocytes were prepared immediately following the blood drawing procedure. For the preparation of mononuclear leukocytes, platelet-rich plasma was separated by low speed centrifugation. After collecting plasma, the remaining cells were diluted by the sample volume with Hanks’ Balanced Salt Solution (HBSS) and the lymphocytes were isolated by density centrifugation utilizing Ficoll-Paque (GE Healthcare) and washed twice in PBS according to the method of Boyum ^61^. The final cell pellet was re-suspended in a medium (RPMI-1640) containing 10% fetal calf serum, penicillin, streptomycin, and L-glutamate (Life Technologies, Grand Island, NY) at a density of 1.75-2.00 x 106 cells/ml. The test for examining the inhibition of lysozyme synthesis and release was carried out in 96-well culture plate in a total volume of.22 mL, modified from ^62^. Lysozyme activity was measured by turbidimetric method using Micrococcus lysodeikticus (Sigma) as the substrate. Micrococcus lysodeikticus was prepared in 0.1 mol/L phosphate buffer, pH 6.3, at a concentration of.05% and homogenized with a tissue grinder equipped with a Teflon pestle (Wheaton, St. Millville, New Jersey) with 3 strokes. 20 µL of supernatant of cell culture was incubated with 150 µL of substrate in a 96-well plate at 37°C for 7–10 min with shaking and then kinetically read by a microplate reader at 450 nm for 20 min. Cells (3.5-4.0 × 105) were incubated with 0,.5, 1, 2.5, 5, 10, 50, and 100 nmol/L of dexamethasone (DEX) (Sigma) at 37°C in a humidified atmosphere with 5% CO2 for 3 days. Each concentration of DEX was incubated in triplicate. After centrifuging the plate, 120 µL of supernatant were removed and pooled from each triplicate well. The standards were prepared using pure lysozyme from chicken egg white (Sigma) dissolved in RPMI-1640 as used for the cell culture. The inhibition curve was drawn as concentration of DEX versus relative activity of lysozyme. Results were expressed as IC_50_-DEX (nmol/L) based on the concentration of DEX at which 50% of lysozyme activity was inhibited. The intra- and inter-assay coefficients of variation for the measurement of lysozyme activity were 6.9% and 9.8% respectively ^63^.

#### Plasma Cortisol

Cortisol levels in plasma were assayed using Cortisol ELISA Kit from IBL-America (Minneapolis, MN), a solid phase enzyme-linked immunosorbent assay, based on the principle of competitive binding. The microtiter wells were coated with a monoclonal antibody directed towards an antigenic site on the cortisol molecule. Endogenous cortisol from an unknown competes with a cortisol-horseradish peroxidase conjugate for binding to the coated antibody. After incubation the unbound conjugate was washed off. The amount of bound peroxidase conjugate is inversely proportional to the concentration of cortisol in the unknown. After addition of the substrate solution, the intensity of color developed is inversely proportional to the concentration of cortisol in the unknown. Assay sensitivity: 2.5 ng/mL. The intra-assay and inter-assay coefficients of variation for this assay 5.3% and 9.8%, respectively. Two blood samples were assayed for the determination of cortisol before and after DEX administration. Decline of cortisol from Day 1 to Day 2 was used as a measure of DEX suppression.

#### Plasma ACTH

ACTH levels in plasma was assayed by using ACTH ELISA kit (ALPCO Diagnostics, Windham NH). In this assay, calibrators and research samples were simultaneously incubated with the enzyme labeled antibody and a biotin coupled antibody in a streptavidin-coated micro plate well. At the end of the assay incubation, the microwell was washed to remove unbound components and the enzyme bound to the solid phase was incubated with the substrate, tetramethylbenzidine (TMB). An acidic stop solution was added to stop the reaction and convert the color to yellow. The intensity of the yellow color is directly proportional to the concentration of ACTH in the sample. A dose response curve of absorbance unit vs. concentration was generated using results obtained from the calibrators. Concentrations of ACTH present in the samples was determined directly from this curve. Assay sensitivity: 0.5 pg/mL. The intra-assay and inter-assay coefficients of variation for this assay 5.7% and 8.0%, respectively. Two blood samples were assayed for the determination of ACTH before and after DEX administration.

#### Plasma DHEA

DHEA and DHEA-S were measured using ALPCO ELISA DHEA and DHEA-S kits (ALPCO Diagnostics, Windham NH). Both kits utilize a competitive immunoassay specifically designed and validated for the in vitro diagnostic measurement of dehydroepiandrosterone (DHEA) and dehydroepiandrosterone sulfate (DHEA-S) in human blood. Assay sensitivity for DHEA: 0.1 ng/mL. The intra-assay and inter-assay coefficients of variation for this assay 3.6% and 6.1%, respectively. Assay sensitivity for DHEA-S: 5.0 pg/ml. The intra-assay and inter-assay coefficients of variation for this assay 5.7% and 10.0%, respectively.

#### 24-hour Urine Collection

At the end of the first study visit, subjects were given instructions and materials to collect urine at home over 24 hours. Urine was kept in a freezer for the duration of the collection and kept frozen until it was returned to the laboratory.

#### Urinary catecholamines

Urinary catecholamines (E, NE, and DA) were extracted using Urinary Catecholamine Kit developed by Bioanalytical Systems, Inc (BAS). Extraction of catecholamines from 0.5 ml of urine sample was performed on the Solid Phase Extraction (SPE) Columns using the company’s proprietary reagents. 12µl of dihydroxybenzoic acid (DHBA) was added to each sample as an internal standard. HPLC analysis of the elute was performed on Thermo Scientific Dionex UltiMate 3000 with an autosampler and Dionex Coulochem III electrochemical detector. Quantitation was performed by integrating peak areas and comparing the ratios of the analyte to those of the internal standard with reference to a calibration curve across the range of concentrations using Chromeleon 7 Chromatography Data System.

#### Metabolomics and cytokine data data

The blood samples of 82 combat veterans without PTSD and 83 combat veterans who developed PTSD were used for metabolomics profiling. These two group served as controls and cases in our current study. The subjected reported to the laboratory in the morning 7.30 AM under fasting condition and the blood samples were colled around 8 AM. The metabolic profiling of the blood samples were performed by Metabolon, Inc. (Durham, NC). The primary metabolic data and details of the sample collection and metabolic profiling are reported in *Mellon et al*., (in review at PLOS one). The cytokine data was obtained as per our previous reports ^5^.

#### Statistical analysis

In the present analysis, we used the sample of 82 controls and 83 PTSD subjects for the statistical analysis. The data for metabolomics, neuro-endocrine, clinical labs and cytokines was log-transformed and median normalized before the statistical analysis. Due to heterogeneity in the distributions of some features in the data across cohorts, we used non-parametric Mann-Whitney U test to identify the features that show statistically significant difference in the PTSD and control groups. The statistical significance for all analysis was set at α=0.05 and trend level significance was set to 0.05<α<0.1. The false discovery rate for the multiple comparisons was reported by the q-values obtained from q-value package in R. All the statistical analysis was performed in R.

To assess the model based hypothesis we performed correlation analysis using Spearman Correlation Coefficients (ρ) between the regulatory components and the metabolites that show statistically significant differences in PTSD case versus controls. Correlation matrices for both controls and PTSD samples were obtained separately and a relative difference (Rd) of the correlations with respect to correlations in control subjects were estimated after rescaling the two matrices in the range of 1 to 10. The relative difference was obtained by,

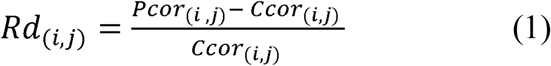

where, *Pcor_i_* and *Ccor_i_* represent the (*i,j)th* correlation coefficient in the correlation matrix of *i* number of rows and *j* number of columns for PTSD and control, respectively. We used the psych package in R to estimate correlation coefficients and the corresponding p-values and the R package corrplot to plot correlation plots ^64^.

#### Mathematical model development

To analyze the defects in metabolism we used a published mathematical model of the human liver to simulate metabolic trends. To study the regulatory effects of the HPA axis and inflammation on metabolism, we further integrated the metabolic simulator with the models for HPA axis, inflammation and hypoxia. We adopted four sub models published in literature, namely hepatic metabolism ^65^, HPA axis and inflammation model ^66^, glucocorticoid receptor model ^67^, and hypoxia signaling by ^68^. These models were integrated by linking the regulatory nodes according to the interactions reported in the literature. We used a semi-empirical approach to incorporate regulatory effects by using the saturating rate equations modeled by Michalis-Menten and Hill type of biochemical kinetics. In the semi-empirical approach, we emphasize on incorporating the known regulatory interactions from the literature and reproducing the qualitative trends in input-output physiological response. The equations were formulated to satisfy the experimentally observed physiological responses at basal conditions and the qualitative behavior on changing the regulatory stimulus. The regulatory functions were applied to metabolic pathways with an assumption of parallel activation mechanism, wherein, if a rate limiting step in a pathway is regulated by a regulator, the regulatory effect is applied to series of reactions in the pathway until a branch point or a store or a sink, or the next regulated variable of the pathway.

The source of the additional interactions in the integrated model and the scheme of model development are reported in the Appendix II-Table S2. The model is composed of ODEs which can simulate the dynamical profiles for metabolites, signaling and transcriptional regulatory components in the metabolic regulatory network. The model is comprised of major metabolic pathways of glycolysis, TCA cycle, amino acid metabolism, urea cycle, lipid metabolism and plasma metabolite transport along with regulatory signaling pathways for insulin (IRS-AKT), glucagon and catecholamine (GPCR-cAMP-PKA), HPA axis for cortisol and glucocorticoid receptor (GRs) dynamics, inflammation and hypoxia (Appendix II : Figure S2 & S3). The model was validated to reproduce the qualitative physiological responses for different input conditions (Appendix II : Figure S4). The overall model is comprised of 189 ODEs for the state variables. The model was developed and analyzed using MATLAB (2017a).

#### Metabolic control analysis

Metabolic control analysis ^27^ was performed by perturbation of parameters in the model, wherein parameters represents rate of reactions and strength of interaction in the metabolic and regulatory network, respectively. We performed the metabolic control analysis to obtain concentration response coefficients to identify the rate parameters that can yield the metabolic signatures as showed by statistically significant group differences in subjects with and without PTSD. To quantify the changes in metabolite concentrations with respect to changes in the parameters for reactions or regulatory interactions we calculate metabolite concentration response coefficients (MCRCs) and as,

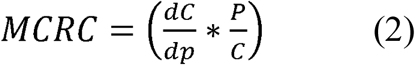

where, *C* and *P* are the concentration of a metaboliteand parameters for reaction rates in the model, respectively. We recorded MCRCs for 100 metabolic reaction rates and 260 signaling and transcriptional regulatory interaction rates, with respect to the trends observed in 12 metabolites that were identified significantly different in PTSD dataset. Due to the nonlinear nature of the regulatory influences of hormonal signaling and transcriptional factors, instead of infinitesimal perturbation we obtained the response coefficients for a modest perturbation across 50% change in the reaction and interaction rates (i.e. 1.5 fold and 0.5 fold of the native value). We obtained the net response coefficnet by taking the mean of response coeffeicinets recoreded for decreasing (0.5 fold) and increasing (1.5 fold) the parameter across the nominal values, with at least MCRC of 0.001 for each of the 12 metabolites. The steady state simulation results for those 12 metabolites were compared with the pattern of change in direction of significantly different metabolites in PTSD subjects. The directions of the change in metabolite concentrations (positive or negative) with respect to baseline levels for a particular rate perturbation were used to determine whether a concentration of a metabolite would decrease or increase for a perturbation in a rate of reaction. The control coefficients that yielded all the changes as observed in the MD signature were recorded for further inferences related to the disease. The simulations for MCA were carried out under contant glycogen levels to mimic the normal supply of energy source and maintain steady state levels in the model.

#### Causal inference and sensitivity analysis

We used the covariate balancing propensity scores (CBPS) ^19^ package in R to obtain the weights for covariate/confounder adjustments while deducing causal effects. The covariate balancing generalized propensity scores obtained from CBPS are optimized to maximize both the covariate balance and prediction of treatment assignment in the sample by minimizing the association between covariates and the treatment/exposure variable. A non-parametric version called npCBPS was used to obtain the weights for fitting the average causal effects (ACEs). To obtain the population level causal estimates we used the entire sample for analysis by controlling for the group effects. We computed the average causal effects (ACE-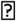) of six regulatory components using the CBPS package, with respect to 35 metabolites in our sample. The ACEs (represented by (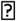) gamma) can be interpreted as the percent change in the affected metabolite per unit percent change in the regulatory component. ACEs capture the average causal association between two variables controlling for covariates without referring to the direction of causality.

We used treatSens package in R ^69^ to perform sensitivity analysis of the causal estimates for unmeasured confounding. The sensitivity analysis (SA) employs a simulation based non-parametric method, wherein a sensitivity parameter is the coefficient of the association between the unknown confounder and the treatment and outcome variables. The method determines the sensitivity estimates that can account for the bias due to model misspecification and unmeasured confounding. The data was standardized for the SA to obtain the sensitivity coefficient ranging from 0-1. The sensitivity is assessed graphically by plotting the 2D plot of the causal estimates with respect to bi-parametric variation (correlation of unknown confounder with treatment and outcome variables) ^20^. We recorded sensitivity parameters representing the intersection of the bi-parametric curve with the x=y line on the plot. Two sensitivity parameters were recorded, namely τ_1_: the point of intersection of the x=y line with the curve that represents the causal estimate equals to zero; τ_2_: the point of intersection of the x=y line with the curve that represents the lack of significance (where p>0.05) of the causal estimate. The sensitivity parameters were evaluated only for the statistically significant causal associations.

#### Causal mediation analysis

We used the Medflex package in R ^70^ to perform causal mediation analysis (CMA) using natural effect models ^25^. The package employs computations on conditional mean models for nested counterfactuals to estimate the causal effects. These models allow for estimation of natural direct and indirect or mediated effects (NDE and NIE) through its coefficients, which provides easier interpretation of effects with respect to the exposure variables. The CMA uses a counterfactual framework of causal inference which works by identifying the difference between the potential outcomes of a design or a graph with and without a mediator estimated by following equations,

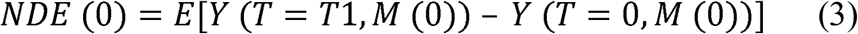

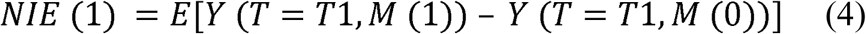

where, *Y, T* and *M* are the outcome variable, treatment or exposure variable and mediator variable, respectively. The suffix 1 and 0 in the parenthesis represent the notation for with and without the change in treatment or mediator variables. To obtain generalized estimates of the causal effects we performed a population level mediation analysis on the entire cohort controlling for the disease status. The population level effects were thus obtained by weighting by the reciprocal of the conditional exposure density of the treatment variable through inverse probability weighting in the functional models. The causal inference problem is based on assumptions of sequential ignorability on the causal structure invoked by non-parametric structural equation models with independent error terms (NPSEM-IE) ^71^. The package employs generalized linear models (glm) for mediation analysis and 1000 non-parametric bootstrapped simulations to generate inference and related standard errors. The natural direct effect (NDE: *ψ*_*d*_) is interpreted as, for a subject with baseline covariates, one percent change from average level of cortisol suppression results in a percent change in a corresponding metabolite level by the factor of NDE. The natural indirect effect (NIE: *ψ*_*i*_) can be interpreted as, for altering the level from that would have been observed at the average levels of cortisol suppression to the level that would have been observed at one percent change in cortisol suppression would result in the percent change in metabolite level by the factor of NIE. The total causal effect (TCE: *ψ*_*t*_) is the sum of the direct and indirect effects.

## Supporting information

## Data availability

The metabolomics, neuroendocrine and cytokine data analyzed during this study are available at https://sysbiocube-abcc.ncifcrf.gov/ and will be made available on reasonable request.

## Software and code availability

The codes used for the analysis in this study will be made available on reasonable request.

## Supplemental Information

Appendix-I : Outline of work and demographics of the cohort

Appendix-II : Mathematical model development

Appendix-III : Metabolic control analysis

Appendix-IV : Correlational Analysis

Appendix-V : Causal mediation inference

## Acknowledgement

This work was supported by a research grant from U.S Army Medical Research and Material Command (USAMRMC) under award number W911NF-17-2-0086, W81XWH-10-1-0021 (PI: Wolkowitz) and W911NF-17-1-0069 (PI: Daigle). We are thankful to Dr. Sunil Deshpande (Harvard University) and Mr. Christian Fong (Stanford University) for providing methodological insights on causal analysis. We are thankful to Dr. David Jackson and Dr.Mathew Kobe from US ARMY for reviewing and editing the manuiscript. We also thank to Dr. Kerry Ressler, and other PTSD Biomarker consortium members for group discussions.

## Author contributions

PRS conceived the work, performed the analysis and wrote the manuscript. SM and OW generated the metabolomics data and supervised the manuscript. FJ and RY generated the neuroendocrine data and supervised the manuscript. DA and CM recruited the subjects and generated clinical data. FJD, MJ and CM supervised the study. All the PTSD consortium authors reviewed and edited the manuscript.

## Disclaimer

The views, opinions, and findings contained in this report are those of the authors and should not be construed as official Department of the Army position, policy, or decision, unless so designated by other official documentation. Citations of commercial organizations or trade names in this report do not constitute an official Department of the Army endorsement or approval of the products or services of these organizations.

## References

1 DSM-5. American Psychiatric Association. Diagnostic and statistical Manual of Mental Disorders. American Psychiatric Association Fifth Edition (2013).

2 Chakraborty, N., Meyerhoff, J., Jett, M. & Hammamieh, R. in Neuroproteomics: Methods and Protocols (eds Firas H. Kobeissy & Jr Stanley M. Stevens) 117–154 (Springer New York, 2017).

3 Shalev, A., Liberzon, I. & Marmar, C. Post-Traumatic Stress Disorder. N. Engl. J. Med. 376, 2459–2469, doi:10.1056/NEJMra1612499 (2017).

4 Blessing, E. M. et al. Biological predictors of insulin resistance associated with posttraumatic stress disorder in young military veterans. Psychoneuroendocrinology 82, 91–97, doi:https://doi.org/10.1016/j.psyneuen.2017.04.016 (2017).

5 Lindqvist, D. et al. Increased pro-inflammatory milieu in combat related PTSD – A new cohort replication study. Brain. Behav. Immun. 59, 260–264, doi:https://doi.org/10.1016/j.bbi.2016.09.012 (2017).

6 Bersani, F. S. et al. Mitochondrial DNA copy number is reduced in male combat veterans with PTSD. Prog. Neuropsychopharmacol. Biol. Psychiatry 64, 10–17, doi:https://doi.org/10.1016/j.pnpbp.2015.06.012 (2016).

7 Yehuda, R. et al. Lower Methylation of Glucocorticoid Receptor Gene Promoter 1F in Peripheral Blood of Veterans with Posttraumatic Stress Disorder. Biol. Psychiatry 77, 356–364, doi:https://doi.org/10.1016/j.biopsych.2014.02.006 (2015).

8 Mellon, S. H., Gautam, A., Hammamieh, R., Jett, M. & Wolkowitz, O. M. Metabolism, Metabolomics, and Inflammation in Post-Traumatic Stress Disorder. Biol. Psychiatry 83, 866–875, doi:https://doi.org/10.1016/j.biopsych.2018.02.007 (2018).

9 Preston, G., Kirdar, F. & Kozicz, T. The role of suboptimal mitochondrial function in vulnerability to post-traumatic stress disorder. J. Inherit. Metab. Dis. 41, 585–596, doi:10.1007/s10545-018-0168-1 (2018).

10 Flaquer, A. et al. Mitochondrial genetic variants identified to be associated with posttraumatic stress disorder. Translational Psychiatry 5, e524, doi:10.1038/tp.2015.18 (2015).

11 Su, Y. A. et al. Dysregulated Mitochondrial Genes and Networks with Drug Targets in Postmortem Brain of Patients with Posttraumatic Stress Disorder (PTSD) Revealed by Human Mitochondria-Focused cDNA Microarrays. Int. J. Biol. Sci. 4, 223–235 (2008).

12 Yehuda, R. Post-Traumatic Stress Disorder. N. Engl. J. Med. 346, 108–114, doi:10.1056/NEJMra012941 (2002).

13 Lukaschek, K. et al. Relationship between posttraumatic stress disorder and Type 2 Diabetes in a population-based cross-sectional study with 2970 participants. J. Psychosom. Res. 74, 340–345, doi:https://doi.org/10.1016/j.jpsychores.2012.12.011 (2013).

14 Logue, M. W. et al. An analysis of gene expression in PTSD implicates genes involved in the glucocorticoid receptor pathway and neural responses to stress. Psychoneuroendocrinology 57, 1–13, doi:10.1016/j.psyneuen.2015.03.016 (2015).

15 Wester, V. L. et al. Glucocorticoid receptor haplotype and metabolic syndrome: the Lifelines cohort study. European Journal of Endocrinology 175, 645–651, doi:10.1530/eje-16-0534 (2016).

16 Yehuda, R. Status of Glucocorticoid Alterations in Post-traumatic Stress Disorder. Ann. N. Y. Acad. Sci. 1179, 56–69, doi:10.1111/j.1749-6632.2009.04979.x (2009).

17 Koenen, K. C. et al. Post-traumatic stress disorder and cardiometabolic disease: improving causal inference to inform practice. Psychol. Med. 47, 209–225, doi:10.1017/S0033291716002294 (2017).

18 Winship, C. & Morgan, S. L. The Estimation of Causal Effects from Observational Data. Annu. Rev. Sociol. 25, 659–706, doi:10.1146/annurev.soc.25.1.659 (1999).

19 Christian Fong, C. H., Kosuke Imai. Covariate Balancing Propensity Score for a Continuous Treatment: Application to the Efficacy of Political Advertisements. Annals of Applied Statistics 12, 156–177 (2018).

20 Dorie, V., Harada, M., Carnegie, N. B. & Hill, J. A flexible, interpretable framework for assessing sensitivity to unmeasured confounding. Stat. Med. 35, 3453–3470, doi:10.1002/sim.6973 (2016).

21 Leistner, C. & Menke, A. How to measure glucocorticoid receptor’s sensitivity in patients with stress-related psychiatric disorders. Psychoneuroendocrinology 91, 235–260, doi:https://doi.org/10.1016/j.psyneuen.2018.01.023 (2018).

22 Saugstad, O. D. Hypoxanthine as an Indicator of Hypoxia: Its Role in Health and Disease through Free Radical Production. Pediatr. Res. 23, 143–150, doi:10.1203/00006450-198802000-00001 (1988).

23 Holden, M. S. et al. Urinary Hypoxanthine as a Measure of Increased ATP Utilization in Late Preterm Infants. Infant Child Adolesc. Nutr. 6, 240–249, doi:10.1177/1941406414526618 (2014).

24 Koenig, G. & Seneff, S. Gamma-Glutamyltransferase: A Predictive Biomarker of Cellular Antioxidant Inadequacy and Disease Risk. Dis. Markers 2015, 18, doi:10.1155/2015/818570 (2015).

25 Steen, J., Loeys, T., Moerkerke, B. & Vansteelandt, S. Flexible Mediation Analysis With Multiple Mediators. Am. J. Epidemiol. 186, 184–193, doi:10.1093/aje/kwx051 (2017).

26 VanderWeele, T. J. & Vansteelandt, S. Mediation Analysis with Multiple Mediators. Epidemiologic methods 2, 95–115, doi:10.1515/em-2012-0010 (2014).

27 Fell, D. A. in Systems Biology: Definitions and Perspectives (eds Lila Alberghina & H. V. Westerhoff) 69–80 (Springer Berlin Heidelberg, 2005).

28 Rabasa, C. & Dickson, S. L. Impact of stress on metabolism and energy balance. Current Opinion in Behavioral Sciences 9, 71–77, doi:https://doi.org/10.1016/j.cobeha.2016.01.011 (2016).

29 Kuo, T., McQueen, A., Chen, T.-C. & Wang, J.-C. in Glucocorticoid Signaling: From Molecules to Mice to Man (eds Jen-Chywan Wang & Charles Harris) 99–126 (Springer New York, 2015).

30 Dengler, V. L., Galbraith, M. & Espinosa, J. M. Transcriptional Regulation by Hypoxia Inducible Factors. Crit. Rev. Biochem. Mol. Biol. 49, 1–15, doi:10.3109/10409238.2013.838205 (2014).

31 Bartels, K., Grenz, A. & Eltzschig, H. K. Hypoxia and inflammation are two sides of the same coin. Proc. Natl. Acad. Sci. U. S. A. 110, 18351–18352, doi:10.1073/pnas.1318345110 (2013).

32 Wei, K. et al. A liver HIF-2α/IRS2 pathway sensitizes hepatic insulin signaling and is modulated by VEGF inhibition. Nat. Med. 19, 1331–1337, doi:10.1038/nm.3295 (2013).

33 Lee, Dong C. et al. A Lactate-Induced Response to Hypoxia. Cell 161, 595–609, doi:https://doi.org/10.1016/j.cell.2015.03.011 (2015).

34 Derks, T. G. J. et al. Inhibition of mitochondrial fatty acid oxidation in vivo only slightly suppresses gluconeogenesis but enhances clearance of glucose in mice. Hepatology 47, 1032–1042, doi:10.1002/hep.22101 (2008).

35 Letteron, P. et al. Glucocorticoids inhibit mitochondrial matrix acyl-CoA dehydrogenases and fatty acid beta-oxidation. Am. J. Physiol. Gastrointest. Liver Physiol. 272, G1141–G1150, doi:10.1152/ajpgi.1997.272.5.G1141 (1997).

36 Liu, Y. et al. HIF-1α and HIF-2α are critically involved in hypoxia-induced lipid accumulation in hepatocytes through reducing PGC-1α-mediated fatty acid β-oxidation. Toxicol. Lett. 226, 117–123, doi:https://doi.org/10.1016/j.toxlet.2014.01.033 (2014).

37 Zhang, L. et al. Mitochondria-focused gene expression profile reveals common pathways and CPT1B dysregulation in both rodent stress model and human subjects with PTSD. Translational Psychiatry 5, e580, doi:10.1038/tp.2015.65 (2015).

38 Gathercole, L. L., Bujalska, I. J., Stewart, P. M. & Tomlinson, J. W. Glucocorticoid Modulation of Insulin Signaling in Human Subcutaneous Adipose Tissue. The Journal of Clinical Endocrinology and Metabolism 92, 4332–4339, doi:10.1210/jc.2007-1399 (2007).

39 Peckett, A. J., Wright, D. C. & Riddell, M. C. The effects of glucocorticoids on adipose tissue lipid metabolism. Metabolism 60, 1500–1510, doi:https://doi.org/10.1016/j.metabol.2011.06.012 (2011).

40 Denton, R. M. et al. The hormonal regulation of pyruvate dehydrogenase complex. Adv. Enzyme Regul. 36, 183–198, doi:https://doi.org/10.1016/0065-2571(95)00020-8 (1996).

41 Burt, M. G., Gibney, J. & Ho, K. K. Y. Protein metabolism in glucocorticoid excess: study in Cushing’s syndrome and the effect of treatment. Am. J. Physiol. Endocrinol. Metab. 292, E1426–E1432, doi:10.1152/ajpendo.00524.2006 (2007).

42 Christian, H., Stephan, D., Florian, L., Wolfgang, G. & Dieter, H. Inhibition of hepatic proteolysis by insulin. European Journal of Biochemistry 199, 467–474, doi:doi:10.1111/j.1432-1033.1991.tb16145.x (1991).

43 Sun, R. C. & Denko, N. C. Hypoxic regulation of glutamine metabolism through HIF1 and SIAH2 supports lipid synthesis that is necessary for tumor growth. Cell Metabolism 19, 285–292, doi:10.1016/j.cmet.2013.11.022 (2014).

44 Calbet, J. A. L. & MacLean, D. A. Plasma Glucagon and Insulin Responses Depend on the Rate of Appearance of Amino Acids after Ingestion of Different Protein Solutions in Humans. The Journal of Nutrition 132, 2174–2182, doi:10.1093/jn/132.8.2174 (2002).

45 Watford, M. The urea cycle: Teaching intermediary metabolism in a physiological setting. Biochem. Mol. Biol. Educ. 31, 289–297, doi:10.1002/bmb.2003.494031050249 (2003).

46 Okun, J. G. et al. Molecular regulation of urea cycle function by the liver glucocorticoid receptor. Molecular Metabolism 4, 732–740, doi:10.1016/j.molmet.2015.07.006 (2015).

47 Bersani, F. S. et al. Global arginine bioavailability, a marker of nitric oxide synthetic capacity, is decreased in PTSD and correlated with symptom severity and markers of inflammation. Brain. Behav. Immun. 52, 153–160, doi:https://doi.org/10.1016/j.bbi.2015.10.015 (2016).

48 Groeneweg, F. L., Karst, H., de Kloet, E. R. & Joëls, M. Rapid non-genomic effects of corticosteroids and their role in the central stress response. Journal of Endocrinology 209, 153–167, doi:10.1530/joe-10-0472 (2011).

49 Görlach, A., Bertram, K., Hudecova, S. & Krizanova, O. Calcium and ROS: A mutual interplay. Redox Biology 6, 260–271, doi:10.1016/j.redox.2015.08.010 (2015).

50 Mungai, P. T. et al. Hypoxia Triggers AMPK Activation through Reactive Oxygen Species-Mediated Activation of Calcium Release-Activated Calcium Channels. Molecular and Cellular Biology 31, 3531–3545, doi:10.1128/MCB.05124-11 (2011).

51 Duque, E. d. A. & Munhoz, C. D. The Pro-inflammatory Effects of Glucocorticoids in the Brain. Front. Endocrinol. (Lausanne) 7, 78, doi:10.3389/fendo.2016.00078 (2016).

52 Newton, R., Shah, S., Altonsy, M. O. & Gerber, A. N. Glucocorticoid and cytokine crosstalk: Feedback, feedforward, and co-regulatory interactions determine repression or resistance. Journal of Biological Chemistry 292, 7163–7172, doi:10.1074/jbc.R117.777318 (2017).

53 Bjelaković, G. et al. in J. Basic Clin. Physiol. Pharmacol. Vol. 18 115 (2007).

54 Chaudhari, N., Talwar, P., Parimisetty, A., Lefebvre d’Hellencourt, C. & Ravanan, P. A Molecular Web: Endoplasmic Reticulum Stress, Inflammation, and Oxidative Stress. Front. Cell. Neurosci. 8, 1–15, doi:10.3389/fncel.2014.00213 (2014).

55 Chen, L., Chen, R., Wang, H. & Liang, F. Mechanisms Linking Inflammation to Insulin Resistance. Int. J. Endocrinol. 2015, 1–9, doi:10.1155/2015/508409 (2015).

56 Mühle, C., Reichel, M., Gulbins, E. & Kornhuber, J. in Sphingolipids in Disease (eds Erich Gulbins & Irina Petrache) 431–456 (Springer Vienna, 2013).

57 Maceyka, M., Harikumar, K. B., Milstien, S. & Spiegel, S. Sphingosine-1-phosphate signaling and its role in disease. Trends Cell Biol. 22, 50–60, doi:10.1016/j.tcb.2011.09.003 (2012).

58 Apostolopoulou, M. et al. Specific Hepatic Sphingolipids Relate to Insulin Resistance, Oxidative Stress, and Inflammation in Nonalcoholic Steatohepatitis. Diabetes Care 41, 1235–1243, doi:10.2337/dc17-1318 (2018).

59 Zuccoli, G. S., Saia-Cereda, V. M., Nascimento, J. M. & Martins-de-Souza, D. The Energy Metabolism Dysfunction in Psychiatric Disorders Postmortem Brains: Focus on Proteomic Evidence. Front. Neurosci. 11, 493:491-414, doi:10.3389/fnins.2017.00493 (2017).

60 Schiavone, S. & Trabace, L. Pharmacological targeting of redox regulation systems as new therapeutic approach for psychiatric disorders: A literature overview. Pharmacol. Res. 107, 195–204, doi:https://doi.org/10.1016/j.phrs.2016.03.019 (2016).

61 Boyum, A. 1-stage procedure for isolation of granulocytes & lymphocytes from human blood. Sedimentation of white blood cells in a 1g gravity field. Scand. J. Clin. Lab. Invest. Suppl. 97, 51–76 (1968).

62 Panarelli, M., Holloway, C. D., Mulatero, P., Fraser, R. & Kenyon, C. J. Inhibition of lysozyme synthesis by dexamethasone in human mononuclear leukocytes: an index of glucocorticoid sensitivity. The Journal of Clinical Endocrinology and Metabolism 78, 872–877, doi:10.1210/jcem.78.4.8157714 (1994).

63 Yehuda, R., Golier, J. A., Yang, R.-K. & Tischler, L. Enhanced sensitivity to glucocorticoids in peripheral mononuclear leukocytes in posttraumatic stress disorder. Biol. Psychiatry 55, 1110–1116, doi:https://doi.org/10.1016/j.biopsych.2004.02.010 (2004).

64 Wei, T. & Simko, V. R package “corrplot”: Visualization of a Correlation Matrix (Version 0.84). https://github.com/taiyun/corrplot (2017).

65 Somvanshi, P. R., Patel, A. K., Bhartiya, S. & Venkatesh, K. V. Influence of plasma macronutrient levels on hepatic metabolism: role of regulatory networks in homeostasis and disease states. RSC Advances 6, 14344–14371, doi:10.1039/C5RA18128C (2016).

66 Bangsgaard, E. O., Hjorth, P. G., Olufsen, M. S., Mehlsen, J. & Ottesen, J. T. Integrated Inflammatory Stress (ITIS) Model. Bull. Math. Biol. 79, 1487–1509, doi:10.1007/s11538-017-0293-2 (2017).

67 Rao, R., DuBois, D., Almon, R., Jusko, W. J. & Androulakis, I. P. Mathematical modeling of the circadian dynamics of the neuroendocrine-immune network in experimentally induced arthritis. Am. J. Physiol. Endocrinol. Metab. 311, E310–E324, doi:10.1152/ajpendo.00006.2016 (2016).

68 Nguyen, L. K. et al. A dynamic model of the hypoxia-inducible factor 1α (HIF-1α) network. J. Cell Sci. 126, 1454–1463, doi:10.1242/jcs.119974 (2013).

69 Carnegie, N. B., Harada, M. & Hill, J. treatSens: A Package to Assess Sensitivity of Causal Analyses to Unmeasured Confounding. Version R package version 2.1.3. (2018).

70 Steen, J., Loeys, T., Moerkerke, B. & Vansteelandt, S. medflex: An R Package for Flexible Mediation Analysis using Natural Effect Models. Journal of Statistical Software 76, 1–46, doi:doi:10.18637/jss.v076.i11 (2017).

71 Pearl, J. An Introduction to Causal Inference. The International Journal of Biostatistics 6, 7, doi:10.2202/1557-4679.1203 (2010).

