## Supplementary material for "Mechanistic insights on metabolic dysfunction in PTSD: Role of glucocorticoid receptor sensitivity and energy deficit"

**Appendix I: Outline and Demographics**

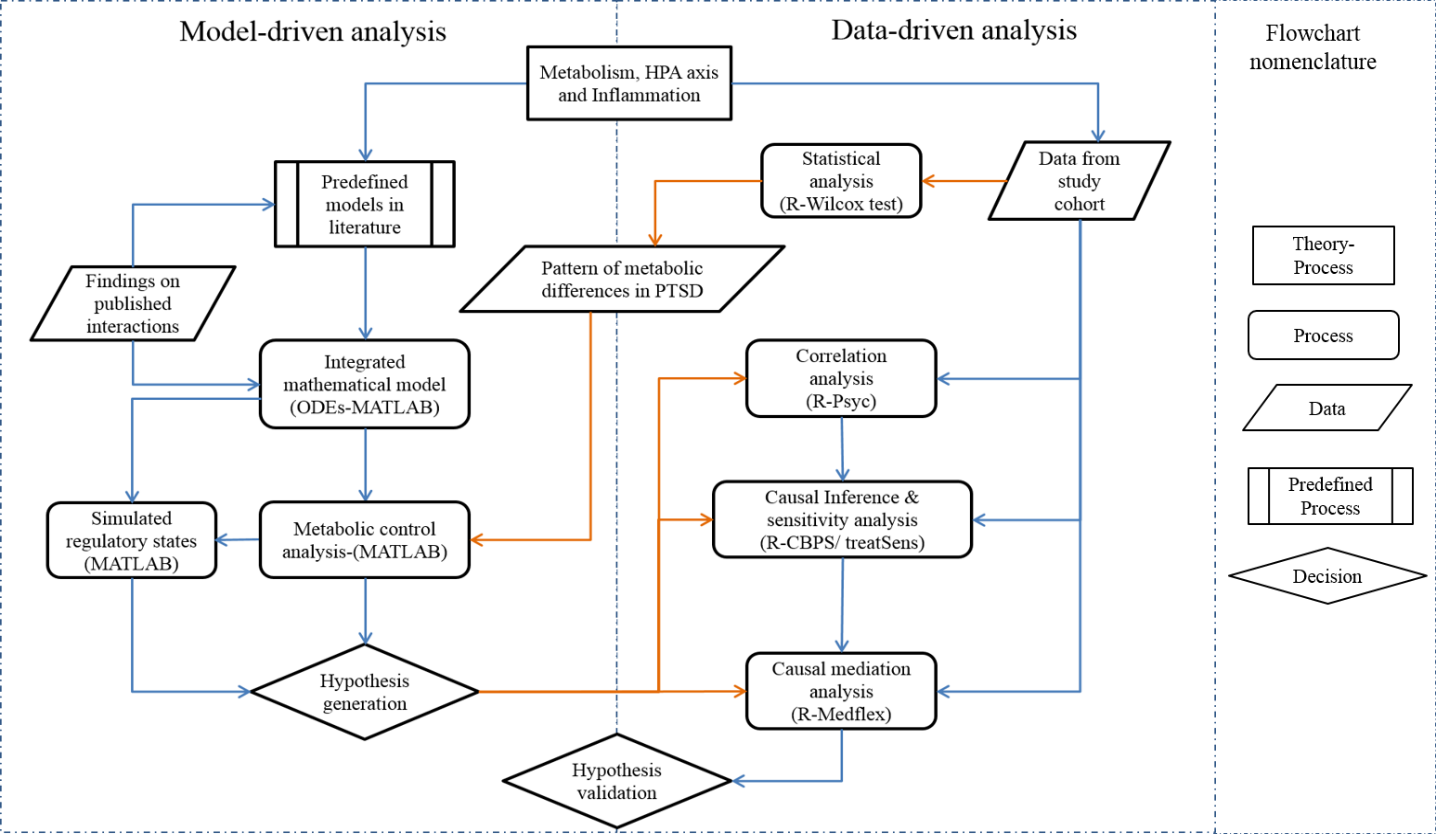

Figure S1: Process flowchart representing an outline of the steps that were implemented in the analysis presented in the paper. The published mathematical models for metabolism, HPA axis, inflammation and hypoxia signaling were integrated into a single composite model based on the data reported in literature for crosstalk and feedback regulations between the sub-models. The model was recalibrated and metabolic control analysis was performed to obtain metabolic concentration control coefficients and the associated regulatory states with reference to the pattern of metabolite differences observed in subjects with PTSD using statistical analysis. Orange arrows represent how the observations from data were used for model analysis and inference. The control coefficients were used to generate hypothesis on the potential process that could be affected in PTSD. To validate the hypotheses, correlation analysis followed by causal inference and mediation analysis was performed using models for propensity score estimation and natural effects models. The average causal effects were further augmented with sensitivity analysis to record the degree of violations of ignorability assumption.

Table S1. Demographic and clinical measures of combat veterans with PTSD and controls

| **Demographics** | **PTSD - (N:82)** | **PTSD + (N:83)** | **Demographics** | **PTSD - (N:82)** | **PTSD + (N:83)** |
| --- | --- | --- | --- | --- | --- |
| **Sociodemographic** |  |  | **Medication use (n)** | | |
| Gender | All males | All males | Sedatives | 4 | 16 |
| Age (Mean±SD) | 32.29 ± 7.60 | 33.20 ± 8.12 | Statins | 1 | 4 |
| Years of Education (Mean±SD) | 14.78 ± 2.32 | 13.81 ± 1.93 | Anti-depressants | 4 | 24 |
| Hispanic/Non-Hispanic (n) | 26/56 | 38/45 | Anticonvulsants | 0 | 9 |
| Smoking (n) | 18 | 33 | Anti-inflammatories | 6 | 9 |
| Alcohol use (Mean±SD) | 1.65 ± 1.08 | 1.38 ± 1.22 | Anti-diabetics | 2 | 2 |
| **Biometric measurements (Mean± SD)** | |  | Anti-hypertensives | 5 | 6 |
| BMI | 28.44 ± 4.79 | 30.02 ± 5.047 | Antacids | 3 | 3 |
| Weight | 192.78 ± 35.46 | 205.09 ±37.77 | Anti-allergics | 4 | 5 |
| Height | 69.05 ± 2.89 | 69.26 ± 2.84 | Pain medicines | 4 | 10 |
| Waist to Hip ratio | 0.88 ± 0.15 | 0.89 ± 0.16 | **Comorbid diseases (n)** |  |  |
| Pulse | 64.68 ± 11.21 | 72.65 ± 10.33 | Clinical hypertention | 7 | 15 |
| **Metabolic measurement (Mean±SD)** | |  | Heart attack | 1 | 1 |
| HbA1c | 5.37 ± 0.44 | 5.39 ± 0.85 | Stable angina | 1 | 3 |
| Cholesterol | 171.35 ± 27.44 | 180.21 ± 35.97 | Diabetics | 2 | 4 |
| LDL | 49.79 ± 13.08 | 47.27 ± 12.05 |  |  |  |
| HDL | 100.56 ± 25.12 | 107.96 ± 32.49 |  |  |  |
| **Clinical measures (Mean±SD)** |  |  |  |  |  |
| CAPS total current | 9.24 ± 8.39 | 91.55 ± 15.8 |  |  |  |
| CAPS total lifetime | 3.73 ± 5.01 | 69.42 ± 16.91 |  |  |  |
| PCLSCORE | 25.93 ± 8.95 | 61.51 ± 11.84 |  |  |  |
| MCS | 64.34 ± 14.21 | 118.53 ± 19.69 |  |  |  |
| PSQI | 5.19 ± 4.02 | 7.46 ± 5.72 |  |  |  |
| ETISR | 5.98 ± 3.77 | 13.25 ± 3.28 |  |  |  |
| Number of deployment | 1.77 ± 0.89 | 1.82 ± 0.84 |  |  |  |
| MDD diagnosis (n) | 1 | 47 |  |  |  |

**Appendix II : Mathematical Model Development**

The general scheme of model formulations implemented the methodology followed in (Bulik et al., 2016; König et al., 2012; Li et al., 2010; Somvanshi et al., 2016), wherein the concentrations of metabolic state variables were modeled by ordinary differential equations (ODEs),

$\frac{{dM}_{i}}{dt}*V=\sum_{j=1}^{nj} {RP}_{j}-\sum_{k=1}^{nk} {RC}_{k}\pm T_{t}$	(1)

where *M_i_* represents the *i*th metabolite, *V* represents the volume of the compartment to which the metabolite belongs; *RP_j_* and *RC_k_* are the rate of production and consumption of a metabolite, respectively. *T_t_* represents the transport of metabolite across blood and tissue. The rates of formation and consumption are further modelled as

${RP}_{j}={V_{max}}_{j}*{F\_RP}_{j}*\prod_{f=1}^{nfj} \left( \frac{M_{f,j}}{M_{f,j}+{Km}_{f,j}} \right)$	(2)

${F\_RP}_{j}=\prod_{r=1}^{nrj} \left( {F_{Act}}_{r,j}*{F_{Deact}}_{r,j}*F_{pi}*{F\_Sig\_trans}_{r,j} \right)$	(3)

${F_{Act}}_{r,j}=\left( \frac{A}{A+K_{j}} \right) ; {F_{Deact}}_{r,j}=\left( \frac{K_{i}}{I+K_{i}} \right) ;$	$F_{pi}=\left( \frac{A}{A+Ki*\left( 1+\frac{I}{Kp} \right)} \right)$	(4)

${F_{Sig_{trans}}}_{r,j}=W_{f}*\left( 1+\sum_{a}^{an} {{Reg}_{s}}_{{act}_{a}}+\sum_{b}^{bn} {{Reg}_{t}}_{{act}_{b}} \right)*\prod_{d}^{dn} \left( {Reg}_{s}\_{deact}_{d}*{Reg}_{t}\_{deact}_{d} \right)$ (5)

${Reg}_{st}\_{act}_{j}=\prod_{p}^{pn} \left( \frac{{S_{p}}^{n}}{{S_{p}}^{n}+{K_{p}}^{n}} \right)$(6)

$${Reg}_{s}\_{deact}_{j}=\prod_{q}^{qn} \left( \frac{{K_{p}}^{n}}{{I_{p}}^{n}+{K_{p}}^{n}} \right) (7)$$

$T_{tjpassive}=\epsilon_{j}*(C_{bj}-C_{cytj})$	(8)

$T_{tjFaciltated}=T_{j}*\left( \frac{C_{bj}}{K_{bj}+C_{bj}}-\frac{C_{cytj}}{K_{cytj}+C_{cytj}} \right)$	(9)

where ${F\_RP}_{j}$ is the regulatory function governing the rate of j the enzyme, $M_{f,j}$ is the *f*th metabolite in *j*th reaction in Eqn. (2). ${F_{Act}}_{r,j}$, ${F_{Deact}}_{r,j}$ and $F_{pi}$ represent the activation rate, deactivation rate and product inhibition rate by *r*th metabolite in the *j*th reaction in Eqn. (3 and 4), *A* and *I* represent the activator metabolite and inhibitor metabolite. ${F_{Sig_{trans}}}_{r,j}$ represents the regulation by a signaling or a transcription regulator for *r*th metabolite in the *j*th reaction, $W_{f}$ is the weighting factor, ${Reg}_{st}\_{act}_{j}$ and ${Reg}_{s}\_de{act}_{j}$ are the activation rate and deactivation rate by an activator *S* and an inhibitor *I* with a sensitivity of *n* and saturation constant *K,* respectively in Eqn.(5,6 and 7). $T_{tjpassive}$ and $T_{tjFaciltated}$ represent the passive and facilitated transports of *j*th metabolites to and from the tissues to blood, respectively, where $C_{bj}$ and $C_{cytj}$ are the concentrations of the metabolite in blood and tissue, respectively.

**Modifications in source models for integration:**

The metabolic network and the signaling and transcriptional regulatory network integrated and used in this study are shown in Figure S2 & S3. The source models can be referred to: hepatic metabolism (Somvanshi et al., 2016), HPA axis (Bangsgaard., 2016; Sriram et al., 2012) and inflammation model by (Bangsgaard et al., 2017; Parker et al., 2016), glucocorticoid receptor model (Rao et al., 2016), and hypoxia signaling (Cavadas et al., 2013; Nguyen et al., 2013). The metabolic model was modified to include the regulatory effects of glucocorticoids, inflammatory cytokines and HIF on metabolic enzymes in the metabolic network shown in Figure S2. The previously developed model for the HPA axis by our group was further re-parameterize to fit the data from (Son et al., 2008). The model interaction of inflammatory cytokines with HPA axis were adopted from Bangsgaard et al. (2017) with additional cytokine effects on cortisol secretion adopted from the literature (Table S2), and modeled accordingly. As we were only interested in steady state behavior of the models, in the current model analysis, circadian dynamics was not considered. While the model for inflammation was modified to include the effects from insulin and GPCR signaling pathways, the hypoxia signaling model by Cavadas et al. (2013) was updated to include the effects from inflammation and metabolism (TCA intermediates) on HIFα. The transcriptional regulation model from Somvanshi et al. (2016) was modified to include the effects from HPA axis and inflammation. The regulatory functions were applied to metabolic pathways with an assumption of parallel activation mechanism (Li et al., 2012). The additional interactions and their sources, implemented in the integrated model are reported in Table S2. Below we report only the integrated model of HPA axis, inflammation and the transcriptional compartments. The ODE model for the metabolic simulator is not explicitly reported here, however, the code can be shared on request.

**Parameter estimation**

The majority of the model parameters were maintained from the source models. The parameters encoding the regulatory interactions and the integration of models were obtained from the literature as reported in Table S2. We used a semi-empirical approach for model integration wherein we emphasized on reproducing the qualitative trends for the model state variables under different conditions as reported in corresponding literature. We fitted the Hill functions using a least squares approach to the trends reported in literature. The principle of parallel activation of pathways by regulatory influence was used while assigning regulatory effects on the metabolic pathways. We used simulated annealing in MATLAB to reparametrize the models of HPA axis and inflammation after inclusion of the regulatory functions in the integrated model (Gonzalez et al., 2007). Although the values for fold change were used from the literature, to avoid overfitting, the choices of *km* values were made on ad-hoc basis, depending on the data availability, information on the sensitivity of an interaction and the fold variation observed for a regulatory state. Wherever data was not available, a reasonable estimate of *km* values were made in terms of folds of the basal values of the states as reported in the literature. The time units for the rate parameters in our metabolic model are in *minute* and the concentrations of the metabolites are reported in millimoles (*mM*) or unless reported differently in the source models. The time scale differences (min and hrs.) in the sub models were scaled in the integrated model for the simulations. In the Table S2, we report the values of thresholds (*km* or *ki*) based on the estimates expressed in terms of the folds of the basal values for transcriptional and signaling effects, and sensitivity (*n*) assumed according to the steepness of the input-output response. The *vmax* represents the fitted estimates required to impart the observed fold change effect in the corresponding data. These rate functions are further scaled in the integrated model to obtain the resultant rates in the ODEs such that the overall state response in the model simulations match the output of the source models.

**Qualitative validation of model**

Since our goal was to use the model for hypothesis generation and perform *“what if”* kind of analysis, we were interested in a model that could reasonably reproduce the qualitative trends in the observed physiology. We mostly rely on the published models for the model performance and retain the qualitative features of the model in our integrated model after incorporation of additional interactions reported in literature. Due to the large numbers of states, a precise state to state validation was not performed and we relied on the source models for the validation. However, the model was qualitatively validated for the reported trends and observations of several experimental reports on the output response with respect to model inputs of the source models, such as, response to meal input for metabolic model, stress response of the HPA axis, and response to LPS for the inflammation model. Figure S4 shows the qualitative profiles for the model simulations for meal response, stress response and the response for infection induced by LPS impulse. The trends observed in the figure, matches the qualitative and quantitative features of the source models and the observed physiological responses in the respective cases.

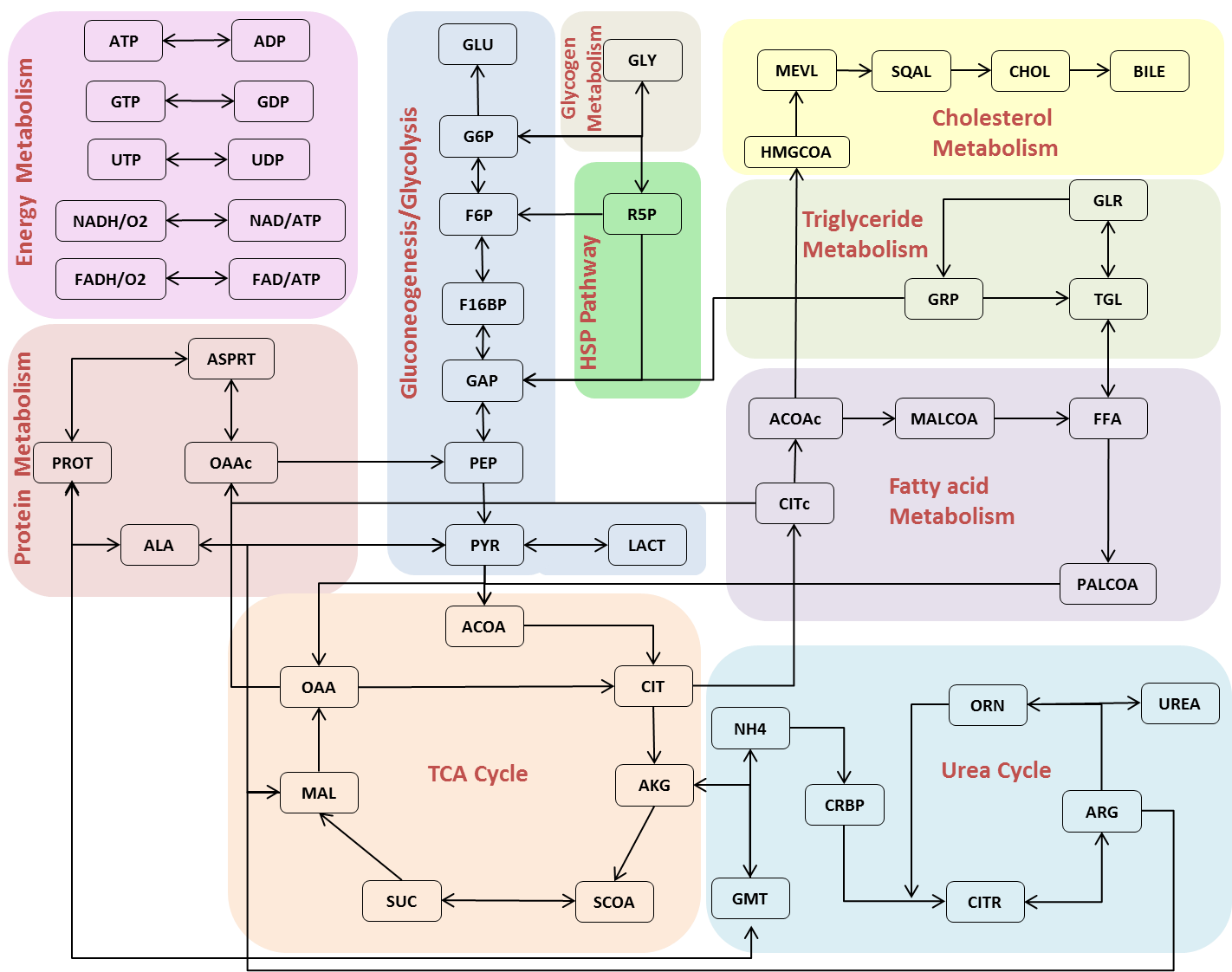

Figure S2: Metabolic pathways coded in the mathematical model for metabolic simulator. The model constitutes glycolysis, gluconeogenesis, TCA cycle, urea cycle, lipogenesis, amino acid metabolism, hexose amine pathway, pentose phosphate pathway, oxidative phosphorylation and cholesterol pathway, plasma metabolite fluxes. This model is further integrated with regulations from insulin signing, glucagon signaling, transcription regulation, inflammation, hypoxia signaling and HPA axis.

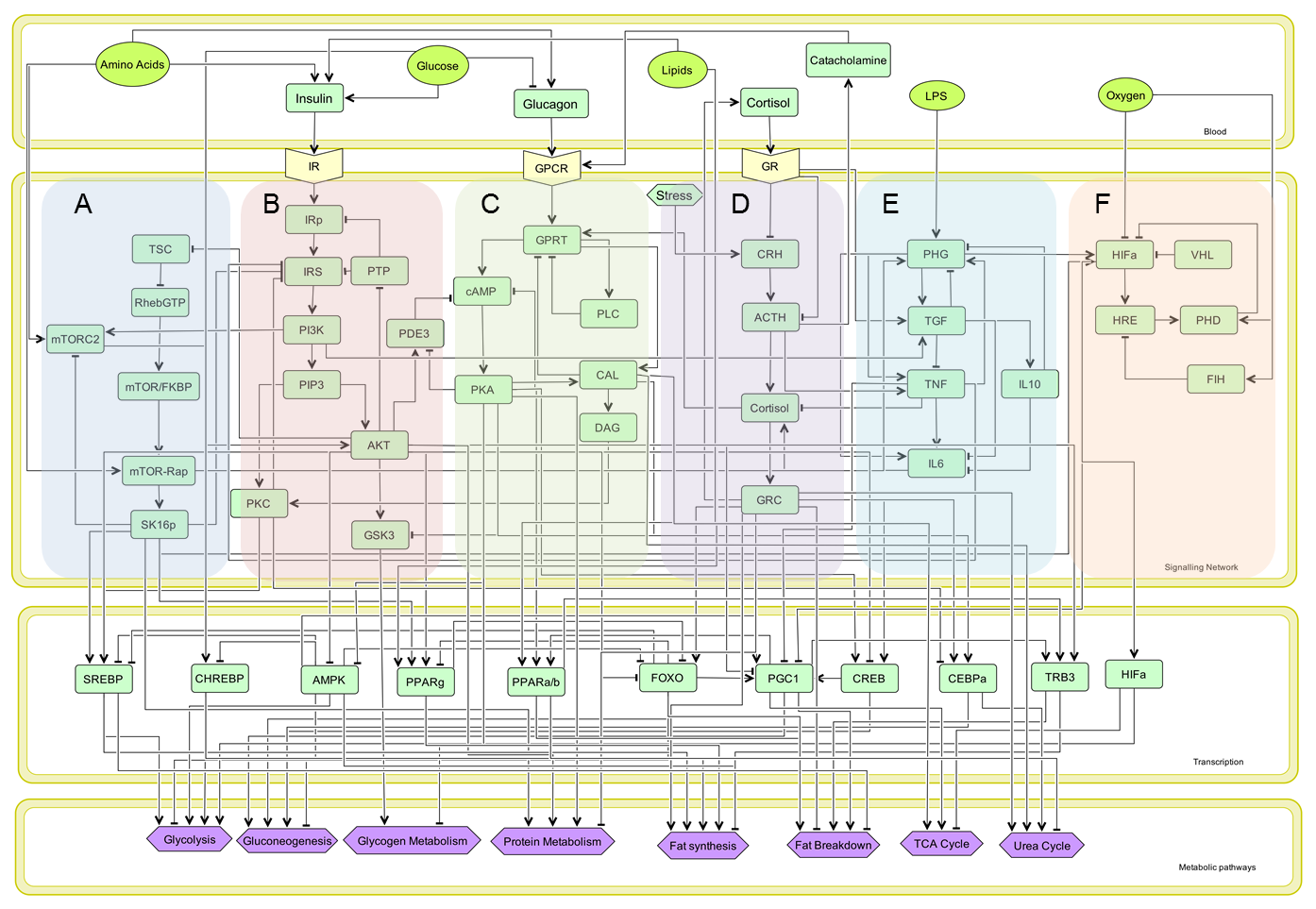

Figure S3: Regulatory network representing of six signaling pathways, ten transcription factors and the metabolic process that are regulated by the network. The network comprises of (A) mTOR pathway, (B) insulin signaling pathway, (C) GPCR signaling pathway, (D) HPA axis, (E) inflammatory signaling and (F) hypoxia signaling. These signaling pathways interact to activate downstream transcription regulatory network as shown in the transcription factor compartment. The signaling and transcription network collectively influence metabolic processes as represented in the metabolic pathways compartment.

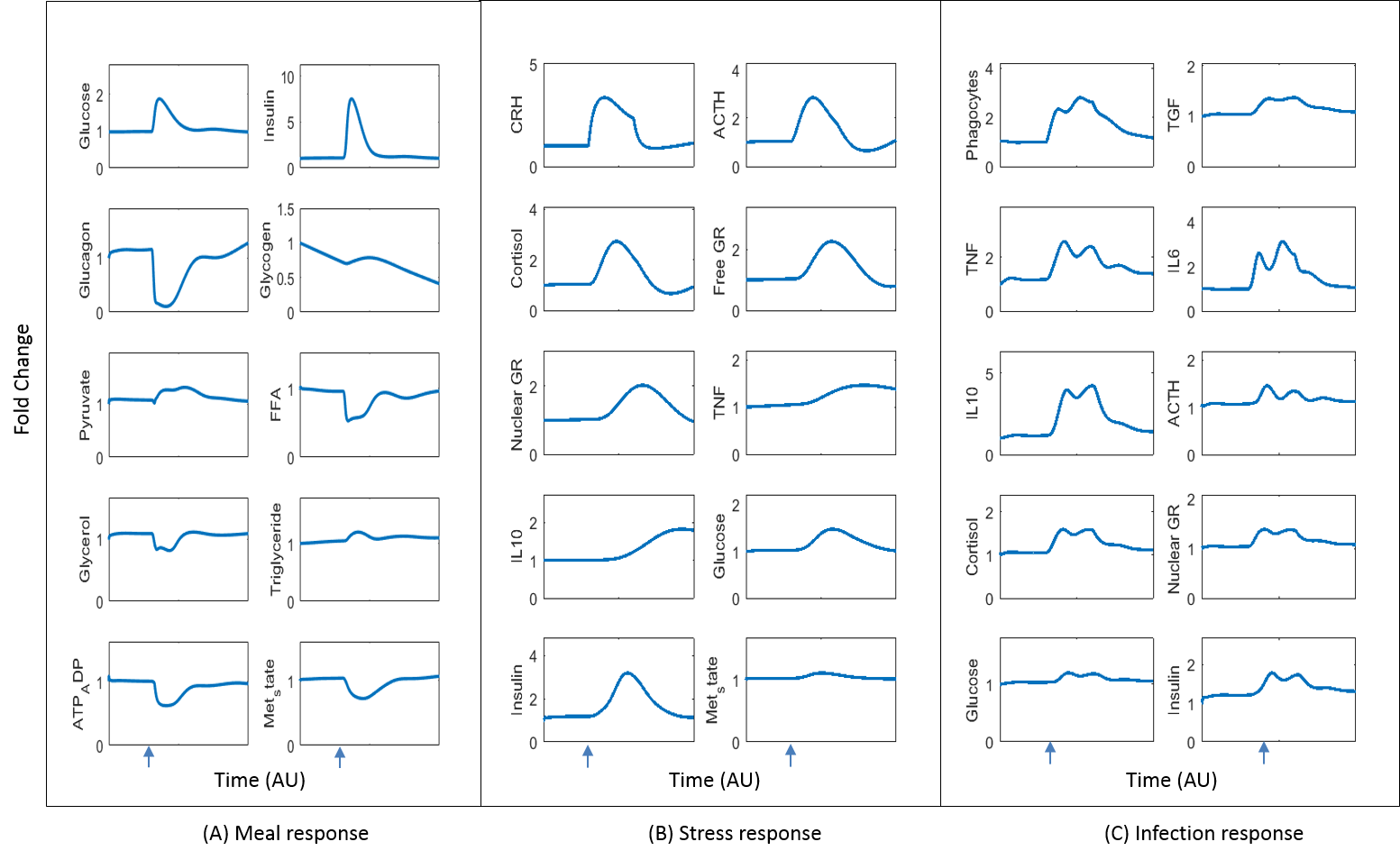

Figure S4: Dynamics of fold change (normalized response) in the state variables of the model for input impulses of CHO meal, stress and infection (lipopolysaccharides) starting at the time represented by the arrow on x-axis. (A) Meal response: The model was simulated for 75 gm of carbohydrate meal impulse and the corresponding dynamics of the fold change in the metabolic states are shown. The trends ae reproduced as per the source models and observed profiles in experimental data (DallaMan et al., 2007; Galgani et al., 2006; Krssak et al., 2004). (B) Stress response: The model was simulated for a stress impulse (5 fold change in the stress parameter). The corresponding dynamics of fold change in HPA axis and cytokine state variables are shown. The associated changes in metabolic variables for insulin, glucose and metabolic state are also reported. The trends match the source models and response reported in (Kudielka et al., 2004; Li et al., 2013; Maes et al., 1998). (C) Response to infection/LPS: The model was simulated for an impulse of LPS (10e-5 AU). The dynamics of the corresponding fold change in pro-inflammatory and anti-inflammatory cytokines are shown, along with the HPA axis and metabolic components. The model reproduces trends from the source models and observations reported in (Copeland et al., 2005; Grinevich et al., 2001; Kariagina et al., 2004; Matsuzaki et al., 2001; Nguyen et al., 2014).

Table S2: Regulatory interactions implemented in the integration of sub-models for signaling and transcription regulation of metabolism, HPA axis, inflammation and hypoxia.

| **Regulatory Interactions** | **References for regulatory interaction** | **Experimental organism** | **KO mutants/ dose response** | **Fold change/ +/-** | **Parameter estimates for Hill functions** | | | |
| --- | --- | --- | --- | --- | --- | --- | --- | --- |
|  |  |  |  |  | *Vmax*  *AU* | *km/ki*  *AU* | | *n* |
| **Metabolic regulation by glucocorticoids** | | | | | | | | |
| GR mediated effects on gluconeogenesis (Pck1 and G6pc) | (Cassuto et al., 2005) | Human HepG2 cells, | Gene expression | ~5  (PEPCK) |  | |  |  |
|  | (Frijters et al., 2010) | Mice | Prednisone treated | ~1.5-4  (several genes) |  |  |  |  |
| Activation of lipogenesis | (Gathercole et al., 2011) | Human Chub-S7 cell culture | DEX treatment | ~2-4  (FAS and ACC) | 5 | | 20 | 2 |
|  | (Lemke et al., 2008) | Mice liver | GR mutants | ~1.2-1.5 (TG) |  |  |  |  |
| Inhibition of β-oxidation | (Samra et al., 1998) | Human | Hydrocortisone infusion | ~2 | 2 | | 10 | 3 |
|  | (Letteron et al., 1997) | Mice | DEX treatment | ~1.25~2  (enzymes) |  |  |  |  |
| Activation of proteolysis and plasma amino acids | (Wang et al., 2016) | Murine Myoblasts | DEX treatment | ~1.25-2 | 3 | | 20 | 1 |
|  | (Löfberg et al., 2002) | Human | Prednisolone treated | ~1.25-2  (several plasma amino acids) |  |  |  |  |
| Activation of urea cycle | (Okun et al., 2015) | Human serum and Mouse | Liver GR mutant | ~1.25-2.5  (Arginine/ Ornithine / urea) | 3 | | 20 | 1 |
|  | (Haggerty et al., 1982) | Rat hepatocytes | Hydrocortisone treated | ~10 (arginase activity) |  |  |  |  |
| Inhibition of insulin secretion | (Jeong et al., 2001) | Rat pancreatic Islets | DEX treated | ~3 | 0.25 | | 20 | 2 |
|  | (Lambillotte et al., 1997) | Mouse islets | DEX treated | ~3 |  |  |  |  |
| **Transcription regulation by glucocorticoids** | | | | | | | | |
| Activation of CREB | (Nagao et al., 2012) | Human pre B cell lines | DEX treated  (1μm for 72 hrs) | ~4 | 5 | | 20 | 2 |
|  | (Imai et al., 1993) | Rat H4IIE Hepatocytes | DEX treated  (0.5μm ,4 hrs) | ~3 |  |  |  |  |
|  | (Rahnert et al., 2016) | Rat L6 myoblasts | DEX treated  (0.1μm, 48 hrs) | ~1.5 |  |  |  |  |
| Activation of FOXO | (Lützner et al., 2012) | Mice | DEX treated | ~2 | 5 | | 20 | 2 |
|  | (Zhao et al., 2009) | Mice C2C12 cells | Mutant | ~1.5 |  |  |  |  |
|  | (Kaiser et al., 2013) | Mice | DEX treated  (0.1μm, 24 h) | ~2.5 |  |  |  |  |
| Activation of CEBP | (Cram et al., 1998)  (Ramos et al., 1996) | Rat hepatocytes | DEX treated | ~2 | 5 | | 20 | 2 |
|  | (Matsuno et al., 1996) | Rat hepatocytes | DEX treated | ~2 |  |  |  |  |
| Activation of PGC1 | (Kok et al., 2013) | Rat liver and myocytes | DEX treated , Gene expression | ~2 | 5 | | 20 | 2 |
|  | (Frijters et al., 2010) | Mice | Prednisone treated | ~3 |  |  |  |  |
|  | (Herzig et al., 2001) | Human HepG2 hepatocytes | DEX treated  (0.1μm) | ~4 |  |  |  |  |
| **Signaling regulation by glucocorticoids** | | | | | | | | |
| Activation of adrenergic receptors/GPCR | (Haigh et al., 1990) | Rat | DEX treated  Adrenalectomy | ~2  (Gi/Gs protein) | 3 | | 20 | 2 |
|  | (Jazayeri and Meyer, 1988) | Rat | DEX treated  (0.1μm, 24 hrs) | ~2 |  |  |  |  |
| Inhibition of mTORC1 | (Shimizu et al., 2011) | Rat | Gene Expression | ~1.5-2 | 1.1 | | 30 | 2 |
|  | (Wang et al., 2006) | Rat | DEX treated | ~1.5-2 |  |  |  |  |
|  | (Kim et al., 2016) | Mice C2C12 cell culture | DEX/Cortisol  Treated | ~1.25-1.5 |  |  |  |  |
| Inhibition of IRS/PI3k | (Kuo et al., 2012) | Mice C57BL/6, C2C12 cell culture | Gene expression | ~1.5-2 | 1.25 | | 20 | 2 |
|  | (Saad et al., 1993) | Rat Liver/ Muscle | DEX treatment | ~1.2-1.5 |  |  |  |  |
|  | (Giorgino et al., 1993) | Rat | Cortisone treatment | ~2 |  |  |  |  |
| **Regulation by TNF and IL6/ Inflammation** | | | | | | | | |
| Inhibition of Insulin signaling by TNF) (tyrosine phosphorylation of IRS) | (Hotamisligil et al., 1996) | Murine 3T3-L1 cells | TNF treated | ~4 | 5 | | 100 | 4 |
|  | (Klover et al., 2003) | Mice Liver | IL6 treated | ~5 |  |  |  |  |
| TNF and IL6 inhibits PGC1α | (Li et al., 2014) | LO2 cell culture and Rat Liver | Mutants/Gene expression | ~5 | 5 | | 100 | 2 |
|  | (Sun Kim et al., 2007) | Human Hep3B cells and mice liver | LPS treated | ~2 |  |  |  |  |
| Inflammatory cytokines activates HPA axis components | (Bethin et al., 2000)  (Bornstein et al., 2004) | Mice/  Human adrenal cells | IL6 stimulation/ mutant | ~2-3 (ACTH/ Cortisol) | Reported and modified as per source models  (Bangsgaard., 2016) | | | |
|  | (Kanczkowski et al., 2013) | Mice | LPS treated | ~2-3  (ACTH/ Cortisol) |  |  |  |  |
| Inflammatory cytokine /phagocytes activates HIFα | (Frede et al., 2006) | Human Cell lines: monocyte THP-1 | LPS treated | ~2 | 5 | | 1.00E+05 | 2 |
|  | (van Uden et al., 2008) | Human HEK293 cells | NF-kb treated | ~2-3 |  |  |  |  |
| Inflammation inhibits oxidative phosphorylation | (Samavati et al., 2008) | Mouse Liver tissue, Murine H2.35 cells | TNF treated | ~2  (CcO activity) | 1 | | 50 | 2 |
|  | (Duvigneau et al., 2007) | Rats livers | LPS treated | ~3-4  (ATP depletion) |  |  |  |  |
| Inflammation induces ROS production/ATP depletion | (Kastl et al., 2014) | Human hepatic Hepa1-6, Murine hepatocytes | TNF treated | ~2-3 |  | |  |  |
| **Regulation by HIFα/ hypoxia** | | | | | | | | |
| Alpha-ketoglutarate inhibits HIFα by activation of PHD | (Koivunen et al., 2007) | Human HEK293 cells | Hypoxia induced | ~2-3 | 2.5 | | 0.4 | 2 |
|  | (MacKenzie et al., 2007) | Human HEK293 cells | Plasmid infected | Dose dependent |  |  |  |  |
| Succinate activates HIFα by inhibiting PHD | (Selak et al., 2005) | Human HEK293 cells | Plasmid infected | ~2-3 | 2.5 | | 1.6 | 2 |
|  | (Koivunen et al., 2007) | Human HEK293 cells | Hypoxia induced | ~2-3 |  |  |  |  |
| HIFα upregulates glycolysis | (Semenza et al., 1994)  (Corcoran and O’Neill, 2016) | Human Hela and Hep3B cells | Hypoxia induced  Gene expn. | ~3-8  (ALDA,PGK1, enolase, PFK) | 5 | | 0.25 | 2 |
|  | (Obach et al., 2004) | Human T98G and U-87 cell lines | Hypoxia treated | ~10  (PFK2 and GLUT1) |  |  |  |  |
| HIFα downregulates mitochondrial biogenesis transcription/PGC1 | (Zhang et al., 2007) | Human RCC4 cells | Mutant | ~2  (C-MYC) | 1 | | 0.5 | 2 |
| HIFα inhibits Pyruvate dehydrogenase (activation of PDK) | (Lu et al., 2008) | Human Hela cells | Gene expression | ~3-4  (PDK3) | 1 | | 0.25 | 2 |
|  | (Kim et al., 2006) | Mice: Mouse embryonic fibroblasts | Gene expression | ~3-4  (PDK1) |  |  |  |  |
| HIFα activates lactate dehydrogenase | (Firth et al., 1995) | Human Hela cells | Gene expression | ~2 | 3 | | 0.25 | 3 |
|  | (Semenza et al., 1996) | Human Hep3B cells | Mutant | ~4 |  |  |  |  |
| HIFα inhibits oxidative phosphorylation/ Respiratory chain complex | (Chen et al., 2010) | Human HCT116 cells | Gene expression | ~2  (COX1) | 1 | | 0.25 | 2 |
|  | (Fukuda et al., 2007) (Semenza, 2007) | Human HeLa, 293T cells | Gene expression | ~5  (COX4-1) |  |  |  |  |
| HIFα activates inflammatory cytokines/phagocytosis | (Anand et al., 2007) | Mice: Murine macrophages | Hypoxia treated | ~2 | 5 | | 0.5 | 2 |
|  | (Kuhlicke et al., 2007) | Human cells:HMEC-1 | Hypoxia treated | ~3 |  |  |  |  |
| HIFα activates ROS generation | (Bhogal et al., 2010) | Human hepatocytes | Hypoxia treated | ~4 | 1 | | 0.5 | 1 |
| HIFα induce triglyceride accumulation | (Nath et al., 2011) | Mice | Mutant | ~2 | 3 | | 0.25 | 2 |
|  | (Rankin et al., 2009) | Mice | Mutant | ~2 |  |  |  |  |
| HIFα inhibits beta oxidation and lipogenesis | (Rankin et al., 2009) | Mice | Mutant | ~2.5-4 | 1 | | 0.5 | 2 |
|  | (Huss et al., 2001) | Mice | Mutant | ~2 |  |  |  |  |
|  | (Huang et al., 2014) | Human Hep3B cells | Gene expression | ~2 |  |  |  |  |

**Integrated model for HPA axis, GR signaling and inflammation**

Table S3: Parameters for integrated HPA axis-inflammation model

| **Parameter** | **Value** | **Units** | **Parameter** | **Value** | **Units** | **Parameter** | **Value** | **Units** |
| --- | --- | --- | --- | --- | --- | --- | --- | --- |
| $n$ | 1 | *-* | $kphg$ | 4.9956E7 | *kg/hr/pg* | $xil6il10$ | 1.1818 | *pg/ml* |
| $Ki_{n1}$ | 1.2 | *nM/ml* | $kptnf$ | 12.94907 | *-* | $dil6$ | 0.43605 | */h* |
| $K_{strs}$ | 1 | *µg/dL/min* | $xnTNF$ | 1693.9509 | *pg/ml* | $il6b$ | 0.25 | *pg/ml. h* |
| $n3$ | 2 | *-* | $xTGF$ | 0.07212 | *pg/ml* | $q1$ | 0.5 | *ml/pg.h* |
| $Vs3$ | 0.032 | */ min* | $xIL10$ | 147.68 | *pg/ml* | $q2$ | 625 | *µg/dL* |
| $Kp2$ | 0.41 | */ min* | $dpg$ | 0.144 | */h* | $kmpi3k$ | 1.4 | *AU* |
| $Vs4$ | 0.016 | */ min* | $ktgf$ | 0.15625E-8 | *ml/pg.U.h* | $kmcreb$ | 0.75 | *AU* |
| $Vs5$ | 0.0266 | */ min* | $dtgf$ | 0.03177 | */h* | $fpi3k$ | 5 | *-* |
| $kp4$ | 45E-4 | */min* | $ktnf$ | 25.5194 | *pg/ml.h* | $fcreb$ | 5 | *-* |
| $q3$ | 2.8014 | *pg/mL.min* | $TNFn$ | 550E4 | *U* | $wt$ | 0.66 | *-* |
| $q4$ | 11.2 | *pg/mL.min* | $xtntg$ | 0.1589*1.25 | *pg/ml* | $fmtor$ | 5 | *-* |
| $q5$ | 40 | *pg/mL* | $ktnff$ | 3.5514E4 | *pg/ml.h* | $kIL10$ | 267480 | *pg/ml. h* |
| $q6$ | 7 | *pg/mL* | $kmtor$ | 2 | *-* | $kil6$ | 5E2 | *pg/ml. h* |
| $q8$ | 40 | *pg/mL* | $dtnf$ | 0.0307 | *ml/pg. h* | $xil6$ | 33E4 | *pg/ml* |
| $ksynt_{rm}$ | 3.625 | */h* | $sil$ | 1187.2 | *pg/ml.h* | $kil6tnf$ | 4.4651 | *pg/h.ml.U* |
| $nkm_{Grn}$ | 26 | *nM/mg protein* | $xil$ | 8.0506E7 | *U* | $xil6tnf$ | 1211.3 | *pg/ml* |
| $kdeg_{rm}$ | 0.1124 | */h* | $dil$ | 98.932 | */h* | $It$ | 1E-6 | *U* |
| $ksynt_{rp}$ | 1.2 | */h* | $kiltg$ | 43875 | *pg/ml.h* | $kmtnf$ | 100 | *pg/ml* |
| $vp\_rp$ | 0.0279 | */h* | $xiltg$ | 0.38*2 | *pg/ml* | $dlp$ | 1.35E-7 | */h.U* |
| $kon$ | 0.00329 | *nM/h* | $vtnf$ | 1 | *nM/h* | $nf1$ | 4 | *-* |
| $deg\_rp$ | 0.0572 | */h* | $kp_{tnf}$ | 10 | *pg/ml* | $nf4$ | 3 | *-* |
| $krt$ | 0.63 | */h* | $m$ | 4 | *-* | $n4$ | 2 | *-* |
| $kre$ | 0.57 | */h* | $nx$ | 1 | *-* | $nf2$ | 6 | *-* |
| $taup$ | 0.1401 | *h* | $nf3$ | 2 | *-* |  |  |  |

**Model equations for integrated HPA axis-inflammation model**

$$\dot{\boldsymbol{Crh}}=K_{strs}*\left( \frac{Ki_{n1}^{n}}{Ki_{n1}^{n}+{GRn}^{n}} \right)+ q3*\left( \frac{TNF}{q8+ TNF} \right)-Vs3*Crh$$

$$\dot{\boldsymbol{Acth}}= Kp2*Crh*\left( \frac{Ki_{n1}^{n}}{Ki_{n1}^{n}+{GRn}^{n}} \right)*\left( 1+q4*\left( \frac{TNF^{n3}}{q5^{n3}+ TNF^{n3}} \right) \right)-Vs4*Acth$$

$$\dot{\boldsymbol{Cort}}= kp4*Acth*\left( 1+\left( \frac{TNF}{1+q6*\left( TGF+IL6 \right)} \right) \right)-Vs5*Cort$$

$$\dot{\boldsymbol{Cortp}}=\left( \frac{1}{taup} \right)*\left( Cort-Cortp \right)$$

$$\dot{\boldsymbol{GRmrna}}= ksynt_{rm}*\left( 1 - \left( \frac{{GRn}^{nx}}{nkm_{Grn}^{nx}+{GRn}^{nx}} \right) \right)+vtnf*\left( \frac{TNF^{m}}{TNF^{m}+kp_{tnf}^{m}} \right)- kdeg_{rm}*GRmrna$$

$$\dot{\boldsymbol{GRprt}}= ksynt_{rp}*GRmrna+ vp_{rp}*GRn - kon*Cortp*GRprt- kdeg_{rp}*GRprt$$

$$\dot{\boldsymbol{GRf}}= kon*Cortp*GRprt- krt*GRf$$

$$\dot{\boldsymbol{GRn}}= krt*GRf -kre*GRn$$

$$PI3K_{eff}=wt*\left( 1+\left( fpi3k*\left( \frac{{PI3k}^{n4}}{{PI3k}^{n4}+{kmpi3k}^{n4}} \right) \right) \right)$$

$$Creb_{eff}=fcreb*\left( \frac{CREB^{n4}}{CREB^{n4}+{kmcreb}^{n4}} \right)$$

$$mTOR_{eff}=wt*\left( 1+ fmtor*\left( \frac{{mTOR}_{Raptor}^{n4}}{{kmtor}^{n4}+{mTOR}_{Raptor}^{n4}} \right) \right)$$

$$\dot{\boldsymbol{LPS}}=It -dlp*LPS*Phg$$

$$\dot{\boldsymbol{Phg}}= kphg*\left( \left( 1+ kptnf*\left( mTOR_{eff} \right)* \left( \frac{TNF}{xnTNF+TNF} \right) \right)* \left( \frac{xTGF}{xTGF+ TGF} \right)* \left( \frac{xIL10}{xIL10+IL10} \right) \right)*LPS- dpg*Phg$$

$$\dot{\boldsymbol{TGF}}= ktgf* Phg*\left( PI3K_{eff} \right) + \left( q1*\frac{{GRn}^{nx}}{\left( q2 \right)^{nx}+{GRn}^{nx}} \right)- dtgf*TGF;$$

$$KTGF=xtntg*wt*\left( 1+Creb_{eff} \right)$$

$$\dot{\boldsymbol{TNF}}= \left( \frac{Phg}{kTNFn+Phg} \right)* \left( \frac{KTGF^{nf1}}{KTGF^{nf1}+ TGF^{nf1}} \right)*\left( ktnf+ \left( ktnff*\frac{TNF}{xTNFf+\left( TNF \right)} \right) \right)- dtnf*TNF^{2}$$

$$\dot{\boldsymbol{IL}\boldsymbol{10}}= sil+ \left( kIL10*\frac{Phg^{nf4}}{xil^{nf4}+Phg^{nf4}} \right)+ \left( kiltg*\frac{TGF^{nf2}}{xiltg^{nf2}+ TGF^{nf2}} \right)- dil*IL10$$

$$KIL10=xil6il10*wt*\left( 1+Creb_{eff} \right);$$

$$\dot{\boldsymbol{IL}\boldsymbol{6}}=il6b+kil6*\left( \frac{Phg^{nf3}}{xil6^{nf3}+Phg^{nf3}} \right)*\left( 1+kil6tnf*\left( \frac{TNF+IL6}{xil6tnf+TNF+IL6} \right) \right)*\left( \frac{KIL10}{KIL10+IL10} \right)-dil6*IL6$$

**Transcriptional regulation network**

Table S4: Parameters for transcriptional regulatory network

| **Parameter** | **Value** | **Units** | **Parameter** | **Value** | **Units** | **Parameter** | **Value** | **Units** |
| --- | --- | --- | --- | --- | --- | --- | --- | --- |
| $nfgcr$ | 5 | *AU* | $K_{ATP}$ | 2.8 | *mM* | $fFFA1$ | 5 | *-* |
| $s2$ | 2 | *-* | $AMPKt$ | 1 | *AU* | $kmffa1$ | 1.15 | *mM* |
| $Kmgcr$ | 20 | *µg/dL* | $kcb1$ | 0.003 | *AU/min* | $fpgc$ | 5 | *-* |
| $ksr1$ | 3E-3 | *AU/min* | $kcb2$ | 0.02 | */min* | $kmpgc$ | 2 | *AU* |
| $ksr2$ | 5E-3 | */min* | $kcb0$ | 0.016 | *AU /min* | $fpka2$ | 10 | *-* |
| $ksr0$ | 8E-3 | *AU /min* | $kpc1$ | 8E-4 | *AU /min* | $kmpka2$ | 3 | *AU* |
| $kprg1$ | 9E-3 | *AU /min* | $kpc2$ | 0.02 | */min* | $fakt2$ | 3 | *-* |
| $kprg2$ | 0.02 | */min* | $kpc0$ | 0.016 | *AU /min* | $kmakt2$ | 2 | *AU* |
| $kprg0$ | 0.03 | *AU /min* | $ktr1$ | 1.65E-3 | *AU /min* | $fglu$ | 5 | *-* |
| $kprg3$ | 5E-3 | *AU /min* | $ktr2$ | 0.02 | */min* | $kmglu$ | 10 | *mM* |
| $kpra1$ | 2.5E-3 | *AU /min* | $ktr0$ | 0.014 | *AU /min* | $fpka3$ | 5 | *-* |
| $kpra2$ | 0.02 | */min* | $fs6k$ | 0.5 | *-* | $kmpka3$ | 2 | *AU* |
| $kpra0$ | 0.012 | *AU /min* | $kms6k$ | 1 | *AU* | $fampk1$ | 1.25 | *-* |
| $kpra3$ | 5E-3 | *AU /min* | $fcAMP$ | 2 | *-* | $kiampk$ | 0.4 | *AU* |
| $khr1$ | 0.02 | *AU /min* | $kicAMP$ | 3.2E-6 | *mM* | $fglnac$ | 5 | *-* |
| $khr2$ | 0.02 | */min* | $fAMPK$ | 1.25 | *-* | $kmglnac$ | 0.05 | *mM* |
| $khr0$ | 2.5E-3 | *AU /min* | $kiAMPK$ | 0.5 | *AU* | $fakt3$ | 3 | *-* |
| $kcr1$ | 5E-3 | *AU /min* | $kiFOXO$ | 0.5 | *AU* | $kmakt3$ | 2.2 | *AU* |
| $kcr2$ | 0.03 | */min* | $fins$ | 5 | *-* | $fpparg$ | 1.25 | *-* |
| $CREBt$ | 1 | *AU* | $kins$ | 2 | *-* | $kmpparg$ | 2 | *AU* |
| $kfr11$ | 2.5346E-3 | *AU/min* | $kmffa$ | 1.7 | *mM* | $fakt4$ | 5 | *-* |
| $kfr0$ | 1.215E-3 | *AU/min* | $fFFA$ | 12.5 | *-* | $kmakt4$ | 2 | *AU* |
| $kfr2$ | 0.0215 | */min* | $fakt1$ | 5 | *-* | $fpka4$ | 5 | *-* |
| $kam1$ | 1 | *AU* | $kmakt1$ | 2 | *AU* | $kmpka4$ | 2 | *AU* |
| $Kam2$ | 1.1925 | *AU* | $fFoxo$ | 1.25 | *-* | $fpkc1$ | 3 | *-* |
| $K_{AMP}$ | 0.16 | *mM* | $fpka$ | 5 | *-* | $kmpkc$ | 3 | *AU* |
| $fppar$ | 5 | *-* | $kmcreb1$ | 0.5 | *AU* | $fcAMP1$ | 10 | *-* |
| $kmppar$ | 2 | *AU* | $fampk1$ | 5 | *-* | $kmcAMP1$ | 3 | *AU* |
| $fpgc1$ | 5 | *-* | $ftnfil6$ | 5 | *-* | $ffoxo1$ | 5 | *-* |
| $kmpgc1$ | 2 | *AU* | $kmtnfil6$ | 100 | *pg/ml* | $kmfoxo1$ | 0.5 | *AU* |
| ${PKA}_{b}$ | 9E-6 | *mM* | $fpi3k$ | 4 | *-* | $fakt5$ | 3 | *-* |
| ${cAMP}_{b}$ | 3.2E-6 | *mM* | $kmpi3k$ | 1.25 | *AU* | $kmakt5$ | 2.25 | *AU* |
| $kmpka$ | 2 | *AU* | $fpkc2$ | 2 | *-* | $kmhif$ | 0.5 | *AU* |
| $s3$ | 3.5 | *-* | $kmpkc2$ | 2 | *AU* | $fcreb1$ | 5 | *-* |

**Model equations for transcriptional regulatory network**

$$GCR_{Ptv}=fgcr* \left( \frac{{GRn}^{s2}}{Kmgcr^{s2}+{GRn}^{s2}} \right)$$

$$Inseff=\frac{AKT+PKC}{{AKT}_{b}+{PKC}_{b}}$$

$$S6k_{Ptv}=fs6k*\left( \frac{{S6K1p}^{s2}}{{S6K1p}^{s2}+{kms6k}^{s2}} \right)$$

$$cAMP_{Ntv}=fcAMP*\left( \frac{{kicAMP}^{s2}}{{kicAMP}^{s2}+cAMP^{s2}} \right)$$

$$AMPK_{Ntv}=fAMPK*\left( \frac{{kiAMPK}^{s2}}{{kiAMPK}^{s2}+AMPK_{Eff}^{s2}} \right)$$

$$FO{XO}_{Ntv_{SREBP}}=\left( \frac{{kiFOXO}^{s2}}{FOXO^{s2}+{kiFOXO}^{s2}} \right)$$

$$AKT_{PKC_{Ptv_{SREBP}}}=fins*\left( \frac{Inseff^{s2}}{Inseff^{s2}+{kins}^{s2}} \right)$$

$$\dot{\boldsymbol{SREBP}}=ksr0*S6k_{Ptv}+\left( ksr1*cAMP_{Ntv}*AMPK_{Ntv}*FO{XO}_{Ntv_{SREBP}}*AKT_{PKC_{Ptv_{SREBP}}} \right)-ksr2*SREBP$$

$$FFA_{Ptv}=fFFA*\left( \frac{{FFA}^{s3}}{{FFA}^{s3}+{kmffa}^{s3}} \right)$$

$$AKT_{Ptv_{PPARg}}=fakt1*\left( \frac{{Akt}^{s2}}{{Akt}^{s2}+{kmakt1}^{2s2}} \right)$$

$$FO{XO}_{Ntv_{PPARg}}=fFoxo*\left( \frac{{kiFOXO}^{s2}}{FOXO^{2s2}+{kiFOXO}^{s2}} \right)$$

$$HIF_{Ntv_{PPARg}}=\left( \frac{{kmhif}^{s2}}{{kmhif}^{s2}+HIF\alpha^{s2}} \right)$$

$$\dot{\boldsymbol{PPARg}}=kprg0*S6k_{Ptv}+\left( kprg1*AKT_{Ptv_{PPARg}}*FO{XO}_{Ntv_{PPARg}}*HIF_{Ntv_{PPARg}} \right)+\left( kprg3*FFA_{Ptv} \right)-kprg2*PPARg$$

$$PKA_{Ptv_{PPARa}}=fpka*\left( \frac{{PKA}^{s2}}{{PKA}^{s2}+{kmpka}^{s2}} \right)$$

$$FFA_{Ptv}=fFFA1*\left( \frac{FFA^{s2}}{FFA^{s2}+{kmffa1}^{s2}} \right)$$

$$PGC_{Ptv_{PPARab}}=fpgc*\left( \frac{PGC1^{s2}}{PGC1^{s2}+{kmpgc}^{s2}} \right)$$

$$HIF_{Ntv_{PPARab}}=\left( \frac{{kmhif}^{s2}}{{kmhif}^{s2}+HIF\alpha^{s2}} \right)$$

$\dot{\boldsymbol{PPARab}}$ $=kpra0+\left( kpra1*\left( PKA_{Ptv_{PPARa}}+PGC_{Ptv_{PPARab}} \right)*HIF_{Ntv_{PPARab}} \right)+kpra3*FFA_{Ptv}-kpra2*PPARab$

$$PKA_{Ptv_{CREB}}=fpka2*\left( \frac{{PKA}^{s2}}{{PKA}^{s2}+{kmpka2}^{s2}} \right)$$

$$AKT_{Ntv_{CREB}}=fakt2*\left( \frac{Akt}{kmakt2+Akt} \right)$$

$$\dot{\boldsymbol{CREB}}=\left( kcr1*\left( PKA_{Ptv_{CREB}}+GCR_{Ptv} \right)*\left( CREBt-CREB \right) \right)-kcr2*\left( CREB*AKT_{Ntv_{CREB}} \right);$$

$Glu_{Ptv}=1+fglu*\left( \frac{{Glu}^{5}}{{Glu}^{5}+{kmglu}^{5}} \right)$ ; $PKA=\frac{PKA}{{PKA}_{b}}$

$$PKA_{Ntv}=fpka3*\left( \frac{{PKA}^{s2}}{{kmpka3}^{s2}+{PKA}^{s2}} \right)$$

$$AMPK_{Ntv}=fampk1*\left( \frac{{kiampk}^{s2}}{{kiampk}^{s2}+AMPK_{Eff}^{s2}} \right)$$

$$\dot{\boldsymbol{CHREBp}}=khr0+\left( khr1*Glu_{Ptv}*AMPK_{Ntv} \right)-khr2*\left( CHREBp*PKA_{Ntv} \right);$$

$$GlNAc_{Ptv_{FOXO}}=fglnac*\left( \frac{{GlNAc}^{s2}}{{GlNAc}^{s2}+{kmglnac}^{s2}} \right)$$

$$AMPK_{Ntv}=fampk1*\left( \frac{{kiampk}^{s2}}{{kiampk}^{s2}+AMPK_{Eff}^{s2}} \right)$$

$$AKT_{Ntv_{FOXO}}=fakt3*\left( \frac{Akt}{Akt+kmakt3} \right)$$

$$PPARg_{Ntv_{FOXO}}=fpparg* \left( \frac{{kmpparg}^{s2}}{PPARg^{s2}+{kmpparg}^{s2}} \right)$$

$$\dot{\boldsymbol{FOXO}}=kfr0+\left( kfr11*AMPK_{Ntv}*PPARg_{Ntv_{FOXO}}*\left( GlNAc_{Ptv_{FOXO}}+GCR_{Ptv} \right) \right)-kfr2*\left( FOXO*AKT_{Ntv_{FOXO}} \right)$$

$$AMP_{ATP_{AMPK}}=2*\left( \frac{\frac{AMP}{ATP}}{\left( \frac{K_{AMP}}{K_{ATP}} \right)+\left( \frac{AMP}{ATP} \right)} \right)$$

$$AKT_{Ntv_{AMPK}}=fakt4*\left( \frac{{Akt}^{s2}}{{Akt}^{s2}+{kmakt4}^{s2}} \right)$$

$$PKA_{Ntv_{AMPK}}=fpka4*\left( \frac{{PKA}^{s2}}{{kmpka4}^{s2}+{PKA}^{s2}} \right)$$

$$\dot{\boldsymbol{AMPK}}=\left( kam1*\left( AMP_{ATP_{AMPK}} \right)*\left( AMPKt-AMPK \right) \right)-Kam2*\left( AMPK \right)*\left( AKT_{Ntv_{AMPK}}+PKA_{Ntv_{AMPK}} \right)$$

$PKC_{Ntv_{CEBPa}}=fpkc1*\left( \frac{PKC}{PKC+kmpkc} \right)$; $cAMP=\frac{cAMP}{{cAMP}_{b}}$

$$cAMP_{Ptv_{CEBPa}}=fcAMP1*\left( \frac{{cAMP}^{s2}}{c{AMP}^{s2}+{kmcAMP1}^{s2}} \right)$$

$$\dot{\boldsymbol{CEBPa}}=kcb0+\left( kcb1*\left( cAMP_{Ptv_{CEBPa}}+GCR_{Ptv} \right) \right)-kcb2*\left( CEBPa*PKC_{Ntv_{CEBPa}} \right)$$

$$NAD_{Ptv_{PGC}}=2*\left( \frac{\frac{NAD}{NADH}}{\frac{NAD}{NADH}+\frac{K_{NAD}}{K_{NADH}}} \right)$$

$$FOXO_{Ptv_{PGC}}=ffoxo1*\left( \frac{FOXO^{s2}}{FOXO^{s2}+{kmfoxo1}^{s2}} \right)$$

$$AKT_{Ntv_{PGC}}=fakt5*\left( \frac{Akt}{Akt+kmakt5} \right)$$

$$HIF_{Ntv_{PGC}}=\frac{{kmhif}^{s2}}{{kmhif}^{s2}+HIFa^{s2}}$$

$$CREB_{Ptv_{PGC}}=fcreb1*\left( \frac{CREB^{s2}}{CREB^{s2}+{kmcreb1}^{s2}} \right)$$

$$AMPK_{Ptv_{PGC}}=fampk1*\left( \frac{AMPK^{s2}}{AMPK^{s2}+{kmampk1}^{s2}} \right)$$

$$TNF_{Ntv_{PGC1}}=\left( 1+ftnfil6*\left( \frac{\left( TNF+IL6 \right)^{s2}}{{kmtnfil6}^{s2}+\left( TNF+IL6 \right)^{s2}} \right) \right)$$

$$\dot{\boldsymbol{PGC1}}=kpc0+\left( kpc1*\left( CREB_{Ptv_{PGC}}+FOXO_{Ptv_{PGC}}+GCR_{Ptv}+NAD_{Ptv_{PGC}}+AMPK_{Ptv_{PGC}} \right)*HIF_{Ntv_{PGC}} \right)$$

$$-kpc2*PGC1*\left( AKT_{Ntv_{PGC}}*TNF_{Ntv_{PGC1}} \right);$$

$$PI3K_{Ptv_{TRB}}=fpi3k*\left( \frac{{PI3k}^{1.5}}{{PI3k}^{1.5}+{kmpi3k}^{1.5}} \right)$$

$$PKC_{Ptv_{TRB}}=fpkc2*\left( \frac{PKC}{PKC+kmpkc2} \right)$$

$$PPAR_{Ptv_{TRB3}}=fppar*\left( \frac{PPARab^{s2}}{PPARab^{s2}+{kmppar}^{s2}} \right)$$

$$PGC_{Ptv_{PPARab}}=fpgc1*\left( \frac{PGC1^{s2}}{PGC1^{s2}+{kmpgc1}^{s2}} \right)$$

$$\dot{\boldsymbol{TRB}\boldsymbol{3}}=ktr0+\left( ktr1*PI3K_{Ptv_{TRB}}*PKC_{Ptv_{TRB}}*\left( PPAR_{Ptv_{TRB3}}+PGC_{Ptv_{PPARab}}+GCR_{Ptv} \right) \right)-\left( ktr2*TRB3 \right)$$

**Characterization of the metabolic state**

To quantify the net metabolic state (either catabolic or anabolic) for per parameter perturbation we devised equation analogous to formulation presented for phosphorylation state by (Bulik et al., 2016; König et al., 2012). The formulation represents an ensemble of the difference between the scaled effects of state variables of anabolic and catabolic signaling pathways and the corresponding states of transcriptional factors. The metabolic state is scaled between 0 and 1, with a basal state around 0.5. The metabolic state is represented by,

$Metabolic state=0.5*\left( \begin{aligned} 1+\left( \begin{aligned} 0.2*\left( \left( \frac{PKA}{h_{Pka}+PKA} \right)+\left( \frac{Cal}{h_{Cal}+Cal} \right)+\left( \frac{CREB}{h_{CREB}+CREB} \right)+\left( \frac{AMPK}{h_{AMPK}+AMPK} \right)+\left( \frac{HIF\alpha}{h_{HIF}+HIF\alpha} \right) \right) \\ -0.2*\left( \left( \frac{AKT}{h_{Akt}+AKT} \right)+\left( \frac{mTOR}{h_{mTOR}+mTOR} \right)+\left( \frac{SREBP}{h_{SREBP}+SREBP} \right)+\left( \frac{CHREBp}{h_{CHREBP}+CHREBp} \right)+\left( \frac{PPARg}{h_{PPARg}+PPARg} \right) \right) \end{aligned} \right) \end{aligned} \right)$

where, *PKA, Cal, CREB, AMPK* and *HIF* are the catabolic effector state variables; and *AKT, mTOR, SREBP, CHREBP* and *PPARg* are the anabolic state variables. The *h* factor in the denominator is the half maximal saturation constant which is scaled difference of the maximum and minimum levels of the corresponding variable.

**Appendix III : Metabolic Control Analysis**

Table S5: Metabolic concentration response coefficients for the model parameters that reproduced MD signature. (-) prefix indicates reduction in the parameter resulted in MD signature. The list includes parameters that yielded the MCRC of at least 0.001 for each of the 12 metabolites in the MD signature. The table includes the 34 parameters reported in Figure 2A (MCRC of at least 0.1).

| **Model Parameters** | **MCRCs** | **Model Parameters** | **MCRCs** |
| --- | --- | --- | --- |
| **Metabolic fluxes** |  |  | |
| α-ketoglutarate dehydrogenase flux | 0.317 | Activation of TNF | 0.014 |
| Glycerol phosphorylation | 0.179 | Saturation constant of phagocytes on TNF | 0.182 |
| (-) Lipolysis | 0.028 | Rate of TNF positive feedback | 0.142 |
| Triglyceride synthesis | 0.175 | (-) Degradation of TNF | 0.183 |
| (-) Alanine to protein synthesis | 0.041 | Plasma Endotoxin concentration | 0.203 |
| Protein breakdown to amino acids | 0.032 | (-)Rate of TGF activation by PI3k | 0.003 |
| Plasma glycerol levels | 0.173 | Activation of TNF by CREB/NFkb | 0.134 |
| Plasma Amino acid (except glutamine) | 0.490 | (-) Sensitivity of anti-inflammatory effect of TGF on TNF | 0.070 |
| (-) Degradation rate of SREBP1c | 0.340 | Saturation constant for effect of TGF on TNF | 0.308 |
| **GPCR signaling pathway** |  | **HPA-axis and GCR signaling** |  |
| (-) Saturation constant of glucagon secretion | 0.442 | (-) Activation of anti-inflammatory response by GCR | 0.504 |
| Catecholamine/ Glucagon secretion | 0.494 | Saturation constant of GCR for anti-inflammatory action | 0.405 |
| (-)Degradation of Catecholamine | 0.334 | Sensitivity of GCR negative feedback on inflammation | 0.838 |
| (-)β-adrenergic receptor-ligand dissociation constant | 0.252 | (-) Sensitivity of GCR central negative feedback | 0.710 |
| (-)Rate of GPCR sequestration | 0.258 | Inhibitory constant of HPA negative feedback | 0.383 |
| (-)Rate of G-protein deactivation | 0.260 | Stress parameter at CRH (HPA axis) | 0.192 |
| Degradation of GPCR activated PLC | 0.306 | (-) Sensitivity of HPA axis to inflammatory cytokines | 0.253 |
| Activation of calcium signaling by G-protein | 0.485 | (-)Degradation of CRH | 0.334 |
| (-) Suppression pf calcium signaling | 0.336 | Activation of ACTH by CRH | 0.304 |
| **Inflammatory signaling pathway** |  | (-)Degradation of ACTH | 0.342 |
| Rate of phagocyte activation | 0.207 | Activation rate of CRH by TNF | 0.142 |
| (-)Rate of TGF activation by phagocytes | 0.003 | (-) Activation threshold of HPA axis by TNF | 0.042 |
| Degradation of TGF anti-inflammatory cytokine | 0.410 | Sensitivity of GCR negative feedback (central+immune) | 1.005 |

**Model-based inference from simulated regulatory states**

We recorded the states of regulatory components such as ATP/ADP and NADH/NAD ratio, anabolic and catabolic signaling, transcriptional regulators and inflammation as shown in Figure 3. It was consistently noted that metabolic controller ratio ATP/ADP was reduced for all the 27 parameter perturbation cases. The redox ratio NADH/NAD was elevated for upregulation of GPCR signaling, whereas it was reduced for the parameters for HPA axis activation associated inflammatory state. In the insulin signaling pathway, AKT phosphorylation was increased relative to the basal level except for the parameters of metabolic flux category, with reduced IRS tyrosine phosphorylation for GPCR parameters. The components of amino acid metabolic regulators, mTOR pathway (mTOR raptor and S6K1p) and transcriptional regulators of lipid synthesis pathway (SREBP and PPARg) were also activated for perturbation in all the parameters of the regulatory category. Among the components of GPCR pathway, β-adrenergic receptor, G-protein and calcium signaling were activated for all the parameters along with the upregulation of protein kinase A (PKA) and cyclic AMP (cAMP). The global transcriptional regulator CREB was activated across all parameters. The transcriptional regulator for lipid catabolism PPARα/β was down regulated throughout the perturbations in the parameters of HPA axis and inflammation and was upregulated for GPCR signaling. The transcriptional regulator for urea cycle, CEBPα was upregulated only for the parameters in the inflammation and HPA axis classes, whereas downregulated for GPCR parameters. The transcriptional regulator for mitochondrial biogenesis, PGC1α was also marginally downregulated for the parameter perturbations in glucocorticoid receptor sensitivity and inflammatory signaling, whereas remained unchanged for the parameters in GPCR and other HPA axis signaling. It was also noted that the hypoxia inducible factor HIF1α and AMPK (AMP kinase) were upregulated throughout the perturbations in the parameters of all the four categories. It was also noted that nuclear translocation of glucocorticoid receptor was enhanced for the parameters of the HPA axis and inflammatory signaling, with increased levels of cortisol and ACTH. The levels of inflammatory cytokines TNF and IL6 were upregulated for the parameters of inflammatory signaling and downregulated for the HPA axis perturbations except for increase glucocorticoid sensitivity. The metabolic state was catabolic across all the parameters listed except for elevated SREBP1c expression and plasma amino acid flux.

**Appendix IV: Correlational Analysis**

To assess the model-based hypothesis on the effects of variables associated with the four categories inferred from MCA, we performed correlation analysis to estimate Spearman Correlation Coefficients (SCC: ρ) between the regulatory components and the metabolites that show statistically significant differences in the cohorts.

**Selection of metabolites for correlational analysis**

The selection of metabolites for further analysis was made based on the statistical significance and the relevance in context of the model based hypothesis. Due to their associations with the metabolic pathways that showed statistically significant differences, despite the trend level significance between groups in citrate, isoleucine, glycerate, carnitine and palmitoylcarnitines, branched chain amino acids (leucine and valine) and arginine, these metabolites were included in the correlational analysis (features suffixed by ‘*’ sign in Table 1). Since the essential fatty acids needs to be absorbed from the food and are not likely to be associated with the perturbations in metabolic pathways, we excluded them from further analysis. We also excluded metabolites from other categories, electrolytes and immune cells from the analysis (Table 1), as they were not directly associated with the pathways highlighted by the model analysis. Therefore, we used a total of 35 metabolites and 8 regulatory components were retained for correlational (See Figure S5).

**Correlational Analysis for regulatory correlates in PTSD subjects**

We performed exploratory correlation analysis to obtain SCCs between these metabolites and regulatory components. The statistically significant correlation for the PTSD subjects are reported in the Table S6 along with the correlation coefficients and respective p-values and q-values. We obtained correlation coefficients on 35x8 matrices separately for controls and PTSD and compared the changes in associations across the cohorts. Supplementary Figure 3 shows the statistically significant (p<0.05) correlation matrices for (a) entire cohort (b) controls only, (c) PTSD only and (d) relative difference in correlations between controls and PTSD.

It was observed that the statistically significant correlations were either identical or marginally different in PTSD, maintaining the statistical significance in correlations across the cohorts. It is noted that the cross correlations for hs-CRP, GGT, hypoxanthine and the metabolites were elevated 5-10% in PTSD with respect to controls (Figure S6D). Cortisol suppression by DEX strongly correlated with HOMA-IR, GGT and several other metabolites, corroborating the model based hypothesis (Figure S6A, S6B and S6C). IC50 showed a trend of negative correlation with cortisol levels (ρ=-0.167, p=0.032, q=0.082) in the entire cohort. It also showed a correlation with hypoxanthine (ρ=-0.164, p=0.036, q=0.086) indicating an association of energy deficit or hypoxia with the reduction in IC50 (Leonard et al., 2005). Moreover, in the entire cohort, HOMA-IR, hs-CRP, GGT and hypoxanthine showed statistically significant cross correlations. The correlation coefficients between the 8 regulatory components and 35 metabolites in the entire cohort are reported in Table S6. We divided the analysis based on the affected metabolic pathways in five sections, namely (i) glycolysis and TCA cycle, (ii) amino acid metabolism, (iii) fatty acid metabolism, (iv) oxidative stress, inflammation and insulin resistance and an additional inference on (v) hepatic function. Here we report the statistically significant correlation coefficients with at least p<=0.05 and q<=0.1.

**Glycolysis and TCA cycle**:

In glycolysis, glucose levels were positively correlated hs-CRP (ρ=0.22, p=0.042, q=0.1) and HOMAIR (ρ =0.55, p=7.7E-8, q=1.7E-6). Pyruvate level was positively associated with HOMA-IR (ρ =0.21 trend), hypoxanthine (ρ =0.58, p=6.9E-9, q=1.66E-7) and GGT (ρ =0.25, p=0.022, q=0.072). Plasma lactate levels were correlated with GR methylation (ρ =0.21 trend), cortisol suppression (ρ =0.31, p=4.1E-3, q=0.023), HOMA-IR (ρ =0.45, p=1.97E-5, q=2.25E-4), hypoxanthine (ρ =0.47, p=7.36E-6, q=9.3E-5) and GGT (ρ =0.35, p=1.3E-3, q=9.5E-4). Moreover, hypoxanthine showed a trend level significance of negative association with plasma phosphate levels and positive association with adenosine monophosphate (AMP).

**Amino acid metabolism:**

Plasma levels of alanine were positively correlated with cortisol suppression (ρ =0.33, p=2E-3, q=1.3E-2) and HOMA-IR (ρ =0.19, trend). Plasma glutamine levels showed a negative correlation with GGT (ρ = -0.20, trend). Plasma tyrosine levels were also positively associated with cortisol suppression (ρ =0.24, p=0.028, q=0.08), hs-CRP (ρ =0.26, p=0.017, q=0.062), HOMA-IR (ρ =0.39, p=2.5E-4, q=2.2E-3), hypoxanthine (ρ =0.22, p=0.044, q=0.1) and GGT (ρ =0.52, p=5.4E-7, q=1E-5). Plasma branched chain amino acid levels (BCCAs) isoleucine, leucine and valine were positively correlated with hs-CRP (ρ=0.23, p=0.036, q=0.1; ρ=0.26, p=0.016, q=0.057; ρ =0.25, p=0.025, q=0.075; respectively); HOMA-IR (ρ =0.27, p=0.015, q=0.057; SCC=0.32, p=0.003, q=0.02; ρ =0.25, p=0.021, q=0.07; respectively); hypoxanthine (ρ =0.18 (trend); ρ=0.22, p=0.046, q=0.1; ρ =0.25,p=0.025, q=0.075; respectively); GGT (ρ =0.35, p=1.2E-3, q=0.009; ρ =0.34, p=1.8E-3, q=0.012; ρ =0.30, p=6.5E-3, q=0.03; respectively); whereas, cortisol suppression showed a positive trend and urinary epinephrine showed a negative trend with BCCAs. Moreover, the urea cycle metabolite ornithine, correlated positively with a trend level association with cortisol suppression (ρ =0.20) and a negative association with urinary epinephrine (ρ =-0.34, p=1.65E-3, q=0.012).

**Lipid metabolism:**

Plasma levels of nonadecenoate (19), methylstearate, undecenoate and stearate showed a consistent negative trend of correlation with cortisol suppression. Urine epinephrine was positively associated with nonadecanoates (ρ=0.28, p=0.011, q=0.047), methylstearate (ρ =0.29, p=7.4E-3, q=0.034), hydroxypalmitate (ρ =0.24, p=0.027, q=0.078), arachidonoate (ρ =0.27, p=0.012, q=0.05) and stearate (ρ =0.25, p=0.024, q=0.075). Plasma carnitine showed a positive trend with cortisol suppression (ρ =0.19). Plasma palmitoylcarnitine was positively associated with HOMAIR (ρ =0.23, p=0.035, q=0.09), GGT (ρ =0.3, p=0.005, q=0.026) and hypoxanthine (ρ =0.33, p=0.002, q=0.016). Further, plasma triglycerides were positively correlated with hs-CRP (ρ =0.51, p=9.9E-7, q=1.6E-5), HOMA-IR (ρ=0.47, p=8.24E-6, q=9.9E-5) and GGT (ρ =0.43, p=4.5E-5, q=4.7E-4), whereas glycerate showed a negative association with HOMAIR (ρ=-0.29, p=0.008, q=0.034), hs-CRP (SCC=-0.22, p=0.041, q=0.1), GGT(ρ =-0.44, p=2.8E-5, q=3E-4) and hypoxanthine (ρ =-0.4, p=2E-4, q=0.002).

**Hepatic function:**

Plasma levels of albumin were positively associated with HOMAIR (ρ =0.24, p=0.028, q=0.079). Alkaline phosphatase levels were positively associated with HOMAIR (ρ =0.4, p=2E-4, q=0.002), GGT (ρ =0.48, p=4.3E-6, q=5.7E-5). Plasma total protein levels were positively associated with HOMAIR (ρ =0.26, p=0.019, q=0.066), hs-CRP (ρ =0.31, p=0.004, q=0.023), GGT (ρ =0.32, p=0.003, q=0.02). Moreover cortisol suppression showed a positive trend with plasma alkaline phosphatatse and total proteins (ρ ~0.2).

**Oxidative stress, Inflammation and Insulin Resistance:**

GGT correlated positively with cortisol suppression (ρ =0.25, p=0.022, q=0.07), the components of glutathione pathway: gamma glutamyl tyrosine (ρ =0.34, p=0.002, q=0.013) and negative association with oxoproline (ρ =-0.22, trend) and stearoyl sphingomyeline (ρ =0.24, p=0.03, q=0.08). Gammaglutamyl tyrosine also showed a positive association with HOMAIR (ρ=0.23, p=0.04, q=0.1), hs-CRP (ρ =0.32, p=0.003, q=0.02) and hypoxanthine (ρ =0.22, p=0.044, q=0.1). Oxoproline was also associated with hypoxanthine (ρ=0.28, p=0.011, q=0.047). GGT was also positively correlated with hs-CRP (ρ =0.51, p=7.58E-7, p=1.3E-5), HOMA-IR (ρ =0.49, p=2.5E-6, q=3.6E-5) and hypoxanthine (ρ =0.30, p=6.3E-3, q=0.03). Further, the inflammatory cytokine IL6 showed a positive correlation with GGT (ρ =0.24, p=0.027, q=0.08), HOMA-IR (ρ =0.26, p=0.018, q=0.062), hs-CRP (ρ =0.27, p=0.013, q=0.052), IC50 (ρ =0.24, p=0.028, q=0.08) and a negative correlation with cortisol (ρ =-0.24, p=0.032, q=0.09). Sphingosine 1 phosphate, a signaling component activated by inflammatory process showed positive association with hsCRP (ρ =0.26, p=0.017, q=0.06), GGT (ρ =0.22, p=0.047, q=0.1) and hypoxanthine (ρ =0.3, p=0.006, q=0.028). Moreover, hs-CRP was positively associated with HOMA-IR (ρ =0.31, p=3.9E-3, q=0.022) and hypoxanthine (ρ =0.23). HOMA-IR was also positively associated with cortisol suppression (ρ =0.27, p=0.013, q=0.05) and hypoxanthine (ρ =0.25, p=0.025, q=0.075).

Table S6: Regulatory correlated of metabolic dysfunction in entire cohort (PTSD+Controls). Red and yellow shade highlights the p and q values <=0.05 and between 0.05 and 0.1, respectively.

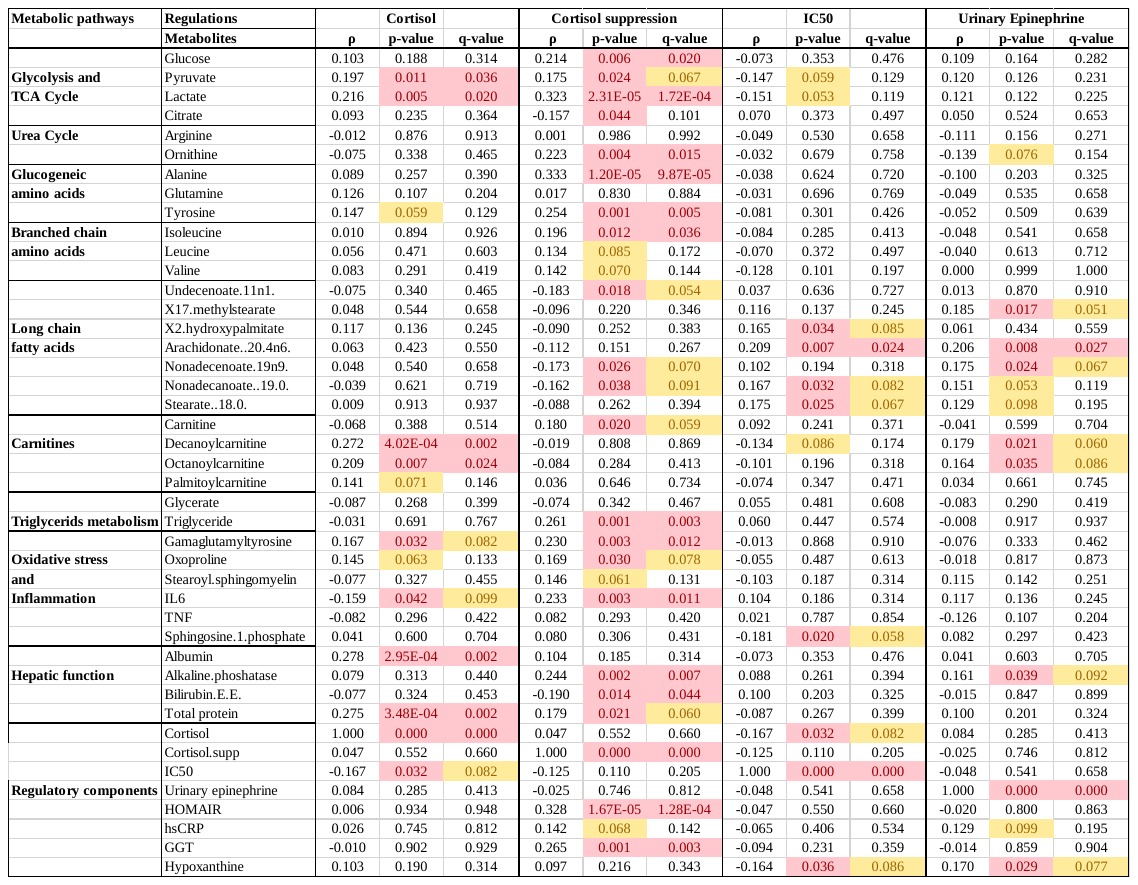

Table S6 (continued)

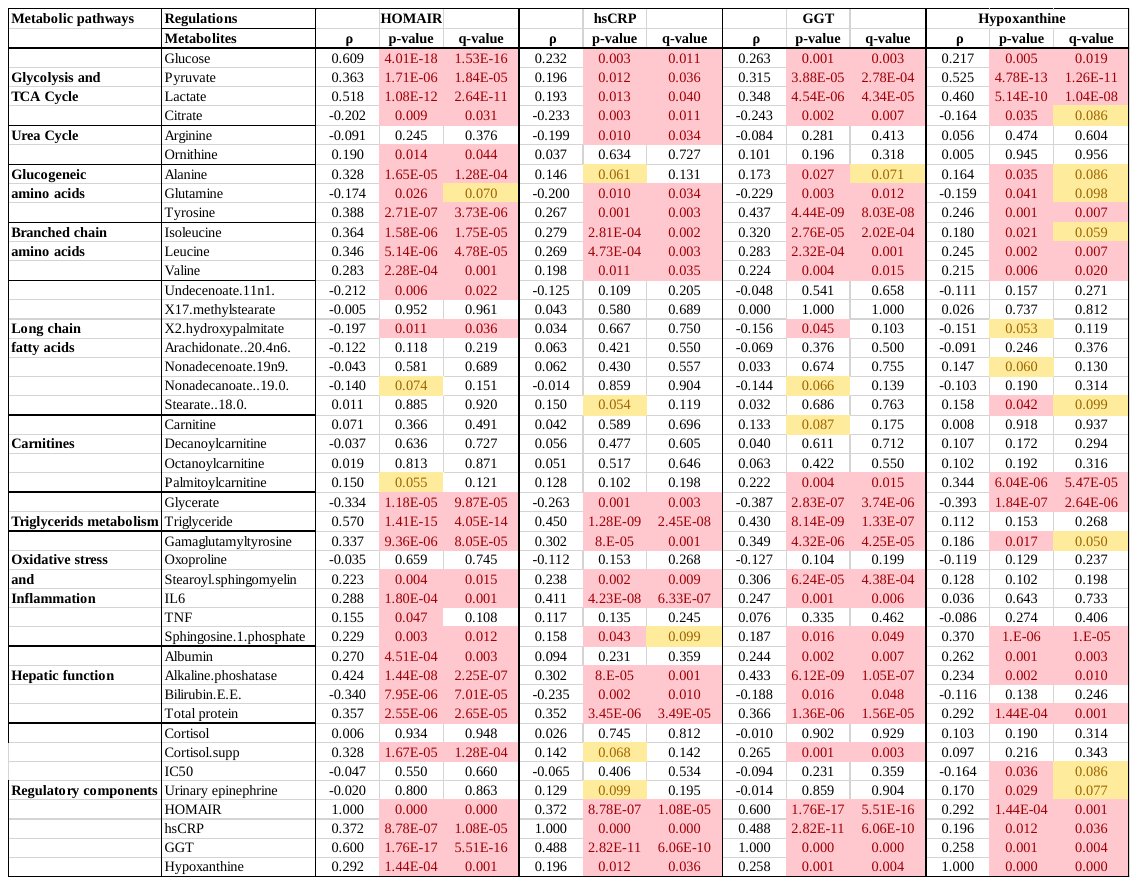

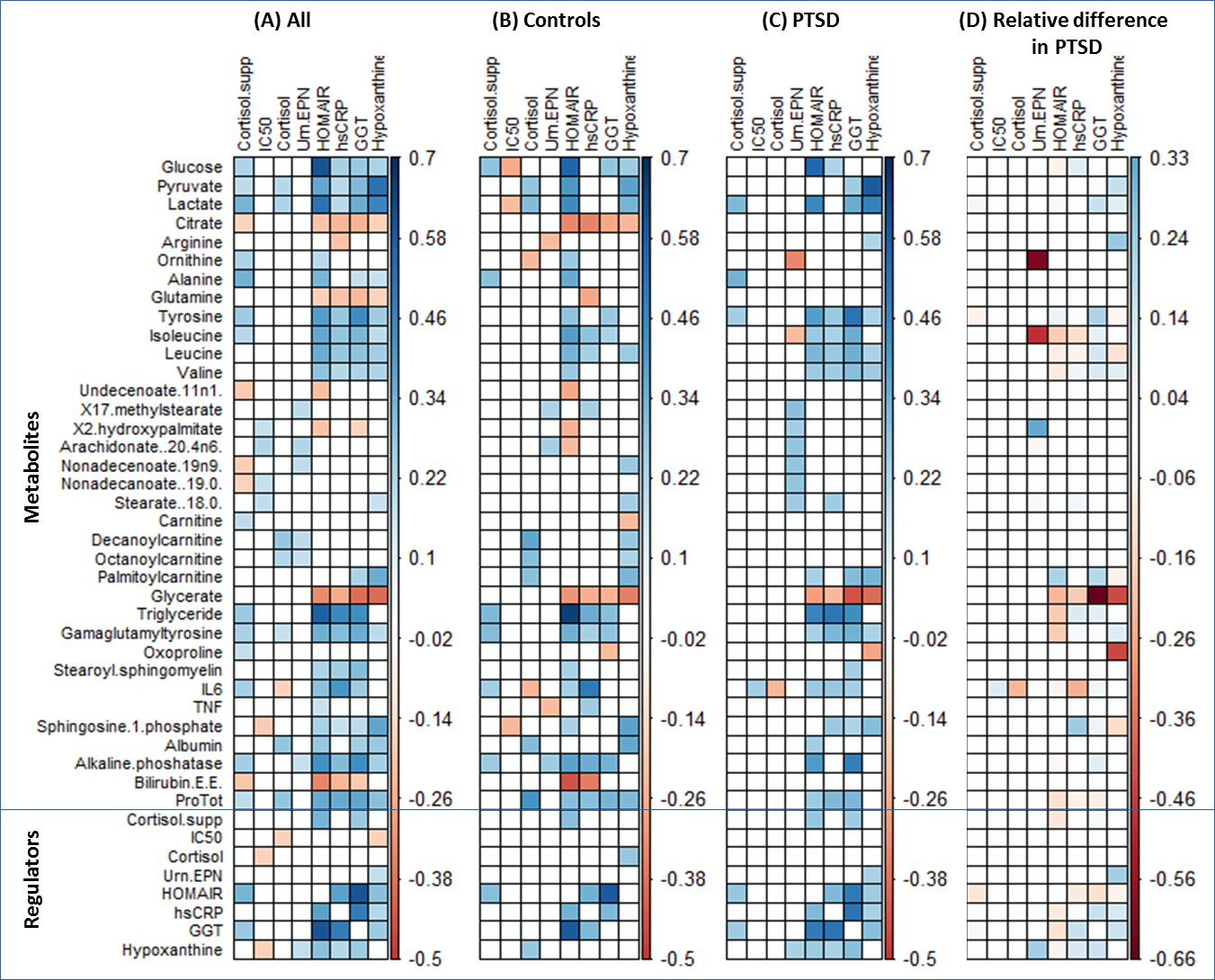

Figure S5: Correlation plot for regulatory components with respect to metabolites in (A) entire cohort, (B) controls, (C) PTSD subjects, and (D) relative difference with respect to correlations in PTSD. The correlations are shown for the statistically significant correlations (p<0.05). It is observed that across the correlation plots A, B and C, significant correlations are conserved for the cross correlations between HOMA-IR, GGT, hsCRP, hypoxanthine and glycolytic metabolites, amino acid metabolites and triglyceride metabolites. On the relative difference plot, it is noted that there is a marginal difference between correlations in controls and PTSD, with around 5-15% higher correlations in PTSD subjects between GGT, hsCRP and hypoxanthine indicating stronger association between oxidative stress and inflammation; along with similar level of increase in correlation between glycolytic metabolites, arginine and hypoxanthine. There is a notable decrease of correlations related to urinary epinephrine with ornithine and tyrosine, GGT and glycerate, HOMA-IR and triglyceride and hypoxanthine and oxoproline in PTSD.

**Appendix-V: Causal mediation Inference**

In the mediation analysis with cortisol suppression as continuous exposure variable and HOMA-IR, hs-CRP, GGT and hypoxanthine as joint mediators, we observed statistically significant joint mediated effects on metabolites. Following are the estimates of the effects and corresponding confidence intervals for metabolites. Glycolytic components: pyruvate (*ψ_i_*=0.644 (0.238, 1.05)) and lactate (*ψ_i_*=0.319 (0.146, 0.493)); citrate *(ψ_i_*=-0.134 (-0.228, -0.039); amino acids: tyrosine (*ψ_i_*=0.189 (0.108, 0.27)), isoleucine (*ψ_i_*=0.162 (0.07, 0.254)), leucine (*ψ_i_*=0.123 (0.051, 0.196)), valine (*ψ_i_*=0.093 (0.017, 0.169)); glycerate (*ψ_i_*=-0.239 (-0.363, -0.116)), triglycerides (*ψ_i_*=0.12 (0.061, 0.163)), gammaglutamyl tyrosine (*ψ_i_*=0.177 (0.077, 0.277)), oxoproline (*ψ_i_*=-0.066 (-0.126, -0.005)), stearoyl sphingomyelin (*ψ_i_*=0.205 (0.081, 0.329)), IL6 (*ψ_i_*=0.433 (0.08, 0.786)), sphingosine 1 phosphate (*ψ_i_*=0.288 (0.071, 0.505)), hepatic function: albumin (*ψ_i_*=0.019 (0.004, 0.033)), alkaline phosphatase (*ψ_i_*=0.05 (0.025, 0.074)) and total plasma protein (*ψ_i_*=0.019 (0.006, 0.031)). Additionally, the total causal effects were also significant on ornithine (*ψ_t_*=0.402 (0.012, 0.793)), alanine (*ψ_t_*=0.472 (0.1, 0.843)) and bilirubin (*ψ_t_*=-1.216 (-2.142, -0.291)). The analysis indicates that the effects of changes in GR sensitivity on metabolic regulation are relayed through changes in inflammation, oxidative stress, insulin resistance and energy deficit. Figure S7 shows the coefficients and statistics for the NDE, NIE and the total causal effect (TCE).

Table S7: The table reports the estimates and statistics for natural direct, natural indirect and total causal effect of glucocorticoid receptor sensitivity assessed by DEX suppression test on metabolites. Red and yellow shade highlights the p and q values <0.05 and between 0.05 and 0.1, respectively.

| **Metabolic pathways** | **Natural Effects** |  |  | **Natural Direct Effect** | | |  |  |  | **Natural Indirect Effect** | | |  |  | **Total Causal Effect** | | |  |  |
| --- | --- | --- | --- | --- | --- | --- | --- | --- | --- | --- | --- | --- | --- | --- | --- | --- | --- | --- | --- |
|  | **Metabolites** | **NDE (ψd)** | **Std. error** | **CI-lower** | **CI-upper** | **p-value** | **q-value** | **NIE (ψi)** | **Std. error** | **CI-lower** | **CI-upper** | **p-value** | **q-value** | **TCE (ψt)** | **Std. error** | **CI-lower** | **CI-upper** | **p-value** | **q-value** |
| **Glycolysis and**  **TCA Cycle** | Glucose | 0.002 | 0.025 | -0.047 | 0.051 | 0.938 | 0.984 | 0.012 | 0.011 | -0.011 | 0.034 | 0.304 | 0.380 | 0.014 | 0.023 | -0.031 | 0.058 | 0.544 | 0.732 |
|  | Pyruvate | 0.181 | 0.413 | -0.628 | 0.990 | 0.662 | 0.896 | 0.644 | 0.207 | 0.238 | 1.050 | 0.002 | 0.007 | 0.825 | 0.404 | 0.032 | 1.617 | 0.041 | 0.138 |
|  | Lactate | 0.000 | 0.230 | -0.450 | 0.450 | 0.999 | 0.999 | 0.319 | 0.088 | 0.146 | 0.493 | 3.01E-04 | 0.002 | 0.319 | 0.216 | -0.105 | 0.743 | 0.140 | 0.273 |
|  | Citrate | -0.347 | 0.204 | -0.747 | 0.054 | 0.090 | 0.492 | -0.134 | 0.048 | -0.228 | -0.039 | 0.006 | 0.016 | -0.480 | 0.195 | -0.864 | -0.097 | 0.014 | 0.098 |
| **Urea Cycle** | Arginine | -0.065 | 0.170 | -0.398 | 0.268 | 0.701 | 0.896 | 0.024 | 0.059 | -0.092 | 0.140 | 0.686 | 0.774 | -0.041 | 0.163 | -0.361 | 0.279 | 0.801 | 0.852 |
|  | Ornithine | 0.292 | 0.210 | -0.119 | 0.703 | 0.164 | 0.492 | 0.110 | 0.089 | -0.065 | 0.286 | 0.218 | 0.294 | 0.402 | 0.199 | 0.012 | 0.793 | 0.044 | 0.138 |
| **Glucogeneic**  **amino acids** | Alanine | 0.347 | 0.224 | -0.091 | 0.785 | 0.121 | 0.492 | 0.125 | 0.077 | -0.025 | 0.275 | 0.103 | 0.172 | 0.472 | 0.189 | 0.100 | 0.843 | 0.013 | 0.098 |
|  | Glutamine | -0.047 | 0.107 | -0.258 | 0.163 | 0.659 | 0.896 | -0.053 | 0.028 | -0.108 | 0.002 | 0.060 | 0.116 | -0.100 | 0.100 | -0.297 | 0.097 | 0.318 | 0.506 |
|  | Tyrosine | 0.141 | 0.176 | -0.204 | 0.487 | 0.423 | 0.779 | 0.189 | 0.041 | 0.108 | 0.270 | 4.34E-06 | 1.52E-04 | 0.330 | 0.177 | -0.016 | 0.676 | 0.061 | 0.161 |
| **Branched chain**  **amino acids** | Isoleucine | 0.123 | 0.120 | -0.111 | 0.358 | 0.302 | 0.621 | 0.162 | 0.047 | 0.070 | 0.254 | 0.001 | 0.003 | 0.285 | 0.123 | 0.045 | 0.526 | 0.020 | 0.110 |
|  | Leucine | 0.036 | 0.103 | -0.165 | 0.237 | 0.728 | 0.896 | 0.123 | 0.037 | 0.051 | 0.196 | 0.001 | 0.004 | 0.159 | 0.104 | -0.044 | 0.362 | 0.124 | 0.256 |
|  | Valine | -0.007 | 0.121 | -0.245 | 0.231 | 0.952 | 0.984 | 0.093 | 0.039 | 0.017 | 0.169 | 0.016 | 0.036 | 0.086 | 0.120 | -0.149 | 0.321 | 0.472 | 0.661 |
| **Long chain**  **fatty acids** | 10-Nonadecenoate 19:1 (ω-9) | 0.139 | 0.280 | -0.410 | 0.688 | 0.619 | 0.896 | -0.127 | 0.099 | -0.320 | 0.066 | 0.198 | 0.278 | 0.012 | 0.268 | -0.513 | 0.538 | 0.964 | 0.964 |
|  | 10-Undecenoate 11:1 (ω-1) | -0.117 | 0.357 | -0.817 | 0.583 | 0.742 | 0.896 | 0.032 | 0.098 | -0.159 | 0.223 | 0.744 | 0.814 | -0.086 | 0.345 | -0.762 | 0.591 | 0.804 | 0.852 |
|  | 17-Methylstearate | 0.040 | 0.142 | -0.239 | 0.319 | 0.777 | 0.903 | -0.070 | 0.040 | -0.147 | 0.008 | 0.080 | 0.140 | -0.029 | 0.135 | -0.293 | 0.235 | 0.828 | 0.852 |
|  | 2-Hydroxypalmitate | -0.484 | 0.327 | -1.125 | 0.156 | 0.139 | 0.492 | -0.019 | 0.081 | -0.178 | 0.140 | 0.813 | 0.862 | -0.503 | 0.305 | -1.100 | 0.094 | 0.098 | 0.215 |
|  | Nonadecanoate 19:0 | -0.869 | 0.453 | -1.758 | 0.019 | 0.055 | 0.435 | 0.004 | 0.090 | -0.174 | 0.181 | 0.968 | 0.968 | -0.866 | 0.445 | -1.739 | 0.007 | 0.052 | 0.151 |
|  | Arachidonate 20:4 (ω-6) | -0.390 | 0.276 | -0.931 | 0.152 | 0.158 | 0.492 | -0.070 | 0.065 | -0.197 | 0.057 | 0.280 | 0.363 | -0.460 | 0.264 | -0.976 | 0.057 | 0.081 | 0.190 |
|  | Stearate 18:0 | -0.276 | 0.214 | -0.694 | 0.143 | 0.197 | 0.492 | 0.036 | 0.053 | -0.068 | 0.140 | 0.494 | 0.577 | -0.239 | 0.206 | -0.643 | 0.164 | 0.245 | 0.412 |
| **Carnitines** | Carnitine | 0.035 | 0.138 | -0.235 | 0.305 | 0.800 | 0.903 | 0.004 | 0.031 | -0.057 | 0.066 | 0.889 | 0.916 | 0.039 | 0.132 | -0.220 | 0.299 | 0.767 | 0.852 |
|  | Decanoylcarnitine | 0.690 | 0.599 | -0.484 | 1.864 | 0.249 | 0.582 | -0.234 | 0.160 | -0.547 | 0.080 | 0.144 | 0.219 | 0.456 | 0.590 | -0.700 | 1.613 | 0.439 | 0.641 |
|  | Octanoylcarnitine | 0.437 | 0.581 | -0.703 | 1.576 | 0.453 | 0.792 | -0.201 | 0.153 | -0.500 | 0.099 | 0.189 | 0.275 | 0.236 | 0.574 | -0.888 | 1.360 | 0.681 | 0.796 |
|  | Palmitoylcarnitine | -0.249 | 0.275 | -0.789 | 0.291 | 0.366 | 0.712 | 0.104 | 0.069 | -0.032 | 0.240 | 0.132 | 0.211 | -0.145 | 0.272 | -0.678 | 0.389 | 0.595 | 0.744 |
| **Triglycerids metabolism** | Glycerate | 0.136 | 0.261 | -0.375 | 0.648 | 0.602 | 0.896 | -0.239 | 0.063 | -0.363 | -0.116 | 1.49E-04 | 0.001 | -0.103 | 0.252 | -0.597 | 0.390 | 0.682 | 0.796 |
|  | Triglyceride | 0.058 | 0.083 | -0.105 | 0.221 | 0.486 | 0.810 | 0.112 | 0.026 | 0.061 | 0.163 | 1.55E-05 | 2.72E-04 | 0.170 | 0.084 | 0.006 | 0.334 | 0.042 | 0.138 |
| **Oxidative stress**  **and**  **Inflammation** | Gamaglutamyltyrosine | 0.203 | 0.188 | -0.166 | 0.572 | 0.281 | 0.615 | 0.177 | 0.051 | 0.077 | 0.277 | 0.001 | 0.003 | 0.380 | 0.188 | 0.012 | 0.748 | 0.043 | 0.138 |
|  | 5-Oxoproline | 0.182 | 0.136 | -0.085 | 0.448 | 0.181 | 0.492 | -0.066 | 0.031 | -0.126 | -0.005 | 0.033 | 0.068 | 0.116 | 0.130 | -0.138 | 0.371 | 0.371 | 0.565 |
|  | Stearoyl sphingomyelin | 0.011 | 0.206 | -0.392 | 0.415 | 0.955 | 0.984 | 0.205 | 0.063 | 0.081 | 0.329 | 0.001 | 0.004 | 0.217 | 0.187 | -0.150 | 0.583 | 0.247 | 0.412 |
|  | IL6 | 0.733 | 0.355 | 0.037 | 1.429 | 0.039 | 0.435 | 0.433 | 0.180 | 0.080 | 0.786 | 0.016 | 0.036 | 1.166 | 0.327 | 0.524 | 1.808 | 3.70E-04 | 0.013 |
|  | TNFα | 0.370 | 0.197 | -0.016 | 0.756 | 0.060 | 0.435 | -0.044 | 0.060 | -0.162 | 0.073 | 0.459 | 0.553 | 0.326 | 0.176 | -0.019 | 0.670 | 0.064 | 0.161 |
|  | Sphingosine 1 phosphate | -0.470 | 0.357 | -1.169 | 0.229 | 0.187 | 0.492 | 0.288 | 0.111 | 0.071 | 0.505 | 0.009 | 0.025 | -0.182 | 0.342 | -0.852 | 0.488 | 0.595 | 0.744 |
| **Hepatic function** | Albumin | 0.009 | 0.025 | -0.040 | 0.058 | 0.717 | 0.896 | 0.019 | 0.007 | 0.004 | 0.033 | 0.012 | 0.030 | 0.028 | 0.023 | -0.018 | 0.073 | 0.232 | 0.412 |
|  | Alkaline phoshatase | 0.092 | 0.049 | -0.005 | 0.189 | 0.062 | 0.435 | 0.050 | 0.013 | 0.025 | 0.074 | 7.11E-05 | 0.001 | 0.142 | 0.048 | 0.047 | 0.237 | 0.003 | 0.058 |
|  | Bilirubin EE | -0.995 | 0.499 | -1.973 | -0.017 | 0.046 | 0.435 | -0.221 | 0.120 | -0.457 | 0.015 | 0.066 | 0.122 | -1.216 | 0.472 | -2.142 | -0.291 | 0.010 | 0.098 |
|  | Total protein | 0.031 | 0.023 | -0.013 | 0.076 | 0.163 | 0.492 | 0.019 | 0.006 | 0.006 | 0.031 | 0.003 | 0.011 | 0.050 | 0.022 | 0.007 | 0.093 | 0.022 | 0.110 |

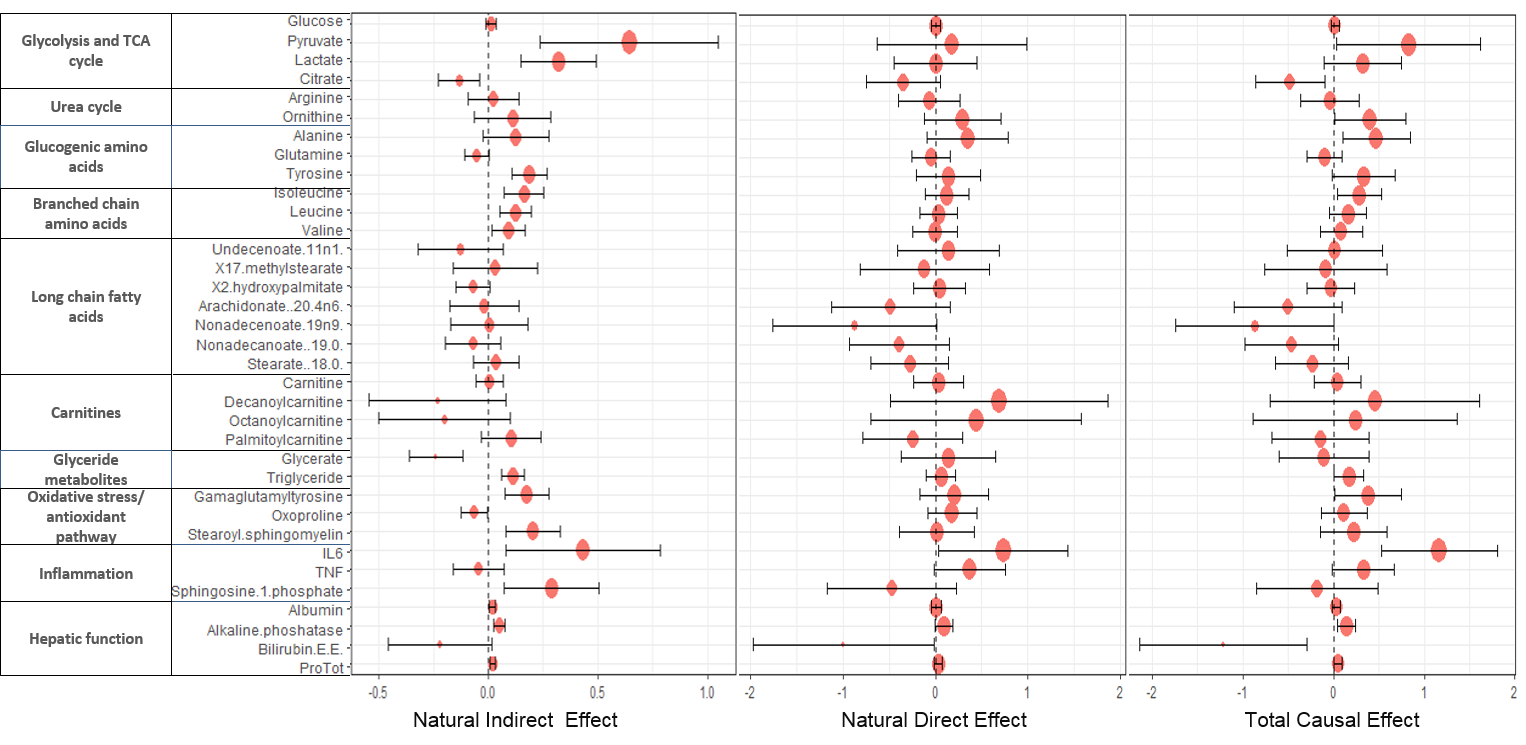

Figure S6: Forest plot representation of the natural indirect effects (joint mediated effects), natural direct effects and total causal effects of GR sensitivity (measured by cortisol suppression test) on 35 metabolites for the causal hypothesis tested on the entire cohort adjusting for the group effects. The error bar represents 95% confidence intervals of the point estimates of the effects. It is noted that the joint mediated effects on pyruvate, lactate, citrate, gluconeogenic and branched chain amino acids, oxidative stress, inflammation and hepatic function components are statistically significant. The total causal effect (TCE) had trend level significance for ornithine, alanine, nonadecenoates, triglycerides, gammaglutamyltyrosine, and hepatic function components. The TCE and NIE were both statistically significant for IL6.

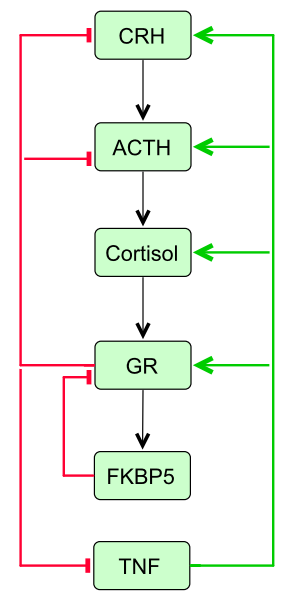

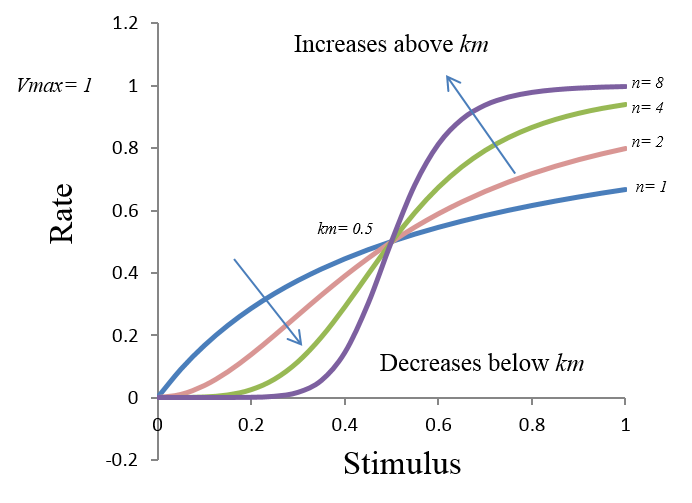

(A) (B)

Figure S7. (A) Network motif of negative feedback system in HPA-immune axis, wherein GR negatively feedbacks on HPA axis and its own synthesis through FKBP5 and downregulates cytokines (TNF). Increasing GR sensitivity of the feedback loop represented in red increases inflammatory response due to two reasons: first is the effect through the central negative feedback reduces the gain on GR synthesis due to lower ACTH stimulated cortisol release and second is the subsequent reduction in the anti-inflammatory response due to realization of increase in the inhibitory saturation threshold because of the higher sensitivity parameter that appears as the power of inhibitory threshold in the saturation kinetics. Further, the activated inflammatory response acts to further upregulate HPA axis in attempt to restore homeostasis (through the positive effect represented in green arrows), however at the cost of dysregulated immune response with increased pro-inflammatory milieu. (B) Simulated representation of change in response rates with respect to changing stimulus (*S*) are shown for different sensitivity (*n*) parameters. The Hill type kinetics as represented as: $e=Vmax*\frac{S^{n}}{S^{n}+{K_{m}}^{n}}$ . A feedback rate with respect to stimulus (*S)* is determined by three parameters (*Vmax, Km, n*), namely a maximum achievable rate, saturation threshold and sensitivity, respectively (Somvanshi and Venkatesh, 2013). It is observed that the direction of change in response rate is opposite above and below the saturation threshold (*km*), meaning on increasing the sensitivity the response is lower for the stimulus below the *km* and higher above *km*, whereas, increasing the *km* reduces the response and vice versa.

**References**

Anand, R.J., Gribar, S.C., Li, J., Kohler, J.W., Branca, M.F., Dubowski, T., Sodhi, C.P., and Hackam, D.J. (2007). Hypoxia causes an increase in phagocytosis by macrophages in a HIF‐1α‐dependent manner. Journal of Leukocyte Biology *82*, 1257-1265.

Bangsgaard, E.O., Hjorth, P.G., Olufsen, M.S., Mehlsen, J., and Ottesen, J.T. (2017). Integrated Inflammatory Stress (ITIS) Model. Bull Math Biol *79*, 1487-1509.

Bangsgaard., E.O. (2016). Mathematical Modelling of the Dynamic Role of the HPA Axis in the Immune System. Master's Thesis. In Department of Applied Mathematics and Computer Science (Technical University of Denmark ).

Bethin, K.E., Vogt, S.K., and Muglia, L.J. (2000). Interleukin-6 is an essential, corticotropin-releasing hormone-independent stimulator of the adrenal axis during immune system activation. Proceedings of the National Academy of Sciences *97*, 9317-9322.

Bhogal, R.H., Curbishley, S.M., Weston, C.J., Adams, D.H., and Afford, S.C. (2010). Reactive oxygen species mediate human hepatocyte injury during hypoxia/reoxygenation. Liver Transpl *16*, 1303-1313.

Bornstein, S.R., Rutkowski, H., and Vrezas, I. (2004). Cytokines and steroidogenesis. Mol Cell Endocrinol *215*, 135-141.

Bulik, S., Holzhütter, H.-G., and Berndt, N. (2016). The relative importance of kinetic mechanisms and variable enzyme abundances for the regulation of hepatic glucose metabolism – insights from mathematical modeling. BMC Biol *14*, 1-22.

Cassuto, H., Kochan, K., Chakravarty, K., Cohen, H., Blum, B., Olswang, Y., Hakimi, P., Xu, C., Massillon, D., Hanson, R.W.*, et al.* (2005). Glucocorticoids Regulate Transcription of the Gene for Phosphoenolpyruvate Carboxykinase in the Liver via an Extended Glucocorticoid Regulatory Unit. J Biol Chem *280*, 33873-33884.

Cavadas, M.A.S., Nguyen, L.K., and Cheong, A. (2013). Hypoxia-inducible factor (HIF) network: insights from mathematical models. Cell Commun Signal *11*, 42:41-16.

Chen, Z., Li, Y., Zhang, H., Huang, P., and Luthra, R. (2010). Hypoxia-regulated microRNA-210 modulates mitochondrial function and decreases ISCU and COX10 expression. Oncogene *29*, 4362-4368.

Copeland, S., Warren, H.S., Lowry, S.F., Calvano, S.E., and Remick, D. (2005). Acute Inflammatory Response to Endotoxin in Mice and Humans. Clin Diagn Lab Immunol *12*, 60-67.

Corcoran, S.E., and O’Neill, L.A.J. (2016). HIF1α and metabolic reprogramming in inflammation. The Journal of Clinical Investigation *126*, 3699-3707.

Cram, E.J., Ramos, R.A., Wang, E.C., Cha, H.H., Nishio, Y., and Firestone, G.L. (1998). Role of the CCAAT/Enhancer Binding Protein-α Transcription Factor in the Glucocorticoid Stimulation of p21 waf1/cip1 Gene Promoter Activity in Growth-arrested Rat Hepatoma Cells. J Biol Chem *273*, 2008-2014.

DallaMan, C., Rizza, R.A., and Cobelli, C. (2007). Meal Simulation Model of the Glucose-Insulin System. IEEE Trans Biomed Eng *54*, 1740-1749.

Duvigneau, J.C., Piskernik, C., Haindl, S., Kloesch, B., Hartl, R.T., Hüttemann, M., Lee, I., Ebel, T., Moldzio, R., Gemeiner, M.*, et al.* (2007). A novel endotoxin-induced pathway: upregulation of heme oxygenase 1, accumulation of free iron, and free iron-mediated mitochondrial dysfunction. Laboratory Investigation *88*, 70-77.

Firth, J.D., Ebert, B.L., and Ratcliffe, P.J. (1995). Hypoxic Regulation of Lactate Dehydrogenase A: Interaction between hypoxia-inducible factor 1 and cAMP response elements. J Biol Chem *270*, 21021-21027.

Frede, S., Stockmann, C., Freitag, P., and Fandrey, J. (2006). Bacterial lipopolysaccharide induces HIF-1 activation in human monocytes via p44/42 MAPK and NF-κB. Biochem J *396*, 517-527.

Frijters, R., Fleuren, W., Toonen, E.J.M., Tuckermann, J.P., Reichardt, H.M., van der Maaden, H., van Elsas, A., van Lierop, M.-J., Dokter, W., de Vlieg, J.*, et al.* (2010). Prednisolone-induced differential gene expression in mouse liver carrying wild type or a dimerization-defective glucocorticoid receptor. BMC Genomics *11*, 359:351-314.

Fukuda, R., Zhang, H., Kim, J.-w., Shimoda, L., Dang, C.V., and Semenza, Gregg L. (2007). HIF-1 Regulates Cytochrome Oxidase Subunits to Optimize Efficiency of Respiration in Hypoxic Cells. Cell *129*, 111-122.

Galgani, J., Aguirre, C., and Díaz, E. (2006). Acute effect of meal glycemic index and glycemic load on blood glucose and insulin responses in humans. Nutr J *5*, 22-22.

Gathercole, L.L., Morgan, S.A., Bujalska, I.J., Hauton, D., Stewart, P.M., and Tomlinson, J.W. (2011). Regulation of Lipogenesis by Glucocorticoids and Insulin in Human Adipose Tissue. PLoS One *6*, e26223.

Giorgino, F., Almahfouz, A., Goodyear, L.J., and Smith, R.J. (1993). Glucocorticoid regulation of insulin receptor and substrate IRS-1 tyrosine phosphorylation in rat skeletal muscle in vivo. The Journal of Clinical Investigation *91*, 2020-2030.

Gonzalez, O.R., Küper, C., Jung, K., Naval, J.P.C., and Mendoza, E. (2007). Parameter estimation using Simulated Annealing for S-system models of biochemical networks. Bioinformatics *23*, 480-486.

Grinevich, V., Ma, X.M., Herman, J.P., Jezova, D., I., A., and G., A. (2001). Effect of Repeated Lipopolysaccharide Administration on Tissue Cytokine Expression and Hypothalamic‐Pituitary‐Adrenal Axis Activity in Rats. J Neuroendocrinol *13*, 711-723.

Haggerty, D.F., Spector, E.B., Lynch, M., Kern, R., Frank, L.B., and Cederbaum, S.D. (1982). Regulation of glucocorticoids of arginase and argininosuccinate synthetase in cultured rat hepatoma cells. J Biol Chem *257*, 2246-2253.

Haigh, R.M., Jones, C.T., and Milligan, G. (1990). Glucocorticoids regulate the amount of G proteins in rat aorta. J Mol Endocrinol *5*, 185-188.

Herzig, S., Long, F., Jhala, U.S., Hedrick, S., Quinn, R., Bauer, A., Rudolph, D., Schutz, G., Yoon, C., Puigserver, P.*, et al.* (2001). CREB regulates hepatic gluconeogenesis through the coactivator PGC-1. Nature *413*, 179-183.

Hotamisligil, G.S., Peraldi, P., Budavari, A., Ellis, R., White, M.F., and Spiegelman, B.M. (1996). IRS-1-Mediated Inhibition of Insulin Receptor Tyrosine Kinase Activity in TNF-α- and Obesity-Induced Insulin Resistance. Science *271*, 665-670.

Huang, D., Li, T., Li, X., Zhang, L., Sun, L., He, X., Zhong, X., Jia, D., Song, L., Semenza, Gregg L.*, et al.* (2014). HIF-1-Mediated Suppression of Acyl-CoA Dehydrogenases and Fatty Acid Oxidation Is Critical for Cancer Progression. Cell Rep *8*, 1930-1942.

Huss, J.M., Levy, F.H., and Kelly, D.P. (2001). Hypoxia Inhibits the Peroxisome Proliferator-activated Receptor α/ Retinoid X Receptor Gene Regulatory Pathway in Cardiac Myocytes: A mechanism for O2-dependent modulation of mitochondrial fatty acid oxidation. J Biol Chem *276*, 27605-27612.

Imai, E., Miner, J.N., Mitchell, J.A., Yamamoto, K.R., and Granner, D.K. (1993). Glucocorticoid receptor-cAMP response element-binding protein interaction and the response of the phosphoenolpyruvate carboxykinase gene to glucocorticoids. J Biol Chem *268*, 5353-5356.

Jazayeri, A., and Meyer, W.J. (1988). Glucocorticoid modulation of beta-adrenergic receptors of cultured rat arterial smooth muscle cells. Hypertension *12*, 393-398.

Jeong, I.-K., Oh, S.-H., Kim, B.-J., Chung, J.-H., Min, Y.-K., Lee, M.-S., Lee, M.-K., and Kim, K.-W. (2001). The effects of dexamethasone on insulin release and biosynthesis are dependent on the dose and duration of treatment. Diabetes Res Clin Pract *51*, 163-171.

Kaiser, G., Gerst, F., Michael, D., Berchtold, S., Friedrich, B., Strutz-Seebohm, N., Lang, F., Häring, H.-U., and Ullrich, S. (2013). Regulation of forkhead box O1 (FOXO1) by protein kinase B and glucocorticoids: different mechanisms of induction of beta cell death in vitro. Diabetologia *56*, 1587-1595.

Kanczkowski, W., Alexaki, V.-I., Tran, N., Großklaus, S., Zacharowski, K., Martinez, A., Popovics, P., Block, N.L., Chavakis, T., Schally, A.V.*, et al.* (2013). Hypothalamo-pituitary and immune-dependent adrenal regulation during systemic inflammation. Proc Natl Acad Sci U S A *110*, 14801-14806.

Kariagina, A., Romanenko, D., Ren, S.-G., and Chesnokova, V. (2004). Hypothalamic-Pituitary Cytokine Network. Endocrinology *145*, 104-112.

Kastl, L., Sauer, S.W., Ruppert, T., Beissbarth, T., Becker, M.S., Süss, D., Krammer, P.H., and Gülow, K. (2014). TNF-α mediates mitochondrial uncoupling and enhances ROS-dependent cell migration via NF-κB activation in liver cells. FEBS Lett *588*, 175-183.

Kim, J., Park, M.Y., Kim, H.K., Park, Y., and Whang, K.-Y. (2016). Cortisone and dexamethasone inhibit myogenesis by modulating the AKT/mTOR signaling pathway in C2C12. Biosci Biotechnol Biochem *80*, 2093-2099.

Kim, J.W., Tchernyshyov, I., Semenza, G.L., and Dang, C.V. (2006). HIF-1-mediated expression of pyruvate dehydrogenase kinase: A metabolic switch required for cellular adaptation to hypoxia. Cell Metab *3*, 177-185.

Klover, P.J., Zimmers, T.A., Koniaris, L.G., and Mooney, R.A. (2003). Chronic Exposure to Interleukin-6 Causes Hepatic Insulin Resistance in Mice. Diabetes *52*, 2784-2789.

Koivunen, P., Hirsilä, M., Remes, A.M., Hassinen, I.E., Kivirikko, K.I., and Myllyharju, J. (2007). Inhibition of Hypoxia-inducible Factor (HIF) Hydroxylases by Citric Acid Cycle Intermediates: Possible links between cell metabolism and stabilization of HIF. J Biol Chem *282*, 4524-4532.

Kok, B.P.C., Dyck, J.R.B., Harris, T.E., and Brindley, D.N. (2013). Differential regulation of the expressions of the PGC-1α splice variants, lipins, and PPARα in heart compared to liver. J Lipid Res *54*, 1662-1677.

König, M., Bulik, S., and Holzhütter, H.-G. (2012). Quantifying the Contribution of the Liver to Glucose Homeostasis: A Detailed Kinetic Model of Human Hepatic Glucose Metabolism. PLoS Comput Biol *8*, e1002577.

Krssak, M., Brehm, A., Bernroider, E., Anderwald, C., Nowotny, P., Man, C.D., Cobelli, C., Cline, G.W., Shulman, G.I., Waldhäusl, W.*, et al.* (2004). Alterations in Postprandial Hepatic Glycogen Metabolism in Type 2 Diabetes. Diabetes *53*, 3048-3056.

Kudielka, B.M., Schommer, N.C., Hellhammer, D.H., and Kirschbaum, C. (2004). Acute HPA axis responses, heart rate, and mood changes to psychosocial stress (TSST) in humans at different times of day. Psychoneuroendocrinology *29*, 983-992.

Kuhlicke, J., Frick, J.S., Morote-Garcia, J.C., Rosenberger, P., and Eltzschig, H.K. (2007). Hypoxia Inducible Factor (HIF)-1 Coordinates Induction of Toll-Like Receptors TLR2 and TLR6 during Hypoxia. PLoS One *2*, e1364.

Kuo, T., Lew, M.J., Mayba, O., Harris, C.A., Speed, T.P., and Wang, J.-C. (2012). Genome-wide analysis of glucocorticoid receptor-binding sites in myotubes identifies gene networks modulating insulin signaling. Proceedings of the National Academy of Sciences *109*, 11160-11165.

Lambillotte, C., Gilon, P., and Henquin, J.C. (1997). Direct glucocorticoid inhibition of insulin secretion. An in vitro study of dexamethasone effects in mouse islets. J Clin Invest *99*, 414-423.

Lemke, U., Krones-Herzig, A., Diaz, M.B., Narvekar, P., Ziegler, A., Vegiopoulos, A., Cato, A.C.B., Bohl, S., Klingmüller, U., Screaton, R.A.*, et al.* (2008). The Glucocorticoid Receptor Controls Hepatic Dyslipidemia through Hes1. Cell Metab *8*, 212-223.

Leonard, M.O., Godson, C., Brady, H.R., and Taylor, C.T. (2005). Potentiation of Glucocorticoid Activity in Hypoxia through Induction of the Glucocorticoid Receptor. J Immunol *174*, 2250-2257.

Letteron, P., Brahimi-Bourouina, N., Robin, M.A., Moreau, A., Feldmann, G., and Pessayre, D. (1997). Glucocorticoids inhibit mitochondrial matrix acyl-CoA dehydrogenases and fatty acid beta-oxidation. Am J Physiol Gastrointest Liver Physiol *272*, G1141-G1150.

Li, J., Ke, W., Zhou, Q., Wu, Y., Luo, H., Zhou, H., Yang, B., Guo, Y., Zheng, Q., and Zhang, Y. (2014). Tumour necrosis factor‐α promotes liver ischaemia‐reperfusion injury through the PGC‐1α/Mfn2 pathway. J Cell Mol Med *18*, 1863-1873.

Li, L., Li, X., Zhou, W., and Messina, J.L. (2013). Acute Psychological Stress Results in the Rapid Development of Insulin Resistance. The Journal of Endocrinology *217*, 175-184.

Li, Y., Lai, N., Kirwan, J.P., and Saidel, G.M. (2012). Computational Model of Cellular Metabolic Dynamics in Skeletal Muscle Fibers during Moderate Intensity Exercise. Cell Mol Bioeng *5*, 92-112.

Li, Y., Solomon, T.P.J., Haus, J.M., Saidel, G.M., Cabrera, M.E., and Kirwan, J.P. (2010). Computational model of cellular metabolic dynamics: effect of insulin on glucose disposal in human skeletal muscle. American Journal of Physiology-Endocrinology and Metabolism *298*, E1198-E1209.

Löfberg, E., Gutierrez, A., Wernerman, J., Anderstam, B., Mitch, W.E., Price, S.R., Bergström, J., and Alvestrand, A. (2002). Effects of high doses of glucocorticoids on free amino acids, ribosomes and protein turnover in human muscle. Eur J Clin Invest *32*, 345-353.

Lu, C.-W., Lin, S.-C., Chen, K.-F., Lai, Y.-Y., and Tsai, S.-J. (2008). Induction of Pyruvate Dehydrogenase Kinase-3 by Hypoxia-inducible Factor-1 Promotes Metabolic Switch and Drug Resistance. J Biol Chem *283*, 28106-28114.

Lützner, N., Kalbacher, H., Krones-Herzig, A., and Rösl, F. (2012). FOXO3 Is a Glucocorticoid Receptor Target and Regulates LKB1 and Its Own Expression Based on Cellular AMP Levels via a Positive Autoregulatory Loop. PLoS One *7*, e42166.

MacKenzie, E.D., Selak, M.A., Tennant, D.A., Payne, L.J., Crosby, S., Frederiksen, C.M., Watson, D.G., and Gottlieb, E. (2007). Cell-Permeating α-Ketoglutarate Derivatives Alleviate Pseudohypoxia in Succinate Dehydrogenase-Deficient Cells. Mol Cell Biol *27*, 3282-3289.

Maes, M., Song, C., Lin, A., De Jongh, R., Van Gastel, A., Kenis, G., Bosmans, E., De Meester, I., Benoy, I., Neels, H.*, et al.* (1998). The effects of psychological stress on humans: Increased production of pro-inflammatory cytokines and Th-1 like response in stress induced anxiety. Cytokine *10*, 313-318.

Matsuno, F., Chowdhury, S., Gotoh, T., Iwase, K., Matsuzaki, H., Takatsuki, K., Mori, M., and Takiguchi, M. (1996). Induction of the C/EBP&beta; Gene by Dexamethasone and Glucagon in Primary-Cultured Rat Hepatocytes. The Journal of Biochemistry *119*, 524-532.

Matsuzaki, J., Kuwamura, M., Yamaji, R., Inui, H., and Nakano, Y. (2001). Inflammatory Responses to Lipopolysaccharide Are Suppressed in 40% Energy-Restricted Mice. J Nutr *131*, 2139-2144.

Nagao, K., Iwai, Y., and Miyashita, T. (2012). RCAN1 Is an Important Mediator of Glucocorticoid-Induced Apoptosis in Human Leukemic Cells. PLoS One *7*, e49926.

Nath, B., Levin, I., Csak, T., Petrasek, J., Mueller, C., Kodys, K., Catalano, D., Mandrekar, P., and Szabo, G. (2011). Hepatocyte-specific Hypoxia Inducible Factor-1α is a determinant of lipid accumulation and liver injury in alcohol-induced steatosis in mice. Hepatology *53*, 1526-1537.

Nguyen, A.T., Mandard, S., Dray, C., Deckert, V., Valet, P., Besnard, P., Drucker, D.J., Lagrost, L., and Grober, J. (2014). Lipopolysaccharides-Mediated Increase in Glucose-Stimulated Insulin Secretion: Involvement of the GLP-1 Pathway. Diabetes *63*, 471-482.

Nguyen, L.K., Cavadas, M.A.S., Scholz, C.C., Fitzpatrick, S.F., Bruning, U., Cummins, E.P., Tambuwala, M.M., Manresa, M.C., Kholodenko, B.N., Taylor, C.T.*, et al.* (2013). A dynamic model of the hypoxia-inducible factor 1α (HIF-1α) network. J Cell Sci *126*, 1454-1463.

Obach, M., Navarro-Sabaté, À., Caro, J., Kong, X., Duran, J., Gómez, M., Perales, J.C., Ventura, F., Rosa, J.L., and Bartrons, R. (2004). 6-Phosphofructo-2-kinase (pfkfb3) Gene Promoter Contains Hypoxia-inducible Factor-1 Binding Sites Necessary for Transactivation in Response to Hypoxia. J Biol Chem *279*, 53562-53570.

Okun, J.G., Conway, S., Schmidt, K.V., Schumacher, J., Wang, X., de Guia, R., Zota, A., Klement, J., Seibert, O., Peters, A.*, et al.* (2015). Molecular regulation of urea cycle function by the liver glucocorticoid receptor. Molecular Metabolism *4*, 732-740.

Parker, R., Hogg, J., Roy, A., Kellum, J., Rimmelé, T., Daun-Gruhn, S., Fedorchak, M., Valenti, I., Federspiel, W., Rubin, J.*, et al.* (2016). Modeling and Hemofiltration Treatment of Acute Inflammation. Processes *4*, 38.

Rahnert, J.A., Zheng, B., Hudson, M.B., Woodworth-Hobbs, M.E., and Price, S.R. (2016). Glucocorticoids Alter CRTC-CREB Signaling in Muscle Cells: Impact on PGC-1α Expression and Atrophy Markers. PLoS One *11*, e0159181.

Ramos, R.A., Nishio, Y., Maiyar, A.C., Simon, K.E., Ridder, C.C., Ge, Y., and Firestone, G.L. (1996). Glucocorticoid-stimulated CCAAT/enhancer-binding protein alpha expression is required for steroid-induced G1 cell cycle arrest of minimal-deviation rat hepatoma cells. Mol Cell Biol *16*, 5288-5301.

Rankin, E.B., Rha, J., Selak, M.A., Unger, T.L., Keith, B., Liu, Q., and Haase, V.H. (2009). Hypoxia-Inducible Factor 2 Regulates Hepatic Lipid Metabolism. Mol Cell Biol *29*, 4527-4538.

Rao, R., DuBois, D., Almon, R., Jusko, W.J., and Androulakis, I.P. (2016). Mathematical modeling of the circadian dynamics of the neuroendocrine-immune network in experimentally induced arthritis. Am J Physiol Endocrinol Metab *311*, E310-E324.

Saad, M.J., Folli, F., Kahn, J.A., and Kahn, C.R. (1993). Modulation of insulin receptor, insulin receptor substrate-1, and phosphatidylinositol 3-kinase in liver and muscle of dexamethasone-treated rats. J Clin Invest *92*, 2065-2072.

Samavati, L., Lee, I., Mathes, I., Lottspeich, F., and Hüttemann, M. (2008). Tumor Necrosis Factor α Inhibits Oxidative Phosphorylation through Tyrosine Phosphorylation at Subunit I of Cytochrome c Oxidase. J Biol Chem *283*, 21134-21144.

Samra, J.S., Clark, M.L., Humphreys, S.M., MacDonald, I.A., Bannister, P.A., and Frayn, K.N. (1998). Effects of Physiological Hypercortisolemia on the Regulation of Lipolysis in Subcutaneous Adipose Tissue1. The Journal of Clinical Endocrinology & Metabolism *83*, 626-631.

Selak, M.A., Armour, S.M., MacKenzie, E.D., Boulahbel, H., Watson, D.G., Mansfield, K.D., Pan, Y., Simon, M.C., Thompson, C.B., and Gottlieb, E. (2005). Succinate links TCA cycle dysfunction to oncogenesis by inhibiting HIF-α prolyl hydroxylase. Cancer Cell *7*, 77-85.

Semenza, Gregg L. (2007). Oxygen-dependent regulation of mitochondrial respiration by hypoxia-inducible factor 1. Biochem J *405*, 1-9.

Semenza, G.L., Jiang, B.-H., Leung, S.W., Passantino, R., Concordet, J.-P., Maire, P., and Giallongo, A. (1996). Hypoxia Response Elements in the Aldolase A, Enolase 1, and Lactate Dehydrogenase A Gene Promoters Contain Essential Binding Sites for Hypoxia-inducible Factor 1. J Biol Chem *271*, 32529-32537.

Semenza, G.L., Roth, P.H., Fang, H.M., and Wang, G.L. (1994). Transcriptional regulation of genes encoding glycolytic enzymes by hypoxia-inducible factor 1. J Biol Chem *269*, 23757-23763.

Shimizu, N., Yoshikawa, N., Ito, N., Maruyama, T., Suzuki, Y., Takeda, S.-i., Nakae, J., Tagata, Y., Nishitani, S., Takehana, K.*, et al.* (2011). Crosstalk between Glucocorticoid Receptor and Nutritional Sensor mTOR in Skeletal Muscle. Cell Metab *13*, 170-182.

Somvanshi, P.R., Patel, A.K., Bhartiya, S., and Venkatesh, K.V. (2016). Influence of plasma macronutrient levels on hepatic metabolism: role of regulatory networks in homeostasis and disease states. RSC Advances *6*, 14344-14371.

Somvanshi, P.R., and Venkatesh, K.V. (2013). Hill Equation. In Encyclopedia of Systems Biology, W. Dubitzky, O. Wolkenhauer, K.-H. Cho, and H. Yokota, eds. (New York, NY: Springer New York), pp. 892-895.

Son, G.H., Chung, S., Choe, H.K., Kim, H.-D., Baik, S.-M., Lee, H., Lee, H.-W., Choi, S., Sun, W., Kim, H.*, et al.* (2008). Adrenal peripheral clock controls the autonomous circadian rhythm of glucocorticoid by causing rhythmic steroid production. Proc Natl Acad Sci U S A *105*, 20970-20975.

Sriram, K., Rodriguez-Fernandez, M., and Doyle III, F.J. (2012). Modeling Cortisol Dynamics in the Neuro-endocrine Axis Distinguishes Normal, Depression, and Post-traumatic Stress Disorder (PTSD) in Humans. PLoS Comput Biol *8*, e1002379.

Sun Kim, M., Sweeney, T.R., Shigenaga, J.K., Chui, L.G., Moser, A., Grunfeld, C., and Feingold, K.R. (2007). TNF and IL-1 Decrease RXRα, PPARα, PPARγ, LXRα, and the Coactivators SRC-1, PGC-1α, and PGC-1β in Liver Cells. Metabolism *56*, 267-279.

van Uden, P., Kenneth, Niall S., and Rocha, S. (2008). Regulation of hypoxia-inducible factor-1α by NF-κB. Biochem J *412*, 477-484.

Wang, H., Kubica, N., Ellisen, L.W., Jefferson, L.S., and Kimball, S.R. (2006). Dexamethasone Represses Signaling through the Mammalian Target of Rapamycin in Muscle Cells by Enhancing Expression of REDD1. J Biol Chem *281*, 39128-39134.

Wang, R., Jiao, H., Zhao, J., Wang, X., and Lin, H. (2016). Glucocorticoids Enhance Muscle Proteolysis through a Myostatin-Dependent Pathway at the Early Stage. PLoS One *11*, e0156225.

Zhang, H., Gao, P., Fukuda, R., Kumar, G., Krishnamachary, B., Zeller, K.I., Dang, Chi V., and Semenza, G.L. (2007). HIF-1 Inhibits Mitochondrial Biogenesis and Cellular Respiration in VHL-Deficient Renal Cell Carcinoma by Repression of C-MYC Activity. Cancer Cell *11*, 407-420.

Zhao, W., Qin, W., Pan, J., Wu, Y., Bauman, W.A., and Cardozo, C. (2009). Dependence of dexamethasone-induced Akt/FOXO1 signaling, upregulation of MAFbx, and protein catabolism upon the glucocorticoid receptor. Biochem Biophys Res Commun *378*, 668-672.
